# Genomic architecture of phenotypic plasticity of complex traits in tetraploid wheat in response to water stress

**DOI:** 10.1101/565820

**Authors:** Andrii Fatiukha, Mathieu Deblieck, Valentina Klymiuk, Lianne Merchuk-Ovnat, Zvi Peleg, Frank Ordon, Tzion Fahima, Abraham B. Korol, Yehoshua Saranga, Tamar Krugman

**Affiliations:** Institute of Evolution, University of Haifa, Haifa, Israel; Department of Evolutionary and Environmental Biology, University of Haifa, Haifa, Israel; Julius Kühn-Institut (JKI) Federal Research Centre for Cultivated Plants, Institute for Resistance Research and Stress Tolerance, Quedlinburg, Germany; R. H. Smith Institute of Plant Science & Genetics in Agriculture, The Hebrew University of Jerusalem, Rehovot, Israel

**Keywords:** drought resistance strategies, flowering phenology, genomic architecture, linear regression, phenotypic plasticity, QTL analysis, wild emmer wheat

## Abstract

Phenotypic plasticity is one of the main mechanisms of adaptation to abiotic stresses via changes in critical developmental stages. Altering flowering phenology is a key evolutionary strategy of plant adaptation to abiotic stresses in order to achieve maximum possible reproduction. The current study is the first to apply the linear regression residuals as a drought plasticity scores, while taking into account the differences in flowering phenology and trait variation under non-stress conditions. We characterized the genomic architecture of 17 complex traits and their drought plasticity using a mapping population derived from a cross between durum wheat (*Triticum durum*) and wild emmer wheat (*T. dicoccoides*). We identified 79 QTLs, of which 33 were plastic in response to water stress and exhibited epistatic interactions and/or pleiotropy between the initial and plasticity traits. *Vrn-B3 (TaTF1)* residing within an interval of a major drought-escape QTL was proposed as a candidate gene. The favorable alleles for most of the plasticity QTLs were contributed by wild emmer, demonstrating the high potential of wild relatives for wheat improvement. Our study presents a new approach for quantification of plant adaptation to various stresses and provides new insights into the genetic basis of wheat complex traits under water-deficit stress.

**Highlight:** The study presents a new approach for quantification of plant adaptation to various stresses and provides new insights into the genetic basis of wheat complex traits under water-deficit stress.

## Introduction

Water stress is one of the main abiotic factors affecting plant growth and limiting crop production. Global climate changes increase the frequency of extreme drought events in many regions, thus becoming a severe threat to food security (Leng *et al*., 2015; Mann andand Gleick, 2015; Nam *et al*., 2015). Wheat is one of the most important crops worldwide, providing about 20% of calories and proteins in human consumption (FAOSTAT). Drought affects more than 42% of the worldwide wheat production area (Kosina *et al*., 2007); hence, the improvement of drought resistance in wheat cultivars is among the main targets for wheat breeders.

Crop wild relatives developed adaptation mechanisms to cope with water-limited conditions that can be used for crop improvement (Henry andand Nevo, 2014). Wild emmer wheat (WEW) (*Triticum dicoccoides*) germplasm represents an important reservoir of genetic variation for useful traits. It can increase the genetic diversity available to breeders for wheat improvement, including resistance to abiotic and biotic stresses (Levy and Feldman, 1987; Nevo *et al*., 2002; Peleg *et al*., 2005, 2009; Huang *et al*., 2016). Previously, we have evaluated WEW populations, representing an aridity gradient across Israel and vicinity, and revealed high diversity for drought stress tolerance with some genotypes displaying better performance under drought than durum wheat cultivars (Peleg *et al*., 2005). Then, we developed a mapping population derived from a cross between durum and WEW and genetically dissected drought-adaptive loci (Peleg *et al*., 2009). Subsequently, several WEW chromosomal regions conferring increased yield and drought-adaptive traits were introgressed into wheat cultivars using marker assisted selection approach (Merchuk-Ovnat *et al*., 2016a, 2016b, 2017). We also conducted whole transcriptome analyses of drought tolerant versus drought susceptible accessions of WEW in response to water stress (Krugman *et al*., 2010, 2011).

The complex responses of plants to water stress encompass multiple physiological, cellular and biochemical processes, coordinated by a large number of genes (Mickelbart *et al*., 2015). Due to the complex quantitative mode of inheritance of traits involved in response to drought stress and their effect on productivity traits, unraveling the genomic architecture of these traits is crucial for further progress in this field. Currently, the most suitable approach for genetic dissection of complex traits, such as drought resistance, is quantitative trait loci (QTL) analysis (Tardieu and Tuberosa, 2010; Blum, 2011; Lopes *et al*., 2014b; Mickelbart *et al*., 2015). QTL analysis of physiological drought adaptive traits (DAT) associated with response to drought can be used for genetic dissection of drought resistance strategies such as escape, avoidance or tolerance, as demonstrated in recent publications (Rebetzke *et al*., 2008; Adiredjo *et al*., 2014; Borrell et al., 2014; Blum, 2017). However, only a few studies were focused on the effect of physiological traits on productivity, taking into account the interaction between yield related traits and DAT (Pinto *et al*., 2010; Ahmad *et al*., 2014; Graziani *et al*., 2014; Mwadzingeni *et al*., 2016b).

Phenotypic plasticity is one of the ways of plants to respond to environmental stress; therefore, a better understanding of this phenomenon can help to improve crop management (Nicotra et al., 2010; Bloomfield *et al*., 2014). Several approaches for QTL analyses can be used for the identification of genomic regions that confer phenotypic plasticity in response to abiotic stress: (a) testing of QTL-by-environment interactions (Messmer *et al*., 2009); (b) mapping of QTLs for plasticity response (Lacaze *et al*., 2009; Adiredjo *et al*., 2014); (c) using a multi-environmental approach (MEA) in QTL analysis (van Eeuwijk *et al*., 2010); or (d) QTL mapping of a susceptibility index calculated for each trait (Peleg *et al*., 2009). Changes in flowering phenology play an important and decisive role in plant development and plasticity in response to water stress (Kamran *et al*., 2014; Riboni *et al*., 2014; Kazan and Lyons, 2016). Therefore, to reduce various biases in QTL analysis, the influence of flowering time should be taken into account when analyzing other traits. The simplest way to solve this problem is to use mapping populations with a narrow distribution of flowering time. Alternatively, the mapping population can be divided into smaller subsets of individuals by their range of flowering (Pinto *et al*., 2010). Another approach is to include various quantitative adjustments of variation in flowering time for QTL mapping of other traits (Sabadin *et al*., 2012; Hill *et al*., 2013; Lopes *et al*., 2014a; Onogi *et al*., 2016). Previously, deviations from the regression line (i.e., residuals) were defined as drought resistant indexes (DRI) for a set of pearl millet cultivars, independently of the effect of heading time and yield potential under control conditions (Bidinger *et al*., 1982). However, despite its simplicity, this approach has not been utilized in QTL mapping.

In the current study, we applied QTL mapping of phenotypic plasticity of complex traits under water-limited conditions using a recombinant inbreed line (RIL) population derived from a cross between durum wheat (*Triticum durum*) and drought resistant WEW. For QTL analysis, we targeted groups of traits related to: (a) yield; (b) phenology; (c) morphology; (d) biomass; and (e) drought-adaptive physiological traits (DAT). We employed residuals of linear regression between values of traits in control and stress conditions as drought plasticity traits. Wide distribution of heading time in the population was taken into account to reduce various biases in QTL analysis of the traits. We further used the whole genome assembly of WEW (Avni *et al*., 2017) for localization of candidate genes (CGs) associated with the studied traits, residing within the QTL intervals, including regulation of flowering and development.

## Materials and Methods

### Plant material and growth conditions

The RIL population (150 F_6_ lines) was derived from a cross between durum wheat (*T. durum*, cv. Langdon; LDN hereafter) and WEW (accession G18-16), developed by single-seed descent; hereafter referred to as G×L population (described by Peleg *et al.* 2009). The continuous water-deficient experiment was conducted in an insect-proof screen-house protected by a polyethylene top, at the experimental farm of the Hebrew University of Jerusalem in Rehovot, Israel (34°47’N, 31°54’E; 54 m above sea level). Two irrigation regimes were applied: well-watered (WW, 750 mm control) and water-limited (WL, 350 mm), irrigated with drip water system. A split-plot factorial (RIL × irrigation regime) block design with three replicates was employed; each block consisted of two main plots (for the two irrigation regimes), with main plots split into subplots as described in Peleg *et al*. (2009).

### DNA extraction and SNP genotyping

DNA was extracted from fresh leaf tissue of the parental genotypes (LDN and G18-16) and from a pooled sample of each of the 150 F_6_ RILs following a standard CTAB protocol (Doyle, 1991). DNA was normalized to 50 ng/μl. Single nucleotide polymorphism (SNP) genotyping was performed using the Illumina Infinium 15K Wheat platform, developed by TraitGenetics, Gatersleben, Germany (Muqaddasi *et al*., 2017), consisting of 12,905 SNPs selected from the wheat 90K array (Wang *et al*., 2014).

### Phenotypic traits

Four sets of phenotypic traits were used in present the QTL analysis: one set of *initial traits* and three sets of *derivative traits*. The *initial* set included 17 traits of which 13 were previously measured in the population under water-limited and well-watered conditions (described by Peleg *et al.* 2009, 2011): grain yield (GY); thousand kernel weight (TKW); kernel number per spike (KNSP); harvest index (HI); spike dry matter (SpDM); total dry matter (TotDM); carbon isotope ratio (δ13C); osmotic potential (OP); chlorophyll content (Chl); flag leaf rolling (LR); culm length (CL); days from planting to heading (DP-H); days from heading to maturity (DH-M). Four additional traits included: (i) vegetative dry matter (VegDM), comprised of stems and leaves, weighed after drying at 80°C for 48 h; (ii) spike length (SpL) (cm) measured from the base of the spike to the start of awns at maturity stage; (iii and iv) flag leaf length (FLL) (cm) and flag leaf width (FLW) (mm), of the longest and widest parts of the flag leaf, respectively. Three representative plants were measured in each plot for each trait.

Three *derivative* sets of traits were obtained by calculating the deviations from the regression line (residuals) that were then used for QTL mapping (Fig. 1). The first derivative set defined here as ‘adjusted phenology traits’ was obtained, for each environment separately, by calculating the residuals of linear regression between the means of the corresponding initial trait values and DP-H values (Fig. 1, A) in order to exclude the effect of differences in flowering phenology on these traits (prefix ‘df’ was added to the initial trait name):

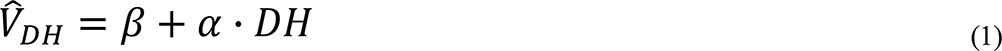

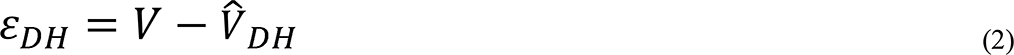

where: *V* is a value of the observed initial trait, is DP-H value, is a predicted value DH is DP-H value,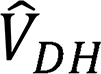a predicted value of the trait based on linear regression, is a value of the trait based on linear regression, ε_DH_ is a residual from the regression line. The regression line. The second derivative set defined here as ‘drought plasticity traits I’ was obtained by calculating the residuals of linear regression between means of the initial trait values in the WW and WL conditions (prefix ‘d’ was added to initial trait name), in order to get a deviation between trait value in WL stress and WW condition, adjusted for the differences in trait values in the population under normal conditions:

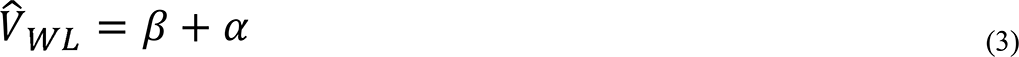

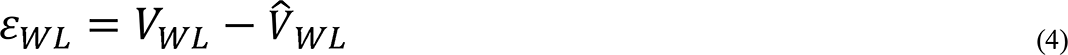

where:*V*_*WW*_ a value of the observed initial trait under WW conditions is a value of conditions *V*_*WL*_ is a value of the observed initial trait under WL conditions, is a conditions, (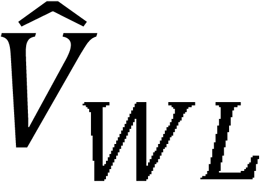) is a predicted value of the trait based on the linear regression, is a regression, ε_*WL*_ is a residual from the regression line.

**Fig. 1.**
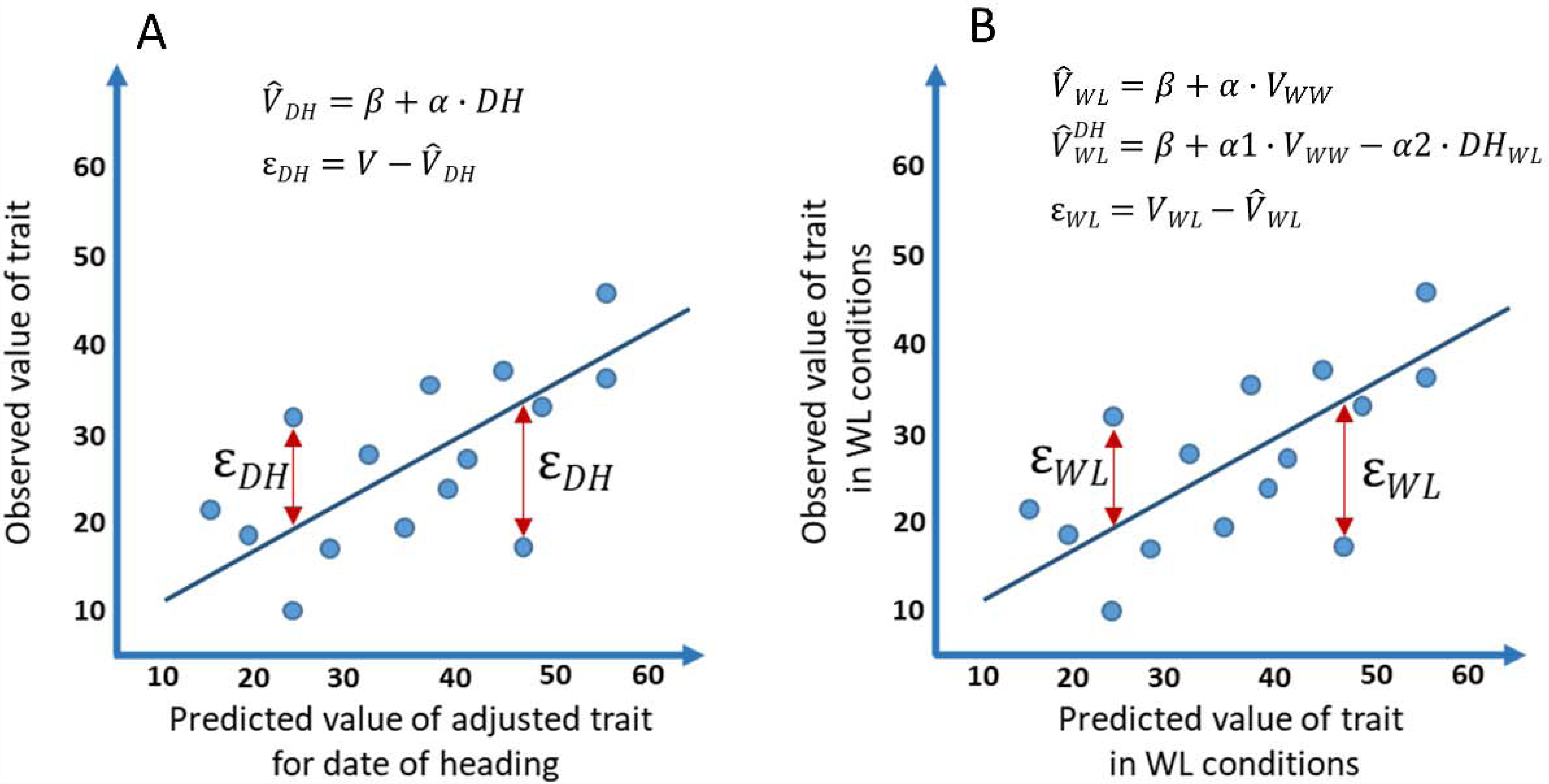
Graphical representation of the regression approach for calculation of derivative traits. A) Linear regression was calculated between means of corresponding initial traits and days from planting to heading (DH) to get predicted values of trait (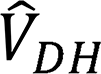) Obtained residuals between observed and predicted values of traits were used as “adjusted phenology traits” for each environment separately. B) Linear regression was calculated between means of the initial trait values in the WW and WL conditions (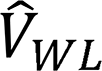) and linear regression between means of the corresponding initial trait values in the WL treatment and trait values in the WW conditions and DP-H values (DH) in the dry treatment (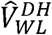) Obtained residuals were used as “drought plasticity traits I” and “drought plasticity traits II”, respectively.

The third derivative set defined here as ‘drought plasticity traits II’ was obtained by calculating the residuals of linear regression between means of the corresponding initial trait values in the WL treatment and trait values in the WW conditions and DP-H values in the WL (prefix ‘ddf’ was added to the initial trait name), in order to exclude the effect of drought escape mechanisms in ‘drought plasticity traits I’ by taking into account the effect of heading time:

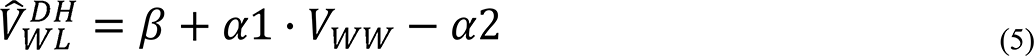

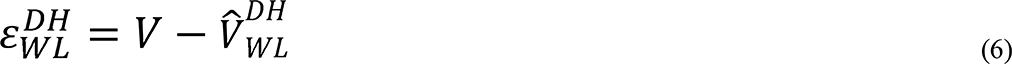

where: *V*_*WW*_ is a value of the observed initial trait under WW conditions, is a value of conditions, *V*_*WL*_ is a value of the observed initial trait under WL conditions, is a conditions, *DH*_*WL*_ is a value of DP-H under WL conditions, is a predicted value of the (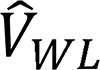) is a predicted value of the trait based on the linear regression, is a residual from the ε_*WL*_ is a residual from the regression line.

### Statistical analysis of phenotypic data

The JMP statistical package, version 11.0 (SAS Institute, Cary, NC, USA) was used for correlation and regression analyses. Correlation network analysis was conducted with the Software JASP 0.9 (JASP Team). Phenotypic values of initial and derivative traits were tested for normal distribution. The analysis of variance (ANOVA) was performed as a factorial model, with the irrigation regimes as fixed effects and genotypes and blocks as random effects. Heritability (h^2^) was calculated for each trait across the two irrigation treatments using variance components of ANOVA:

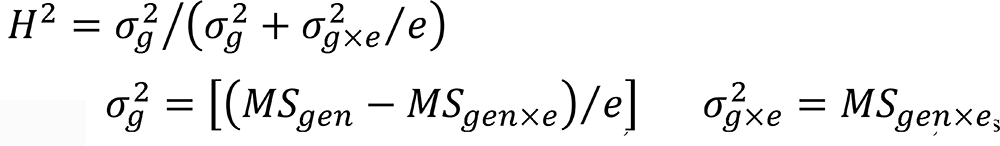

where: ***e*** is the number of environments and MS is the mean square. The correlation values and the correlation values and the descriptive statistics were calculated on the mean values of phenotypic data for each initial trait and corresponding derivative traits.

### Construction of high-density genetic map

The genetic map was constructed using *MultiPoint* software, section «UltraDense» (http://www.multiqtl.com) (Ronin *et al*., 2017). After filtering for missing data (removing markers with more than 10% missing data points) and large segregation distortion (χ^2^ > 35), the function “bound together” was applied to select the best candidate skeleton markers representing groups of co-segregating markers with size of ≥2). Clustering of candidate markers into linkage groups (LG) was performed at the threshold of recombination fraction RF=0.2. The next step included marker ordering and testing of the local map stability and monotonicity for each LG (Mester *et al*., 2003; Korol *et al*., 2009). Reducing of the final number of LGs to 14, corresponding to haploid number chromosomes of tetraploid wheat, was performed by merging the LGs with minimum pairwise RF values expressed by their end markers (end-to-end association). Orientation of each LG in relation to the short (S) and long (L) chromosome arms was performed according to the correspondence of the mapped markers with those on the consensus maps of hexaploid (Wang *et al*., 2014) and tetraploid wheat (Maccafferi *et al*., 2015).

### QTL analysis

QTL analysis was performed using the general interval mapping (IM) procedure of MultiQTL software package (http://www.multiqtl.com). First, single-QTL and two-linked-QTL models were used for screening of genetic linkage for each trait in each environment separately (Korol *et al*., 2009). Multi-environment analysis (MEA) was performed by joint analysis of trait values scored in two environments (WL and WW). After separate analysis for each chromosome, multiple interval mapping (MIM) was used for reducing the residual variation for each QTL under consideration, by taking into account QTLs that reside on other chromosomes (Kao et al., 1999). The significance of the detected QTL effects was tested using 5000 permutation runs. Significant models were further analyzed by 5000 bootstrap runs to estimate standard deviations of the chromosomal position and QTL effect. Overlapping QTL effects, when a detected QTL affects two or more separate traits, were referred to as multi-trait QTLs. The software MapChart 2.2 was used for visualization of the QTL map (Voorrips, 2002).

### Identification of physical position of the mapped SNP markers and CGs residing within QTL intervals

The physical positions of SNP markers were obtained by BLAST search of sequences of probes (Wang *et al*., 2014) against the whole-genome assembly of WEW accession ‘Zavitan’ (Avni *et al*., 2017). A list of genes residing within each QTL interval (1.5 LOD support interval of QTL effect with highest LOD) was obtained from the annotated gene models of the ‘Zavitan’ genome assembly (Avni *et al*., 2017). The putative genetic positions of the potential CGs on the QTL map were calculated based on a local linear approximation of genetic distances using the known physical positions of CGs relative to near markers.

## Results

### High-density genetic map

Genotyping of the G×L RIL population, followed by quality control, resulted in 4,347 polymorphic SNP markers. Out of these, 4,015 SNPs representing 1,369 unique loci (skeleton markers) were clustered into 14 LGs (Fig. S1). The genetic map covered 1835.7 cM (953.1 cM for the A genome and 882.6 cM for the B genome) (Table S1-S2). The number of skeletal markers and length of individual chromosome maps ranged from 51 (84.6 cM) for chr. 4B to 146 (165.3 cM) for chr. 5B. A relatively high proportion (6.3%) of non-recombinant chromosomes was observed among 150×14=2,100 RIL × chromosome combinations (Table S3). A total of 311 (22.7%) skeletal loci showed significant (P≤0.05) segregation distortion (Fig. S2), more frequently in favor of the wild rather than domesticated parent allele (203 vs. 108, respectively). The order of markers on the current genetic map showed highly similar positions on the WEW pseudomolecules (average rank correlation coefficient 0.999) (Table S4).

### Relationships between phenotypic traits

Normal distribution of most of the initial (excluding LR) and derivative quantitative traits of the RIL population was observed in each of the two environments (Fig. S3-S5, Table S5). Most of the initial and dftraits showed a wider distribution under WW that under WL conditions (Table S6). Similar range of variation in WW and WL treatments was observed for initial and dftraits of HI, FLW, LR and OP. Both phenological traits (DP-H and DH-M) and Chl exhibited a wider range under WL (Table S6). ANOVA showed highly significant effects (*P*≤0.001) of irrigation regimes for most the traits (Table S7), except SpL and FLL (0.01≤*P*≤0.05). Genotype effect was highly significant (*P*≤0.001) for most of the traits, except for VegDM and TotDM (0.001<*P*<0.01) (Table S7). The irrigation × genotype interaction was found to be significant only for DH-M and KNSP.

Correlation analysis was performed for the four groups of data: initial traits, traits adjusted for phenology, and drought plasticity traits I and II (with and without adjustment for phenology) (Tables S8-S10, Fig.2). The strongest negative correlation was observed between two initial phenological traits, DP-H and DH-M: −0.93 in WL and −0.46 in WW. Positive correlations were observed between the initial yield related traits, biomass related traits, DH-M and CL in both treatments. These traits showed negative correlation with DP-H under both conditions, with stronger correlation in the WL. This trade-off between developmental periods from planting to heading (DP-H) and from heading to maturity (DH-M) indicates strong interactions of these two phenology related traits with most of the other traits. The relationships between morphological traits and yield/biomass related traits showed varied patterns in different treatments. For example, CL and FLW showed positive correlation with VegDM in the WW (0.29 and 0.28, respectively), but no correlation in the WL. FLL and CL showed opposite directions of correlation in WW and WL (0.29 and −0.20, respectively). Most of the physiological traits were poorly correlated with traits of the other groups. All initial traits under WW were positively correlated with corresponding traits under WL (Table S11) with the lowest association for OP (r = 0.18) and the strongest association for DP-H (r = 0.85). Interestingly, variations in the traits between treatments had strong negative association with the values of traits under WW (Table S11).

The correlations of the derivative traits showed common pattern with those of the initial traits, with few exceptions (Tables S8-S10, Fig 2). Notably, no significant correlation of dDP-H with most plasticity traits was found. However, dDH-M was positively associated with productivity and yield related traits, confirming the importance of grain filling stage duration mainly in the water-limited conditions. Rank correlations (Kendall’s tau) between the initial and adjusted for heading traits (Table S12) were stronger for traits obtained under WW conditions compared to those obtained under WL conditions, suggesting that the influence of heading date to other traits was stronger in WL conditions. Relationships between the initial traits and dftraits showed common, for both WW and WL conditions, pattern. The ranks of genotypes for the dftraits were slightly different from those of the initial traits when the initial traits were uncorrelated to heading date, whereas for traits highly correlated with heading, the ranks of genotypes for dftraits have considerably changed. For example, the rank correlation between DH-M and dfDH-M in WL conditions was only 0.18, since DH-M highly correlated with heading date (−0.93). On the contrary, rank correlations between initial and drought plasticity traits were lower for more plastic to drought traits that showed lower correlations between initial traits in WW and WL conditions (Table S12).

**Fig. 2.**
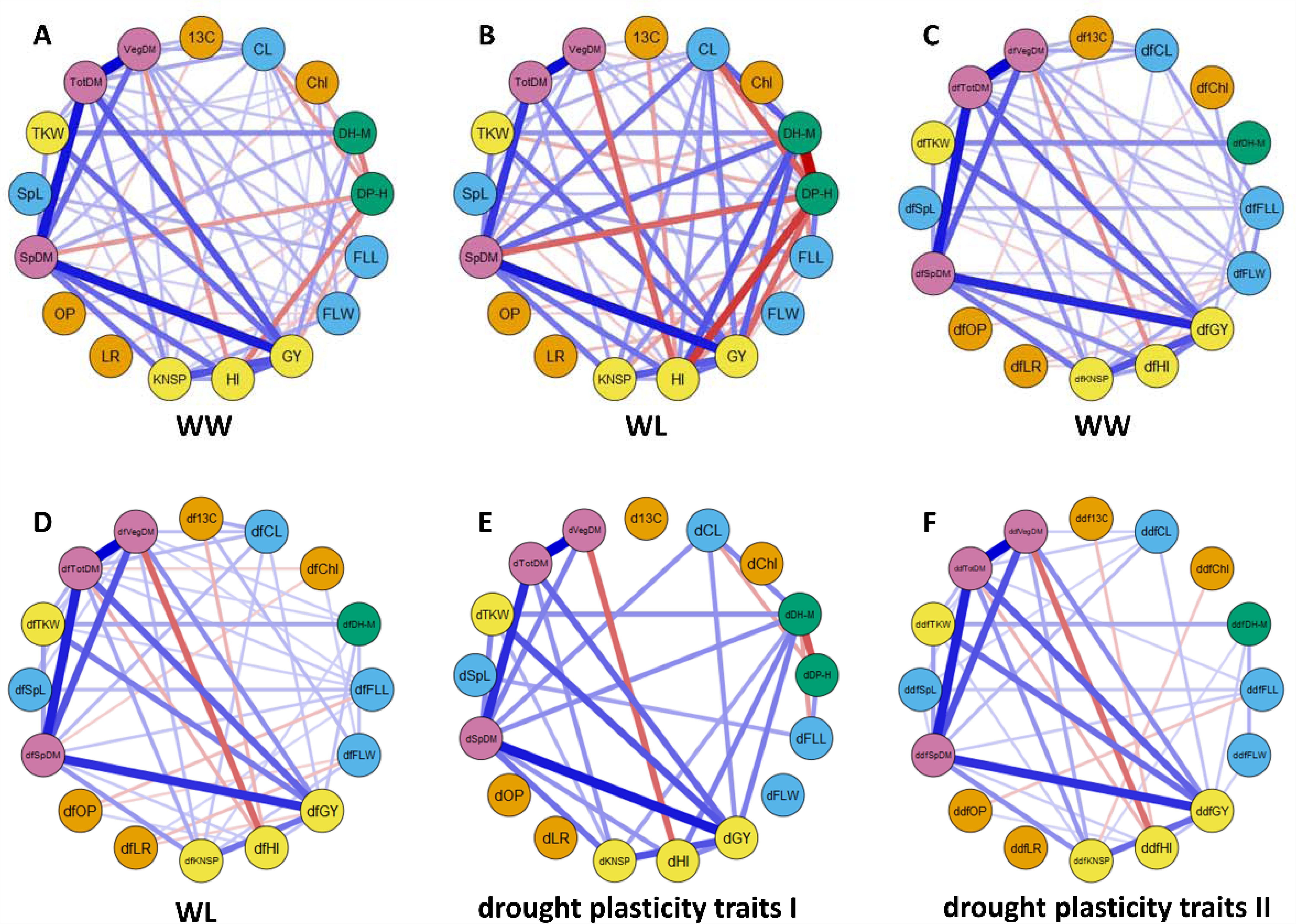
Phenotypic relationships of the analyzed traits based on correlation network analysis in 150 RILs of G×L population. The correlations between traits are shown separately for WW and for WL treatments: (A) and (B) for initial traits; (C) and (D) for traits adjusted for the effect of phenology. Correlations within group of plasticity traits without and with adjustment for the effect of phenology are presented in (E) and (F), respectively. Green hexagons are representing group of yield related traits, blue squares–group of biomass related traits, pink rhombuses –group of morphology related traits, yellow circles – drought adaptive traits and red octagons – phenology related traits. Width of lines represents the strength of correlation (minimum level of correlation is 0.16), red and blue colors correspond to positive and negative association, respectively.

### Genomic dissection of initial and derivative traits

QTL analysis was performed for 17 initial traits and 49 derived traits; among the derived traits, 16 resulted from adjustment for the effect of phenology and 33 are considered here as drought plasticity traits. In total, we detected 291 significant QTL effects distributed among 79 putative QT loci (Tables S13-S14, Figs. S6-S19), out of which 44 revealed a pleiotropic effect on two or more traits and 35 affected only one trait (Table 1, Table S14 and Fig.3). About one third of the 79 mapped loci had QTL effects only on the initial (13) or derivative (15) traits, while the majority of loci (51 out of 79) included QTL effects on both, initial and derivative traits (Tables S13-S14 and Figs. S6-S19).

**Table 1.**
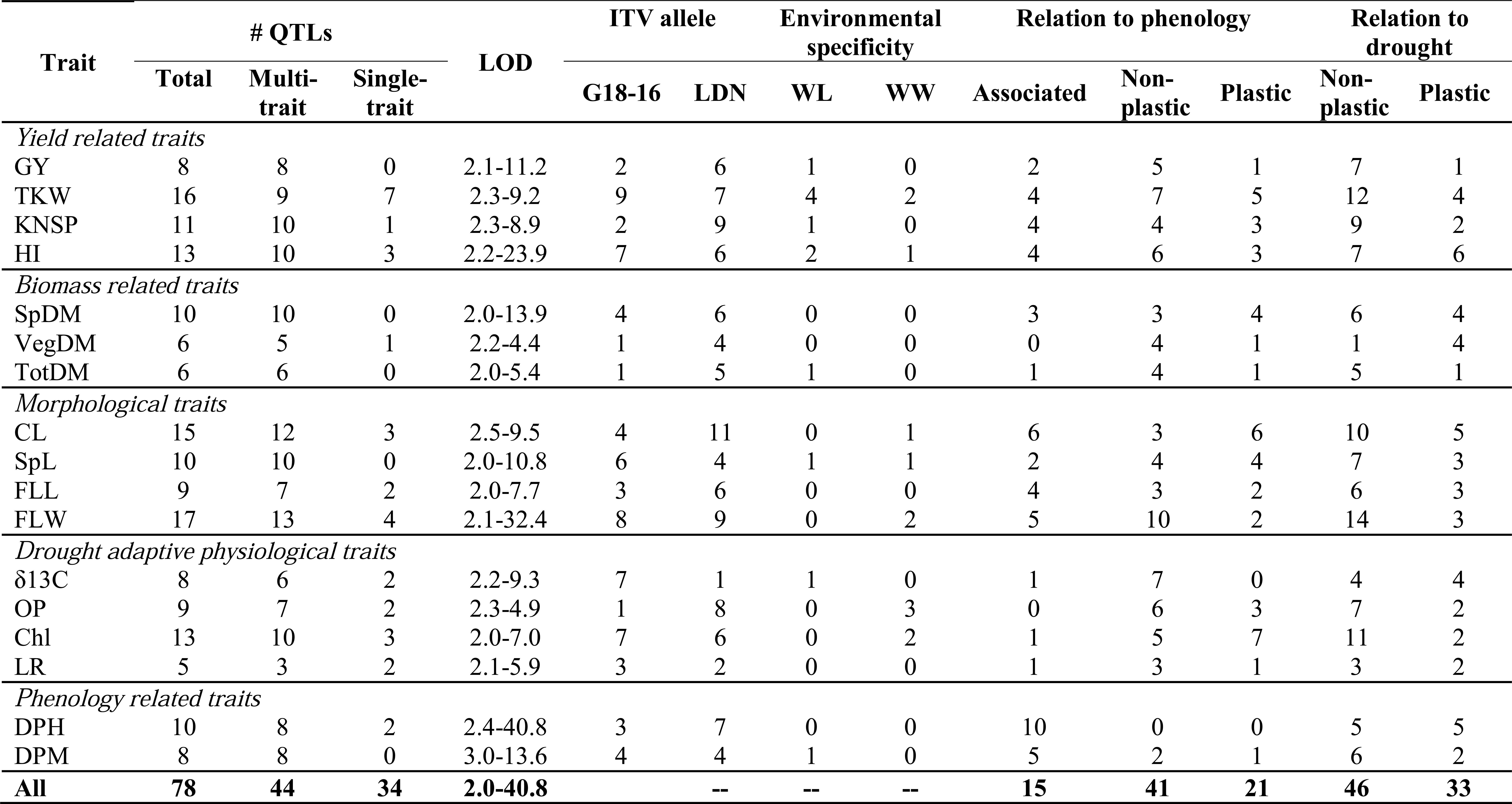
Summary of QTL effects associated with yield, biomass and phenology related, morphological, and drought adaptive physiological, traits, under water limited (WL) and well-watered (WW) conditions

### Genomic architecture of the studied traits in relation to phenology (heading time)

The presence/absence of QTL effects for derivative traits (dftraits) was used for classification of QTLs as ‘plastic’ or ‘non-plastic’ with respect to variation in phenology (Table 1, Table S14, Fig. 3). Most of the QTLs (45 out of 79) showed significant effects on both, initial and derivative traits. For 28 of these 45 QTLs, both effects were rather similar (Table S14). Therefore, we defined them as ‘non-plastic’ with respect to variation of heading date (Fig. 4). The group of QTLs classified as ‘plastic’ comprise of the following categories: (i) 17 QTLs that showed increased LOD scores and estimates of dfQTL effects after adjustment of the initial traits for variation in heading time (Fig. 4); (ii) 11 dfQTLs that affected only the derivative traits (Fig. 4); and (iii) 7 QTLs that displayed pleiotropic effects on additional traits only after correction for heading time (Table S13). Most of these 11 dfQTLs had an effect on a single trait only, excluding QTL 1A.3 that affected TKW, KNSP and OP. A total of 10 QTLs had effect on DP-H and 6 other QTLs displayed full suppression of QTL effects after adjustment for heading time. Those loci were marked as ‘associated’ with heading (Fig. 4).

**Fig. 3.**
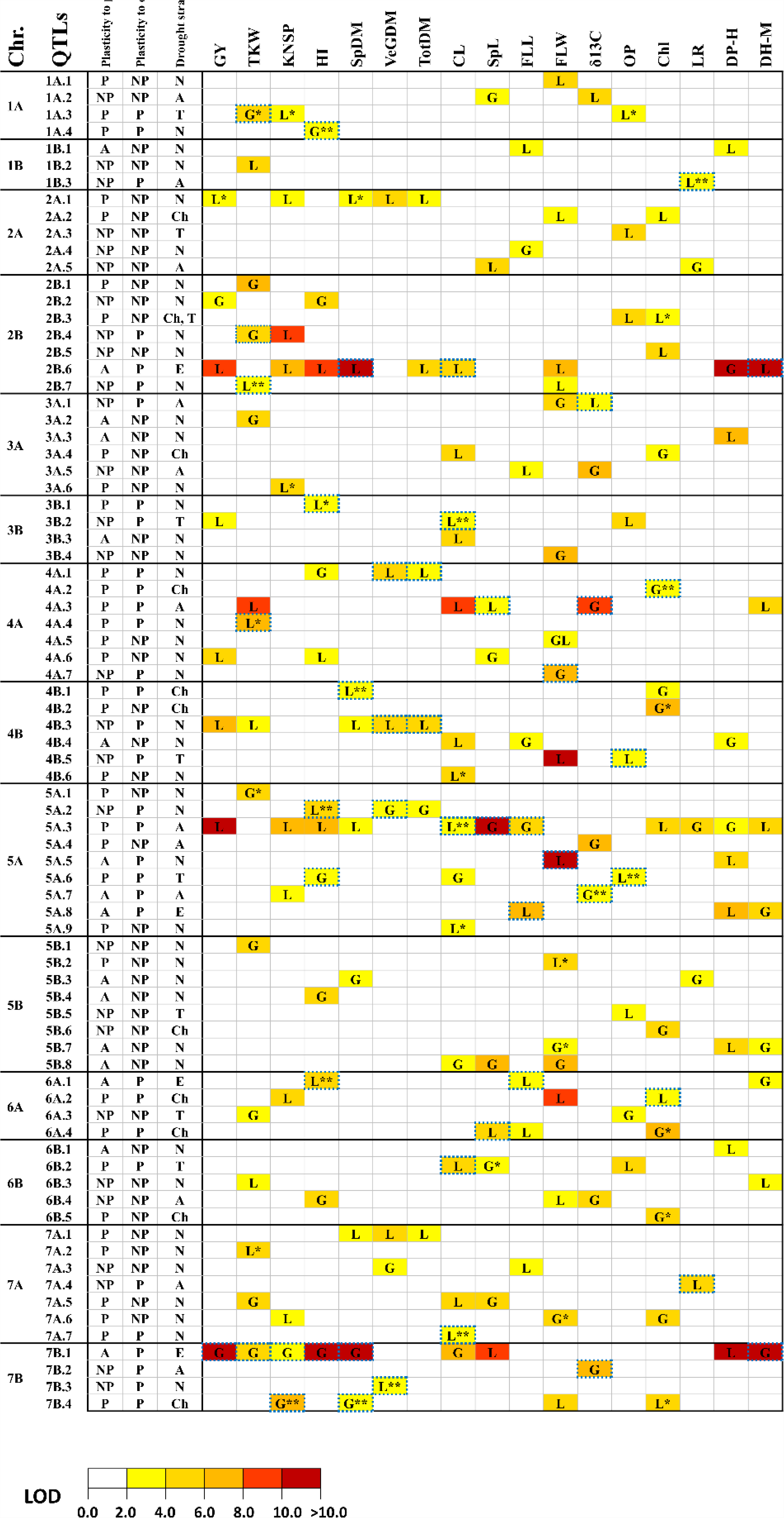
Genetic architecture of 17 traits and their relationships with phenology and plasticity to drought stress. According to our classification, QTLs were marked as follows: non-plastic (NP); plastic (P) and associated to heading (A). With respect to drought resistance strategies the QTLs were marked as: escape (E); avoidance (A); tolerance (T); chlorophyll content (Ch) and ‘no associated strategy’ (N). The origin of ITV allele is indicated as G for G18-16 and L for LDN. QTL effects only on adjusted for phenology were marked by one asterisk (*), only on plasticity traits to drought with two asterisks (**). QTLs with effects on drought plasticity traits were marked by blue dash border.

**Fig. 4.**
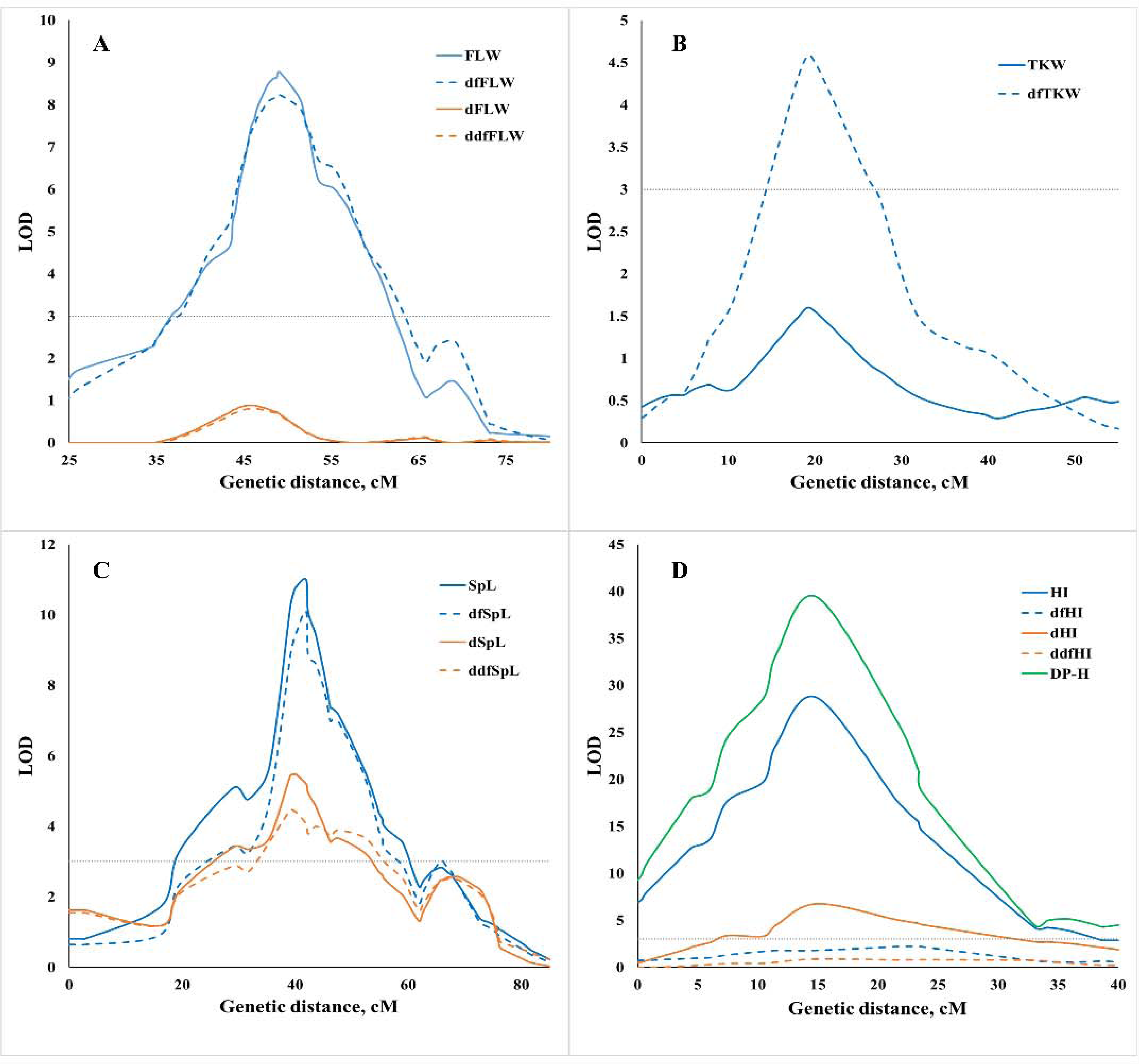
Examples of classification of detected QTLs in relation to phenology and drought plasticity: (a) non-plastic to phenology and drought QTL 6A.2; (b) plastic to phenology QTL 7A.2; (c) non-plastic to phenology and plastic to drought QTL 5A.2; (d) associated to phenology and related to drought escape strategy QTL 7B.1.

### Genomic dissection of drought plasticity traits

QTLs were defined as ‘non-plastic’ in relation to drought when significant effects were revealed for the initial traits, while no effect for drought plasticity traits (‘dtraits’ and/or ‘ddftraits’) was detected in the same QTL region. On the contrary, QTLs that affected only drought plasticity traits without displaying significant effect on the initial traits or QTLs with co-localization of effects on both initial and derivative traits were classified as ‘plastic’ (Table S14, Fig.4). According to this approach, we have identified 33 QTLs with plastic drought effects on at least one trait (Table 1, Table S14, Fig. 4): 9 QTLs for dtraits; 11 QTLs for ddftraits and 13 QTLs for combinations of dtraits and ddftraits. These results highlight the importance of adjusting for the effect of heading time in QTL mapping of plasticity to water availability and adaptation to drought. Most of plastic QTL effects (33 out 49) were collocated with corresponding initial traits, while 15 QTL effects presented 14 QTLs affected on drought plasticity traits only. A major plastic QTL for response to drought, 7B.1, affected six dtraits (dGY, dTKW, dKNSP, dHI, dSpDM, and dDH-M) with ITV allele contributed by G18-16. The highest number of plastic QTLs (six) was found for HI.

A comparison of QTL effects for plasticity to drought conferred by the G18-16 and LDN ITV alleles is presented in Table 2. In most cases, the ITV alleles for the plasticity QTL effects on the yield related traits (GY, TKW, KNSP) contributed by G18-16. For HI and SpDM, the number and PEV scores of plasticity QTL effects, were very similar for ITV alleles contributed by LDN and G18-16 (Table 2), while ITV alleles for plasticity QTL effects on VegDM and TotDM were provided mainly by LDN. Plasticity of morphological traits showed different origin of ITV alleles: equal for LDN and G18-16 for SpL; higher for LDN alleles for CL; higher for LDN for flag leaf length trait, but higher for G18-16 for flag leaf width trait. QTL analysis of plasticity of two DATs (δ13C and OP) showed that allele for higher adaptability originated from G18-16. QTLs for plasticity of Chl had equal number of ITV of G18-16 and LDN alleles. Plasticity of LR was associated with the ITV allele of LDN in the two detected QTLs, suggesting high importance of LR plasticity for plant adaptation to water deficit in LDN, which has wider leaves than G18-16.

**Table 2.** Number of QTL effects with ITV alleles of drought plasticity QTL effects, conferred by the G18-16 and LDN

| Trait | ITV allele of LDN |  | ITV allele of G18-16 |  |
| --- | --- | --- | --- | --- |
|  | Number of QTLs | Total PEV* | Number of QTLs | Total PEV* |
| GY | 0 | 0 | 1 | 0.21 |
| TKW | 1 | 0.13 | 3 | 0.37 |
| KNSP | 0 | 0 | 2 | 0.26 |
| HI | 3 | 0.33 | 3 | 0.39 |
| SpDM | 2 | 0.22 | 2 | 0.29 |
| VegDM | 3 | 0.31 | 1 | 0.13 |
| TotDM | 1 | 0.14 | 0 | 0 |
| CL | 4 | 0.46 | 1 | 0.11 |
| SpL | 2 | 0.17 | 1 | 0.17 |
| FLL | 2 | 0.21 | 1 | 0.14 |
| FLW | 1 | 0.12 | 2 | 0.21 |
| $\delta^{13}\text{C}$ | 0 | 0 | 4 | 0.53 |
| OP** | 2 | 0.30 | 0 | 0 |
| Chl | 1 | 0.09 | 1 | 0.09 |
| LR | 2 | 0.24 | 0 | 0 |
| DH-M | 1 | 0.12 | 2 | 0.32 |

\* Total PEV was calculated as a sum of PEV of effects of dtraits and ddtraits. When effects were co-located, PEV of higher effect was used for sum.
\*\* Higher value of OP was associated with susceptibility to drought, allele contributed more negative values was associated with adaptivity.

### QTLs associated with drought resistance strategies

We attempted to use QTL effects on four DATs in order to classify the QTLs in relation to drought resistance strategies, considering that OP is associated with drought tolerance strategy, δ13C and LR with avoidance strategy, and chlorophyll content as an indicator of the extent of photosynthetic apparatus damage caused by the water-deficit stress. A total of 33 QTLs fell into these categories, out of which 17 were designated as plastic (Table 3). Nevertheless, most of these QTLs (67%) did not affect yield related traits. The ITV alleles for most of QTLs affecting initial and plasticity DATs originated from G18-16 (79% and 65%, respectively). Four major plastic drought QTLs were associated with drought avoidance: three for δ13C (3A-1, 4A-3 and 7B-2 with G18-16 ITV allele) and one for LR (7A-4 with LDN ITV allele) (Table S13-S14). A major QTL effect on OP was identified on chromosome 4B (QTL 4B-5) with the G18-16 allele associated with drought tolerance (lower OP values). This QTL had also strong effect on FLW with LDN ITV allele (Table S13-S14). In the current study, we have classified QTLs as ‘drought-escape’ QTLs if they showed significant effects on drought plasticity I (dtraits), but no effect on the corresponding ddftraits for drought plasticity II, (Table 3, Fig. 4).

**Table 3.**
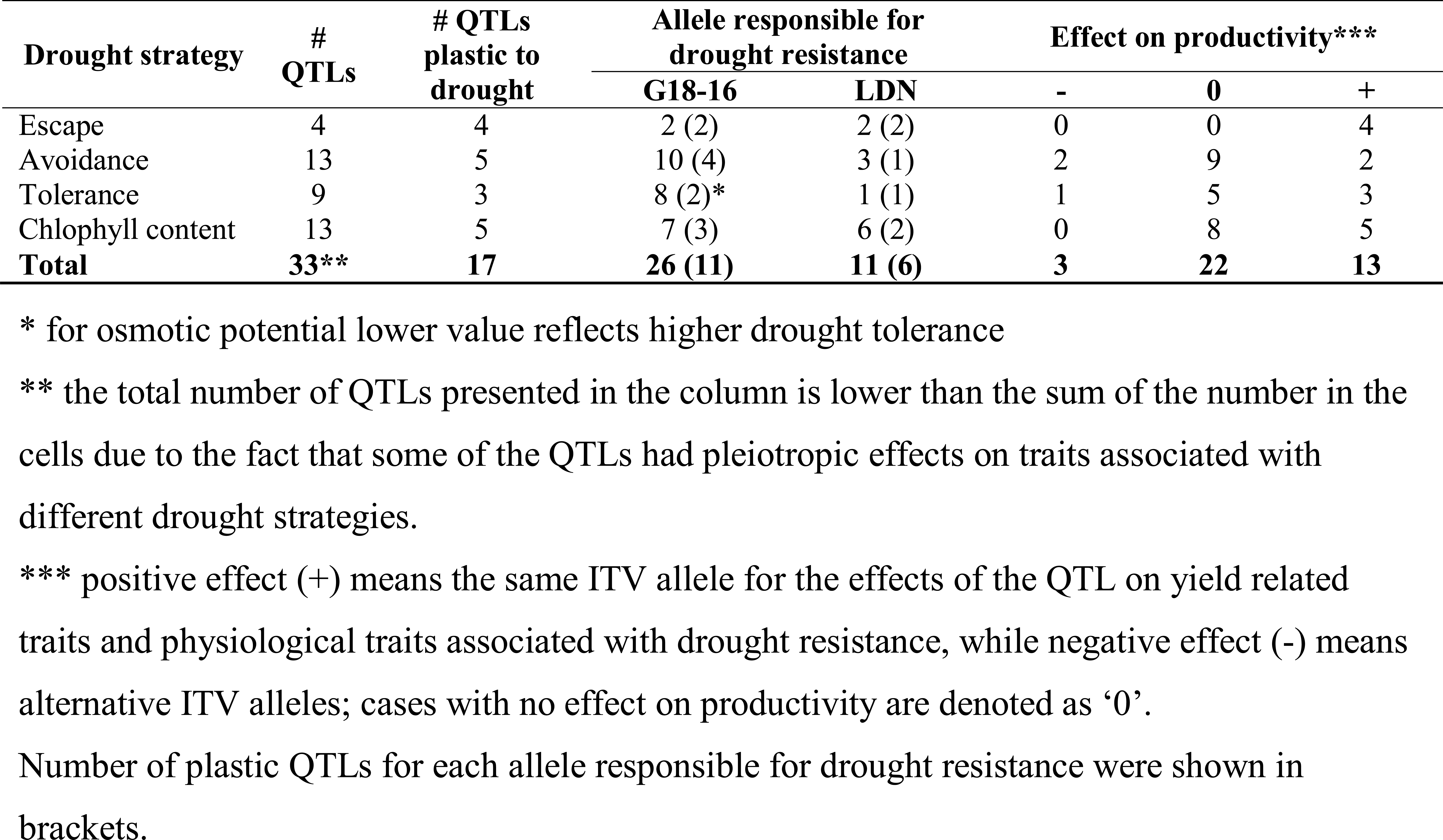
Summary of QTLs associated with drought resistance strategies and chlophyll content.

### Candidate gene analysis

More than 95% of SNPs (3824/4015) from our G×L genetic map were anchored to the reference genome of tetraploid WEW (Avni *et al*., 2017) (Table S2). The physical and genetic positions of these SNP markers (Table S2) enabled us to define the physical intervals of QTLs and the contents of genes within these intervals (Table S15). The physical intervals of QTLs ranged from 3.15 to 487.85 Mbp and the number of genes within these intervals varied from 25 to 2136 (Table S16). Most of QTLs with large intervals (>100 Mbp) were located in pericentromeric regions. On the contrary, 16 QTLs with small physical intervals (<10 Mbp) and relatively low gene content (Table S16) were dispersed along different chromosome parts, except for the pericentromeric regions. Most of QTLs with more than 200 genes within intervals were excluded from CG analysis. Our search for CGs was focused on known genes associated with studied traits, regulation of flowering and development (genes related to hormonal pathways and biosynthesis). We identified 53 potential CGs within our QTL intervals (Table S17). The putative genetic positions of CGs on the QTL map are shown in Figs. S6-S19. The list the candidates includes six CGs with well-known effect on the studied traits, *Glu*-B3 (TKW), *TaCly1* (SpL), *Wx-B1* (GY), *Wx-A1* (TKW), *WAP2-B* (SpL) and *Gpc-B1* (DH-M, TKW). A total of 11 CGs related to phenology were identified within 8 QTL intervals (Table S17). Around half of all CGs (26) were associated with regulation of hormonal balance: 13 CGs of them were related to gibberellin (GA) signaling and biosynthesis and identified within eight QTL intervals (Table S17), 11 CGs of them were associated with the ethylene signaling pathway and located within ten QTLs, and two CGs of them were associated with regulation of auxin and found within two QTLs. In addition, we identified two heat stress associated CGs (*HSFA2C* and *HSP22.0*) within two QTLs, three genes related to transport of nitrate (NRT2.6) and sugars (*STP1* and *SUT4*) within two QTLs affecting TKW, and a cluster of seven CGs with NAC domain within interval of QTL 2A.7 affecting chlorophyll content.

## Discussion

Phenotypic plasticity is one of the main mechanisms of adaptation to abiotic stresses via changes in critical developmental stages, such as the timing of transition from vegetative to reproductive growth (Kamran *et al*., 2014; Riboni *et al*., 2014). Altering flowering time is an evolutionary strategy adopted by plants to cope with environmental stresses, such as drought, in order to ensure maximum reproduction under changing environment (e.g. Kazan and Lyons, 2016). However, the genetic diversity of many crops was eroded during domestication and subsequent improvement under domestication, due to the one-sided selection for increasing of yield that reduced adaptability of cultivars (Matesanz and Milla, 2018). The present study of genomic architecture of agronomic and physiological traits plasticity in response to drought is demonstrating the effect of heading time on adaptation to water limited conditions. The comprehensive genetic analysis of the initial traits and their derivatives, based on regression residuals, enabled to identify plasticity QTLs and tentatively classify them into several drought adaptation strategies.

### Detection of QTLs using a high-density SNP-based genetic map

Several QTL mapping studies were previously conducted based on a genetic map constructed by genotyping of the G×L RIL population with SSR and DArT markers (Peleg *et al*., 2008). These studies included the genetic dissection of drought resistance (Peleg *et al*., 2009a), grain protein content (GPC) and grain micronutrient content (Peleg *et al*., 2009b), and domestication related traits (Peleg *et al*., 2011; Tzarfati *et al*., 2014). Genotyping of this RIL population with a high-throughput SNP array allowed us to achieve a shorter map (1836 cM for SNPs vs. 2317 cM of SSR-DArT map) and considerably increase the number of ordered polymorphic markers. Our current map includes over four-fold higher amount of skeletal (framework) markers (1,369 vs. 307 in previous map) and five-fold smaller average interval lengths between adjacent markers (1.3 cM vs.7.5 cM). In the current SNP map, the short arms of chromosomes 3A, 4A, 5A, 5B and 7A are extended and the short arm of chromosome 4B is fully present, while it was completely absent in the previous map. Our results confirmed the observed earlier patterns of an increased amount of non-recombinant chromosomes and segregation distortions for different chromosomes in this population (Peleg *et al*., 2008). The current SNP-based map allowed us to identify new QTLs, with improved overlap of QTL effects and shorter QTL intervals. For example, on chromosome 7AS we detected two linked QTLs affecting biomass related traits, first distal QTL (7A.1) had effects with ITV allele of LDN and second QTL (7A.3) had ITV allele of G18-16 for SpDM. The second QTL was previously detected (Peleg *et al*., 2009), the corresponding genomic region from G18-16 was introgressed into hexaploid cultivars (Merchuk-Ovnat *et al*., 2016a, 2016b, 2017) and showed an improved GY and biomass under a range of water regimes including water-deficit.

### Complexity of quantitative trait genetic architecture and the interaction with altered phenology

Correlation analysis and obtained results of QTL analysis clearly demonstrate the complexity of the studied traits and their intra- and inter-group relationships. For example, all of the detected QTLs for GY, SpDM, TotDM, SpL and DP-H traits conferred pleiotropic effects on other traits and did not show even one case of a single-trait-only QTL effect. Moreover, the average proportion of single-trait QTLs for the remaining 12 traits was also very low (~20%). Three major QTLs (2B.6 and 7B.1, and 5A.3) for phenological traits, showed strong pleiotropic effects on many other traits, with trade-off relationships between them. Furthermore, these effects were considerably stronger under WL conditions. Similar trade-off related to the influence of phenology was found in collection of Old World lupines (Berger *et al*., 2017). Strong association of phenology and other traits in genotypic (pleiotropic effects) and phenotypic (correlation) levels, together with wide range of heading dates in the studied population, required taking special measures to avoid potential biases in QTL analysis. Our results show that regression analysis used for adjustment to heading date, enabled to increase the QTL detection power and identify the relationships between the mapped QTLs and phenology.

### Drought-plasticity and drought-resistance strategies

A variance ratio and a slope of norm reaction serve as two main methods of the phenotypic plasticity quantification (Sadras and Richards, 2014). However, our approach can be applied as an alternative to these methods with two advantages: (i) normalization of drought plasticity scores for variation in the trait under non-stress conditions, (ii) accounting of the variation in additional important factors, for example phenology. Linear regression residuals were previously used for various purposes in the analysis of quantitative variation, such as the exclusion of the effect of phenology on performance of pearl millet cultivars under drought conditions (Bidinger *et al*., 1983) and characterization of the impact of heat, corrected for differences in size of the siliques between *Arabidopsis* accessions in control conditions (Bac-Molenaar *et al*., 2016). Nevertheless, it seems that the current study is the first to apply this approach for taking into account differences in phenology in order to reduce biases in mapping plasticity QTL. Tétard-Jones *et al*. (2011) used the reaction norm as a measure for plasticity traits to identify QTLs associated with barley performance in response to aphids and rhizobacteria. Although we used a different approach to map plasticity QTLs, we also found co-localization of QTL effects for the initial and the plasticity traits, as well as the presence of separate QTLs affecting only plasticity traits similar to the findings by Tétard-Jones *et al*. (2011). This phenomenon may have resulted from pleiotropic and/or epistatic effects in the genetic control of phenotypic plasticity (Scheiner, 1993). Moreover, Gulisijia and Plotkin (2017) suggested that co-located effects are the result of clustering of genes affecting phenotypic plasticity.

Three main strategies of drought resistance are recognized: drought escape, drought avoidance, and drought tolerance (Levitt, 1980) and some authors recently proposed chlorophyll content as an important drought adaptive trait (Guo and Gan, 2012; Tian *et al*., 2013; Thomas and Ougham, 2014, Borrel *et al*., 2014). Our results suggest that drought escape strategy played a central role in wheat genetic adaptation to water stress especially in the Mediterranean region (Turner, 2004).

### Candidate genes within QTL intervals

When full genome sequence is available, high-density genetic maps can provide sufficient accuracy and resolution for the identification of CGs underlying the QTLs (Thudi *et al*., 2014; Mwadzingeni et al.; 2016a, Zhang *et al*., 2017). For example, genes involved in flowering pathways in cereals were proposed as CGs for QTLs associated with complex traits (Milner *et al*., 2016). In the current study, eight genes regulating flowering time were localized within 6 out of 13 QTL intervals affecting phenology. Although, we did not identify major photoperiodic wheat *Ppd* genes within these intervals, *Ppd-A1* (Beales *et al*., 2007) was found within a QTL interval on chromosome 2A affecting all biomass related traits as well as GY and KNSP after adjustment for flowering time. Furthermore, the *TaGI*, which is known to be associated with circadian clock regulation of photoperiodic response in wheat (Zhao *et al*., 2005), was localized together with *TaFT2-A* within the 3A.2 QTL interval for DP-H. The vernalization gene *Vrn-A1* was localized within the interval of QTL 5A.5 affecting DP-H. Genes of *FT* family, which are involved in regulation of flowering, development and plant adaptation (Halliwell *et al*., 2016), were localized within 4 QTL intervals, which affected phenological traits.

Crosstalk of plant hormones is involved in plant development and its response to abiotic stresses (Peleg and Blumwald, 2011; Colenbrook *et al*., 2014). GA biosynthesis genes and signaling related genes are dispersed along several wheat chromosomes (Pearce *et al*., 2015) and some of them were proposed to be involved in response to abiotic stresses (e.g. Krugman *et al*., 2011; Colebrook *et al*., 2014; Shu *et al*., 2018). Our suggested CGs are in agreement with known function of *GA2ox* genes in response to drought (Colebrook *et al*., 2014) and our previous results that showed differential expression of *GA2ox3* in response to drought for WEW accessions (Krugman *et al*., 2011). *Ga20ox* and GASA family genes are also known to be involved in growth promotion under stress conditions (Peleg *et al*., 2011; Colebrook *et al*., 2014), while *GA13ox* genes were not reported previously as regulators of response to abiotic stresses. Ethylene response factors are also involved in regulation of plant growth (Dubois *et al*., 2018) and stress responses (Dey and Vlot, 2015). Interestingly, these ethylene CGs were identified within seven plastic to drought QTLs that highlights importance of this gene family for drought resistance in wheat.

### *Vrn-B3* is a candidate gene for major drought escape QTL

The 7B.1 QTL with the highest effect on DP-H explained 37% of the variation in flowering time. The wild parent allele of 7B.1, associated with earlier heading in both, WW and WL, prolongation of maturity period, and increasing of yield related traits in WL, appeared to have strong effects on the plasticity of yield related traits. The *Vrn-B3* (TaFT1), known to affect flowering in wheat (Yan *et al*., 2006) and found to reside within this region (together with other 25 genes), seems to be a CG for these strong effects. This gene is a homologous to the *FLOWERING LOCUS T (FT)* gene of Arabidopsis that plays a central role in control of transition from vegetative to reproductive phase in flowering plants (Lv *et al*., 2014). In barley earlier flowering was associated with increasing of *HvFT1* copy number or with haplotype differences in the promoter region and first intron of this gene (Nitcher *et al*., 2013). Orthologs of *FT* in different plant species were associated with regulation of flowering in response to abiotic stresses (Pin and Nelson, 2012; Galbiati *et al*., 2016); however, *TaFT1* was not reported as a flowering regulator in response to drought in wheat previously.

### Conclusions and future perspectives

Global climate changes require a better understanding of the genetic basis of crop plasticity in response to drought and other abiotic stresses. Here we propose a new approach for quantification of plasticity of complex traits measured under contrasting environmental conditions, which can be utilized in classical QTL and GWAS analyses of plant response to a wide range of biotic and abiotic stresses. Furthermore, application of the simultaneous adjustment of drought plasticity and phenological differences in mapping population can improve the accuracy of QTL mapping and reveal hidden during standard analysis plasticity QTLs. The identification of CGs within QTL intervals may lead to the discovery of new pleiotropic effects of these genes by their interactions with additional networks that affect not only developmental processes, but also plant response to environmental stresses. For example, *Vrn-B3 (TaFT1)*, which is proposed here as a CG underlying a major drought plasticity QTL, possibly responsible for accelerated development of plants and significant improvement of yield under WL conditions by the drought escape strategy. In addition, the higher phenotypic plasticity of the WEW parental line is confirming the importance of crop wild relatives’ genepool for improvement of crop adaptability to environmental stresses.

## Supplementary data

Table S1 Summary of the genetic map constructed based on G×L RIL population.

Table S2 Genetic and physical positions of mapped SNPs.

Table S3 Number of RILs with parental (non-recombinant) chromosomes.

Table S4 Rank correlations of genetic positions of the mapped markers with corresponding positions on other wheat genetic maps and physical positions and WEW genome assembly.

Table S5 Normality test of initial and derivative traits.

Table S6 Mean values and ranges of 17 phenotypic traits.

Table S7 Analyses of variance (Anova) and heritability (h2).

Table S8 Association between 17 initial traits in both treatments.

Table S9 Associations between adjusted traits to time of heading in both treatments.

Table S10 Association between plasticity traits to water stress with and without accounting of time of heading.

Table S11 Association between corresponding traits in WW and WL and their variation between treatments.

Table S12 Kendall’s tau coefficients of rank correlations between initial and derivative traits.

Table S13 Parameters of QTL effects.

Table S14 Summary of QTLs.

Table S15 List of HC genes within QTL intervals.

Table S16 Number of HC genes and physical intervals of QTLs.

Table S17 Summary of CGs.

Fig. S1 Distribution of attached markers along 14 wheat chromosomes.

Fig. S2 Segregation distortion for each chromosome in the G×L RIL population.

Fig. S3 Frequency distribution of the 150 F6 RILs for 17 initial phenotypic traits.

Fig. S4 Frequency distribution of the 150 F6 RILs for 16 adjusted for heading date traits.

Fig. S5 Frequency distribution of the 150 F6 RILs for 33 drought plasticity traits.

Fig. S6-S19 LOD score plot with putative genetic positions of CG and 1.5 LOD support intervals of QTL effects for 14 chromosomes.

## Acknowledgements

The research leading to these results has received funding from the European Community’s Seventh Framework Programme (FP7/ 2007-2013) under the grant agreement n°FP7-613556, Whealbi project; the Israeli Ministry of Agriculture and Rural Development, Chief Scientist Foundation (Grants 837-0079-10 and 837-0162-14); the German Federal Ministry of Food and Agriculture (FKZ: 2813IL03); the US-Israel Binational Agricultural Research and Development Fund (US-4916-16); and ISF grant for equipment (grant no. 2289/16). We thank Andy Phillips for his help with GA related genes, and Vered Barak for excellent technical assistance.

