## Supplementary Data 1 for "Genomic architecture of phenotypic plasticity of complex traits in tetraploid wheat in response to water stress"

The following Supporting Data is available for this article:

Table S1 Summary of the genetic map constructed based on G×L RIL population.

Table S2 Genetic and physical positions of mapped SNPs.

Table S3 Number of RILs with parental (non-recombinant) chromosomes.

Table S4 Rank correlations of genetic positions of the mapped markers with corresponding positions on other wheat genetic maps and physical positions and WEW genome assembly.

Table S5 Normality test of initial and derivative traits.

Table S6 Mean values and ranges of 17 phenotypic traits.

Table S7 Analyses of variance (Anova) and heritability (h2).

Table S8 Association between 17 initial traits in both treatments.

Table S9 Associations between adjusted traits to time of heading in both treatments.

Table S10 Association between plasticity traits to water stress with and without accounting of time of heading.

Table S11 Association between corresponding traits in WW and WL and their variation between treatments.

Table S12 Kendall's tau coefficients of rank correlations between initial and derivative traits.

Table S13 Parameters of QTL effects.

Table S14 Summary of QTLs.

Table S15 List of HC genes within QTL intervals.

Table S16 Number of HC genes and physical intervals of QTLs.

Table S17 Summary of CGs.

Fig. S1 Distribution of attached markers along 14 wheat chromosomes.

Fig. S2 Segregation distortion for each chromosome in the G×L RIL population.

Fig. S3 Frequency distribution of the 150 F6 RILs for 17 initial phenotypic traits.

Fig. S4 Frequency distribution of the 150 F6 RILs for 16 adjusted for heading date traits.

Fig. S5 Frequency distribution of the 150 F6 RILs for 33 drought plasticity traits.Fig. S6 LOD score plot with putative genetic positions of CG and 1.5 LOD support intervals of QTL effects for chromosome 1A.

Fig. S7 LOD score plot with putative genetic positions of CG and 1.5 LOD support intervals of QTL effects for chromosome 1B.

Fig. S8 LOD score plot with putative genetic positions of CG and 1.5 LOD support intervals of QTL effects for chromosome 2A.

Fig. S9 LOD score plot with putative genetic positions of CG and 1.5 LOD support intervals of QTL effects for chromosome 2B.

Fig. S10 LOD score plot with putative genetic positions of CG and 1.5 LOD support intervals of QTL effects for chromosome 3A.

Fig. S11 LOD score plot with putative genetic positions of CG and 1.5 LOD support intervals of QTL effects for chromosome 3B.

Fig. S12 LOD score plot with putative genetic positions of CG and 1.5 LOD support intervals of QTL effects for chromosome 4A.

Fig. S13 LOD score plot with putative genetic positions of CG and 1.5 LOD support intervals of QTL effects for chromosome 4B.

Fig. S14 LOD score plot with putative genetic positions of CG and 1.5 LOD support intervals of QTL effects for chromosome 5A.

Fig. S15 LOD score plot with putative genetic positions of CG and 1.5 LOD support intervals of QTL effects for chromosome 5B.

Fig. S16 LOD score plot with putative genetic positions of CG and 1.5 LOD support intervals of QTL effects for chromosome 6A.

Fig. S17 LOD score plot with putative genetic positions of CG and 1.5 LOD support intervals of QTL effects for chromosome 6B.

Fig. S18 LOD score plot with putative genetic positions of CG and 1.5 LOD support intervals of QTL effects for chromosome 7A.

Fig. S19 LOD score plot with putative genetic positions of CG and 1.5 LOD support intervals of QTL effects for chromosome 7B.

**Table S1** Summary of the genetic map constructed based on G×L RIL population

| Linkage group | Total markers | Skeletal markers | Co-segregated markers | Length (cM) | Average interval (cM)/locus |
|---|---|---|---|---|---|
| 1A | 267 | 89 | 178 | 123.0 | 1.38 |
| 1B | 325 | 114 | 211 | 112.9 | 0.99 |
| 2A | 242 | 90 | 152 | 159.3 | 1.77 |
| 2B | 446 | 134 | 312 | 147.0 | 1.10 |
| 3A | 256 | 84 | 172 | 138.0 | 1.64 |
| 3B | 351 | 117 | 234 | 136.9 | 1.17 |
| 4A | 184 | 72 | 112 | 127.8 | 1.78 |
| 4B | 121 | 51 | 70 | 84.6 | 1.66 |
| 5A | 300 | 100 | 200 | 157.0 | 1.57 |
| 5B | 393 | 146 | 247 | 165.3 | 1.13 |
| 6A | 278 | 85 | 193 | 107.3 | 1.26 |
| 6B | 336 | 90 | 246 | 107.5 | 1.19 |
| 7A | 257 | 96 | 161 | 140.7 | 1.47 |
| 7B | 259 | 101 | 158 | 128.4 | 1.27 |
| Group 1 | 592 | 203 | 389 | 235.9 | 1.16 |
| Group 2 | 688 | 224 | 464 | 306.3 | 1.37 |
| Group 3 | 607 | 201 | 406 | 274.9 | 1.37 |
| Group 4 | 305 | 123 | 182 | 212.4 | 1.73 |
| Group 5 | 693 | 246 | 447 | 322.3 | 1.31 |
| Group 6 | 614 | 175 | 439 | 214.8 | 1.23 |
| Group 7 | 516 | 197 | 319 | 269.1 | 1.37 |
| A genome | 1784 | 617 | 1235 | 953.1 | 1.54 |
| B genome | 2231 | 753 | 1513 | 882.6 | 1.17 |
| **Total** | **4015** | **1369** | **2646** | **1835.7** | **1.34** |

**Table S3** Number of RILs with non-recombinant chromosomes.

| Chromosome | Langdon | Gitit (G18-16) | Total | % RIL |
|---|---|---|---|---|
| 1A | 3 | 4 | 7 | 4.6 |
| 1B | 1 | 12 | 13 | 8.6 |
| 2A | 2 | 1 | 3 | 2.0 |
| 2B | 2 | 10 | 12 | 7.9 |
| 3A | 3 | 7 | 10 | 6.6 |
| 3B | 6 | 1 | 7 | 4.6 |
| 4A | 7 | 3 | 10 | 6.6 |
| 4B | 0 | 26 | 26 | 17.2 |
| 5A | 4 | 2 | 6 | 4.0 |
| 5B | 5 | 0 | 5 | 3.3 |
| 6A | 3 | 8 | 11 | 7.3 |
| 6B | 6 | 10 | 16 | 10.6 |
| 7A | 2 | 2 | 4 | 2.6 |
| 7B | 4 | 1 | 5 | 3.3 |
| Genome A | 24 | 27 | 51 | 4.8 |
| Genome B | 24 | 60 | 84 | 7.9 |
| **Total** | **48** | **87** | **135** | **6.3** |

**Table S4** Rank correlations of genetic positions of the mapped markers with corresponding positions on other wheat genetic maps and physical positions and WEW genome assembly.

| Chromosome | Svevo×Zavitan | Consensus tetraploid | Consensus hexaploid | Reference genome |
|---|---|---|---|---|
| 1A | 0.99 | 0.99 | 0.98 | 0.99 |
| 1B | 1.00 | 1.00 | 0.99 | 1.00 |
| 2A | 0.97 | 1.00 | 0.93 | 1.00 |
| 2B | 1.00 | 1.00 | 1.00 | 1.00 |
| 3A | 1.00 | 1.00 | 0.98 | 1.00 |
| 3B | 1.00 | 1.00 | 0.96 | 1.00 |
| 4A | 1.00 | 1.00 | 1.00 | 1.00 |
| 4B | 0.99 | 0.99 | 0.97 | 0.98 |
| 5A | 1.00 | 0.99 | 0.99 | 0.99 |
| 5B | 1.00 | 1.00 | 1.00 | 1.00 |
| 6A | 1.00 | 0.97 | 0.99 | 0.99 |
| 6B | 1.00 | 1.00 | 0.99 | 1.00 |
| 7A | 1.00 | 1.00 | 0.99 | 1.00 |
| 7B | 1.00 | 1.00 | 1.00 | 1.00 |
| **Average** | **1.00** | **1.00** | **0.98** | **1.00** |

**Table S6** Mean values and ranges of 17 phenotypic traits.

| | GY | VegDM | SpDM | TotDM | HI | TKW | δ13C | KNSP | DP-H | DH-M | CL | SpL | FLW | FLL | LR | Chl | OP |
|---|---|---|---|---|---|---|---|---|---|---|---|---|---|---|---|---|---|
| **Initial traits** | | | | | | | | | | | | | | | | | |
| *Mean* | | | | | | | | | | | | | | | | | |
| WL | 56.2 | 162.5 | 119.5 | 282.0 | 0.425 | 42.4 | -24.8 | 39.6 | 95.0 | 35.7 | 142.9 | 24.2 | 1.6 | 27.1 | 1.8 | 54.5 | -13.5 |
| WW | 27.8 | 111.7 | 70.8 | 182.2 | 0.389 | 30.4 | -23.0 | 32.2 | 91.3 | 32.5 | 116.8 | 23.7 | 1.4 | 20.2 | 2.1 | 59.3 | -16.1 |
| *Range* | | | | | | | | | | | | | | | | | |
| WL | 16.3–93.5 | 98.3–237.5 | 65.3–183.3 | 179.2–417.2 | 0.269–0.50 | 17.4–60.6 | -26.6–-23.5 | 14.1–69.8 | 83.0–107.0 | 27.3–41.0 | 101.0–174.2 | 19.0–47.3 | 1.1–2.4 | 18.1–35.3 | 1.0–3.0 | 47.6–64.3 | -9.7–-16.5 |
| WW | 4.9–59.3 | 70.9–167.8 | 44.7–106.7 | 134.7–254.3 | 0.298–0.523 | 14.8–45.4 | -24.0–-21.9 | 5.4–52.6 | 78.5–110.7 | 17.0–39.7 | 91.3–151.7 | 18.2–32.2 | 1.0–2.1 | 13.2–31.1 | 1.0–3.0 | 44.4–68.4 | -11.9–-19.3 |
| **Adjusted to heading date traits** | | | | | | | | | | | | | | | | | |
| *Range* | | | | | | | | | | | | | | | | | |
| WL | -27.1–28.2 | -35.4–52.1 | -28–26.7 | -45.4–73.9 | -0.077–0.076 | -14.7–13.6 | -0.98–0.86 | -19.4–20.2 | -//- | -6.16–5.66 | -36.7–30.5 | -4.8–7.5 | -0.34–0.73 | -6.10–6.20 | -1.21–1.15 | -13.5–9.2 | -3.1–4.2 |
| WW | -36.1–36.7 | -62.1–75.5 | -41.4–57.4 | -96.4–132.8 | -0.101–0.095 | -25–18.1 | -1.82–1.37 | -24.3–30.7 | -//- | -6.75–5.76 | -46.7–35.1 | -5.4–22.8 | -0.49–0.75 | -8.95–8.58 | -0.92–1.34 | -7.0–7.6 | -3.1–3.8 |
| **Drought plasticity traits I** | | | | | | | | | | | | | | | | | |
| *Range* | | | | | | | | | | | | | | | | | |
| | -24.5–25.8 | -26.3–34.7 | -26.3–34.7 | -47.1–77.2 | -0.073–0.115 | -17.1–13.7 | -1.12–1.19 | -21.3–14.8 | -7.4–11.5 | -14.93–6.29 | -25.3–29.0 | -9.0–7.2 | -0.26–0.55 | -7.40–11.12 | -1.15–1.15 | -13.1–9.7 | -2.7–3.8 |
| **Drought plasticity traits** | | | | | | | | | | | | | | | | | |
| *Range* | | | | | | | | | | | | | | | | | |
| | -30.1–22.1 | -30.9–26.7 | -30.9–26.7 | -47.6–75.8 | -0.074–0.085 | -16.2–12.8 | -0.93–0.93 | -16.6–15.5 | -//- | -5.89–5.39 | -31.3–27.3 | -8.7–6.2 | -0.30–0.53 | -6.43–6.65 | -1.22–1.24 | -12.5–10.4 | -2.6–3.9 |

**Table S7** Analyses of variance (Anova) and heritability ($h^2$) for 17 initial phenotypic traits in 150 RILs.

| | GY | VegDM | SpDM | TotDM | HI | TKW | δ13C | KNSP | DP-H | DH-M | CL | SpL | FLW | FLL | LR | Chl | OP |
|---|---|---|---|---|---|---|---|---|---|---|---|---|---|---|---|---|---|
| *Mean square* | | | | | | | | | | | | | | | | | |
| Genotype | 1898.4 *** | 1875.2 ** | 1205.4 *** | 4456.4 ** | 0.0065 *** | 130.58 *** | 0.860 *** | 305.4 *** | 159.54 *** | 48.82 *** | 738.7 *** | 28.97 *** | 0.2094 *** | 42.25 *** | 1.151 *** | 46.08 *** | 5.51 *** |
| Irrigation | 672286.3 *** | 517347.4 *** | 513293.5 *** | 2005144.5 *** | 0.27 *** | 31296.3 *** | 697.7 *** | 13418.7 *** | 2822.29 *** | 2265.7 *** | 155636.1 *** | 48.41 * | 11.88 *** | 10071.6 * | 17.37 *** | 4874.3 *** | 1377.3 *** |
| G × I | 711.6 n.s. | 1044.8 n.s. | 571.2 n.s. | 2463.6 n.s. | 0.0023 n.s. | 66.9 n.s. | 0.406 n.s. | 83.29 * | 15.15 n.s. | 25.33 *** | 186.1 n.s. | 8.50 n.s. | 0.0415 n.s. | 19.55 n.s. | 0.457 n.s. | 17.95 n.s. | 3.27 n.s. |
| Block | 1137.3 * | 2714.6 n.s. | 1117.6 n.s. | 7281.3 n.s. | 0.0051 * | 1081.2 n.s. | 10.38 *** | 22.38 n.s. | 637.3 *** | 384.7 *** | 11774.7 *** | 3.39 n.s. | 1.04 *** | 194.1 n.s. | 1.494 * | 76.93 n.s. | 19.46 *** |
| Heritability ($h^2$) | 0.63 | 0.44 | 0.53 | 0.45 | 0.65 | 0.49 | 0.53 | 0.73 | 0.91 | 0.48 | 0.75 | 0.71 | 0.80 | 0.54 | 0.60 | 0.61 | 0.41 |

*, **, *** and n.s. indicate significance at P ≤ 0.05, 0.01, 0.001 or non-significant effect, respectively.

**Table S8** Association between 17 initial traits in both treatments. Coefficients of correlation (r) between 17 initial traits in both treatments (WL and WW). Significant correlation coefficients are marked red text. Positive correlations are marked with yellow and negative are marked with blue.

| | | WL | | | | | | | | | | | | | | | | |
|---|---|---|---|---|---|---|---|---|---|---|---|---|---|---|---|---|---|---|
| | | GY | KNSp | TKW | HI | SpDM | VegDM | TotDM | CL | SpL | FLW | FLL | LR | Chl | OP | δ13C | DP-H | DH-M |
| WW | GY | 1.00 | 0.60 | 0.55 | 0.52 | 0.84 | 0.21 | 0.53 | 0.47 | -0.19 | 0.07 | -0.18 | -0.08 | 0.13 | -0.09 | -0.11 | -0.55 | 0.58 |
| | KNSP | 0.62 | 1.00 | 0.06 | 0.28 | 0.50 | 0.14 | 0.33 | 0.37 | -0.07 | 0.18 | -0.11 | -0.06 | 0.15 | -0.08 | -0.03 | -0.35 | 0.39 |
| | TKW | 0.52 | 0.19 | 1.00 | 0.25 | 0.44 | 0.13 | 0.30 | 0.29 | 0.03 | -0.03 | 0.01 | -0.01 | 0.05 | -0.08 | -0.04 | -0.30 | 0.39 |
| | HI | 0.56 | 0.33 | 0.22 | 1.00 | 0.52 | -0.55 | -0.14 | 0.39 | -0.20 | -0.09 | -0.35 | 0.09 | 0.11 | -0.12 | -0.35 | -0.71 | 0.63 |
| | SpDM | 0.82 | 0.49 | 0.35 | 0.53 | 1.00 | 0.38 | 0.73 | 0.50 | -0.01 | -0.02 | -0.11 | 0.04 | 0.04 | 0.05 | -0.17 | -0.56 | 0.54 |
| | VegDM | 0.32 | 0.23 | 0.21 | -0.38 | 0.54 | 1.00 | 0.89 | 0.05 | 0.23 | 0.06 | 0.26 | -0.03 | -0.08 | 0.18 | 0.20 | 0.24 | -0.16 |
| | TotDM | 0.61 | 0.39 | 0.30 | 0.01 | 0.83 | 0.91 | 1.00 | 0.26 | 0.19 | 0.03 | 0.14 | 0.01 | -0.04 | 0.16 | 0.04 | -0.08 | 0.14 |
| | CL | 0.17 | 0.22 | 0.10 | 0.04 | 0.25 | 0.29 | 0.31 | 1.00 | -0.05 | 0.04 | -0.20 | 0.10 | -0.03 | -0.10 | 0.11 | -0.58 | 0.58 |
| | SpL | -0.14 | -0.08 | -0.01 | -0.09 | 0.01 | 0.10 | 0.07 | 0.15 | 1.00 | 0.04 | 0.33 | 0.03 | -0.17 | 0.13 | 0.12 | 0.24 | -0.25 |
| | FLW | 0.16 | 0.32 | 0.12 | -0.07 | 0.17 | 0.28 | 0.26 | 0.16 | 0.21 | 1.00 | 0.16 | -0.24 | 0.33 | -0.25 | 0.27 | 0.07 | 0.02 |
| | FLL | -0.01 | 0.17 | 0.16 | -0.20 | 0.05 | 0.27 | 0.22 | 0.29 | 0.27 | 0.37 | 1.00 | 0.08 | -0.05 | 0.03 | 0.16 | 0.48 | -0.41 |
| | LR | -0.12 | -0.12 | -0.04 | -0.05 | 0.00 | 0.05 | 0.03 | 0.01 | 0.02 | -0.18 | -0.03 | 1.00 | -0.10 | 0.08 | -0.02 | -0.13 | 0.10 |
| | Chl | 0.27 | 0.20 | 0.23 | 0.11 | 0.23 | 0.14 | 0.20 | -0.01 | 0.04 | 0.31 | -0.04 | 0.03 | 1.00 | -0.22 | 0.01 | -0.11 | 0.13 |
| | OP | -0.24 | -0.17 | -0.11 | -0.21 | -0.22 | -0.05 | -0.13 | -0.17 | 0.08 | -0.25 | -0.07 | 0.05 | -0.24 | 1.00 | -0.14 | 0.14 | -0.16 |
| | δ13C | -0.02 | -0.09 | 0.02 | -0.11 | 0.10 | 0.25 | 0.21 | 0.26 | 0.12 | 0.12 | 0.01 | 0.11 | 0.05 | -0.07 | 1.00 | 0.22 | -0.16 |
| | DP-H | -0.31 | -0.06 | -0.06 | -0.47 | -0.37 | 0.05 | -0.16 | -0.34 | -0.06 | 0.00 | 0.09 | -0.19 | -0.21 | 0.30 | -0.07 | 1.00 | -0.93 |
| | DH-M | 0.35 | 0.07 | 0.41 | 0.21 | 0.30 | 0.10 | 0.22 | 0.26 | 0.08 | -0.04 | 0.00 | 0.13 | 0.10 | -0.07 | -0.07 | -0.46 | 1.00 |

**Table S9** Associations between adjusted traits to time of heading in both treatments. Coefficients of correlation (r) between adjusted traits to time of heading in both treatments (WW and WL). Significant correlation coefficients are marked red text. Positive correlations are marked with yellow and negative are marked with blue.

| | | Adjusted traits to time of heading (df traits) in WL | | | | | | | | | | | | | | | |
|---|---|---|---|---|---|---|---|---|---|---|---|---|---|---|---|---|---|
| | | dfGY | dfKNSp | dfTKW | dfHI | dfSpDM | dfVegDM | dfTotDM | dfCL | dfSpL | dfFLW | dfFLL | dfLR | dfChl | dfOP | dfδ13C | dfDH-M |
| Adjusted traits to time of heading (df traits) in WW | dfGY | 1.00 | 0.52 | 0.49 | 0.23 | 0.77 | 0.42 | 0.58 | 0.21 | -0.07 | 0.13 | 0.11 | -0.19 | 0.09 | -0.02 | 0.02 | 0.23 |
| | dfKNSp | 0.63 | 1.00 | -0.05 | 0.06 | 0.39 | 0.25 | 0.32 | 0.22 | 0.02 | 0.22 | 0.07 | -0.12 | 0.12 | -0.04 | 0.05 | 0.18 |
| | dfTKW | 0.53 | 0.18 | 1.00 | 0.06 | 0.35 | 0.21 | 0.29 | 0.15 | 0.11 | -0.01 | 0.18 | -0.05 | 0.02 | -0.04 | 0.03 | 0.32 |
| | dfHI | 0.49 | 0.34 | 0.21 | 1.00 | 0.20 | -0.56 | -0.29 | -0.05 | -0.04 | -0.05 | -0.01 | -0.01 | 0.05 | -0.03 | -0.28 | -0.14 |
| | dfSpDM | 0.79 | 0.51 | 0.36 | 0.44 | 1.00 | 0.63 | 0.83 | 0.26 | 0.16 | 0.02 | 0.22 | -0.04 | -0.03 | 0.15 | -0.05 | 0.09 |
| | dfVegDM | 0.36 | 0.23 | 0.22 | -0.40 | 0.60 | 1.00 | 0.94 | 0.24 | 0.19 | 0.05 | 0.17 | 0.01 | -0.06 | 0.16 | 0.15 | 0.19 |
| | dfTotDM | 0.60 | 0.39 | 0.30 | -0.07 | 0.84 | 0.93 | 1.00 | 0.26 | 0.22 | 0.04 | 0.20 | 0.00 | -0.04 | 0.17 | 0.06 | 0.18 |
| | dfCL | 0.07 | 0.21 | 0.09 | -0.14 | 0.14 | 0.32 | 0.28 | 1.00 | 0.12 | 0.10 | 0.11 | 0.03 | -0.12 | -0.03 | 0.30 | 0.11 |
| | dfSpL | -0.17 | -0.08 | -0.01 | -0.14 | -0.01 | 0.10 | 0.07 | 0.13 | 1.00 | 0.02 | 0.25 | 0.07 | -0.15 | 0.10 | 0.07 | -0.07 |
| | dfFLW | 0.17 | 0.32 | 0.12 | -0.08 | 0.19 | 0.28 | 0.27 | 0.17 | 0.21 | 1.00 | 0.15 | -0.24 | 0.34 | -0.26 | 0.26 | 0.23 |
| | dfFLL | 0.02 | 0.18 | 0.17 | -0.17 | 0.09 | 0.27 | 0.24 | 0.34 | 0.28 | 0.37 | 1.00 | 0.17 | 0.00 | -0.04 | 0.06 | 0.12 |
| | dfLR | -0.19 | -0.13 | -0.05 | -0.17 | -0.07 | 0.06 | 0.00 | -0.05 | 0.00 | -0.18 | -0.01 | 1.00 | -0.12 | 0.10 | 0.01 | -0.06 |
| | dfChl | 0.22 | 0.19 | 0.22 | 0.02 | 0.17 | 0.15 | 0.17 | -0.09 | 0.03 | 0.32 | -0.02 | -0.01 | 1.00 | -0.21 | 0.03 | 0.08 |
| | dfOP | -0.16 | -0.15 | -0.10 | -0.09 | -0.12 | -0.07 | -0.09 | -0.07 | 0.10 | -0.26 | -0.10 | 0.12 | -0.19 | 1.00 | -0.17 | -0.09 |
| | dfδ13C | -0.04 | -0.10 | 0.01 | -0.16 | 0.08 | 0.25 | 0.20 | 0.25 | 0.12 | 0.12 | 0.02 | 0.10 | 0.04 | -0.05 | 1.00 | 0.12 |
| | dfDH-M | 0.25 | 0.04 | 0.43 | -0.01 | 0.16 | 0.14 | 0.16 | 0.13 | 0.05 | -0.05 | 0.05 | 0.05 | 0.00 | 0.08 | -0.11 | 1.00 |

**Table S10** Association between plasticity traits to water stress with and without accounting of time of heading. Coefficients of correlation (r) between plasticity traits to water stress with and without accounting of time of heading. Significant correlation coefficients are marked red text. Positive correlations are marked with yellow and negative are marked with blue.

| | | Traits of plasticity to water stress with accounting of time of heading (ddftraits) | | | | | | | | | | | | | | | | |
|---|---|---|---|---|---|---|---|---|---|---|---|---|---|---|---|---|---|---|
| | | ddfGY | ddfKNSp | ddfTKW | ddfHI | ddfSpDM | ddfVegDM | ddfTotDM | ddfCL | ddfSpL | ddfFLW | ddfFLL | ddfLR | ddfChl | ddfOP | ddfδ13C | ddfDP-H | ddfDH-M |
| Traits of plasticity to water stress (dtraits) | dGY | 1.00 | 0.55 | 0.54 | 0.23 | 0.77 | 0.43 | 0.59 | 0.23 | 0.03 | 0.18 | 0.11 | -0.12 | 0.06 | 0.02 | 0.01 | -- | 0.20 |
| | dKNSp | 0.62 | 1.00 | 0.20 | 0.03 | 0.42 | 0.31 | 0.39 | 0.12 | 0.08 | 0.10 | 0.01 | -0.05 | -0.02 | 0.05 | -0.07 | -- | 0.16 |
| | dTKW | 0.60 | 0.29 | 1.00 | 0.11 | 0.35 | 0.18 | 0.26 | 0.14 | 0.11 | 0.07 | 0.14 | -0.05 | 0.06 | -0.04 | -0.03 | -- | 0.32 |
| | dHI | 0.41 | 0.22 | 0.23 | 1.00 | 0.19 | -0.53 | -0.27 | -0.01 | 0.06 | -0.02 | 0.02 | 0.04 | 0.05 | -0.05 | -0.24 | -- | -0.10 |
| | dSpDM | 0.82 | 0.51 | 0.43 | 0.38 | 1.00 | 0.65 | 0.83 | 0.26 | 0.18 | 0.08 | 0.20 | -0.02 | 0.00 | 0.15 | -0.05 | -- | 0.08 |
| | dVegDM | 0.26 | 0.21 | 0.10 | -0.54 | 0.45 | 1.00 | 0.95 | 0.18 | 0.11 | 0.06 | 0.15 | -0.01 | -0.05 | 0.18 | 0.12 | -- | 0.14 |
| | dTotDM | 0.54 | 0.38 | 0.26 | -0.20 | 0.75 | 0.91 | 1.00 | 0.21 | 0.17 | 0.08 | 0.18 | -0.01 | -0.03 | 0.18 | 0.04 | -- | 0.13 |
| | dCL | 0.38 | 0.23 | 0.24 | 0.24 | 0.39 | 0.03 | 0.19 | 1.00 | 0.17 | 0.02 | 0.10 | 0.02 | -0.02 | -0.01 | 0.13 | -- | 0.04 |
| | dSpL | -0.11 | -0.03 | 0.01 | -0.11 | 0.00 | 0.18 | 0.14 | -0.01 | 1.00 | 0.09 | 0.21 | 0.01 | -0.05 | -0.02 | 0.00 | -- | -0.04 |
| | dFLW | 0.06 | 0.04 | 0.02 | -0.12 | -0.03 | 0.09 | 0.07 | -0.07 | 0.13 | 1.00 | 0.15 | -0.10 | 0.25 | -0.10 | 0.13 | -- | 0.30 |
| | dFLL | -0.13 | -0.15 | -0.01 | -0.20 | -0.07 | 0.25 | 0.14 | -0.14 | 0.33 | 0.21 | 1.00 | 0.11 | -0.01 | -0.02 | 0.05 | -- | 0.15 |
| | dLR | -0.05 | -0.02 | -0.03 | 0.07 | 0.04 | -0.03 | -0.01 | 0.05 | -0.01 | -0.11 | 0.06 | 1.00 | -0.01 | 0.00 | 0.07 | -- | -0.03 |
| | dChl | 0.06 | -0.01 | 0.07 | 0.03 | 0.02 | -0.05 | -0.02 | 0.03 | -0.06 | 0.24 | -0.03 | 0.00 | 1.00 | -0.17 | 0.04 | -- | 0.11 |
| | dOP | 0.01 | 0.03 | -0.05 | -0.07 | 0.12 | 0.19 | 0.18 | -0.05 | -0.01 | -0.09 | 0.01 | -0.01 | -0.17 | 1.00 | -0.13 | -- | -0.10 |
| | dδ13C | -0.11 | -0.16 | -0.10 | -0.28 | -0.16 | 0.18 | 0.03 | -0.03 | 0.08 | 0.16 | 0.16 | 0.05 | 0.02 | -0.11 | 1.00 | -- | 0.10 |
| | dDP-H | -0.14 | -0.11 | -0.10 | -0.18 | -0.15 | 0.12 | 0.02 | -0.33 | 0.12 | 0.11 | 0.37 | -0.12 | -0.19 | 0.16 | 0.11 | 1.00 | -- |
| | dDH-M | 0.46 | 0.34 | 0.39 | 0.45 | 0.41 | -0.20 | 0.06 | 0.43 | -0.27 | -0.03 | -0.34 | 0.05 | 0.15 | -0.12 | -0.24 | -0.60 | 1.00 |

**Table S11.** Association between corresponding traits in WW and WL and their variation between treatments. Coefficients of correlation (R1) between values traits under WW and WL conditions and coefficients of correlation (R2) between values traits under WW and trait variation calculated as difference of values of trait under WL and WW. Significant correlation coefficients are marked red text (P≤0.05).

| | R1 | R2 |
|---|---|---|
| **GY** | 0.46 | -0.81 |
| **KNSP** | 0.67 | -0.49 |
| **TKW** | 0.29 | -0.73 |
| **HI** | 0.56 | -0.46 |
| **SpDM** | 0.35 | -0.85 |
| **VegDM** | 0.21 | -0.80 |
| **TotDM** | 0.19 | -0.83 |
| **CL** | 0.54 | -0.53 |
| **SpL** | 0.55 | -0.65 |
| **FLW** | 0.69 | -0.58 |
| **FLL** | 0.30 | -0.49 |
| **LR** | 0.41 | -0.49 |
| **Chl** | 0.41 | -0.43 |
| **OP** | 0.18 | -0.59 |
| **δ13C** | 0.31 | -0.71 |
| **DP-H** | 0.85 | 0.16 |
| **DH-M** | 0.40 | -0.14 |

**Table S12** Kendall's tau coefficients of rank correlations between initial and derivative traits.

| Traits | Adjusted to heading traits | | Drought plasticity trats I* | Drought plasticity trats II* |
|---|---|---|---|---|
| | WW | WL | | |
| **GY** | 0.797 | 0.600 | 0.679 | 0.550 |
| **KNSP** | 0.951 | 0.755 | 0.560 | 0.500 |
| **TKW** | 0.960 | 0.790 | 0.807 | 0.747 |
| **HI** | 0.680 | 0.504 | 0.625 | 0.484 |
| **SpDM** | 0.778 | 0.622 | 0.776 | 0.624 |
| **VegDM** | 0.968 | 0.853 | 0.862 | 0.802 |
| **TotDM** | 0.907 | 0.952 | 0.884 | 0.883 |
| **CL** | 0.790 | 0.609 | 0.639 | 0.519 |
| **SpL** | 0.953 | 0.851 | 0.750 | 0.691 |
| **FLW** | 0.991 | 0.964 | 0.515 | 0.502 |
| **FLL** | 0.941 | 0.701 | 0.803 | 0.634 |
| **LR** | 0.895 | 0.924 | 0.810 | 0.784 |
| **Chl** | 0.862 | 0.937 | 0.772 | 0.771 |
| **OP** | 0.807 | 0.917 | 0.840 | 0.834 |
| **δ13C** | 0.956 | 0.863 | 0.811 | 0.765 |
| **DP-H** | -//- | -//- | 0.280 | -//- |
| **DH-M** | 0.704 | 0.185 | 0.720 | 0.187 |

*For calculation of correlations for drought plasticity traits only WL initial set was used.

**Table S13** Parameters of QTL effects. Parameters of QTL effects for 17 initial traits, 16 traits adjusted to heading and 33 drought plasticity traits in G×L RIL population under two water regimes (WL – water limited and WW – well-watered).

| QTL effects | QTL | LOD | Position (cM) | Interval (cM) | Length (cM) | Nearest marker | WL | | WW | | ITV |
|---|---|---|---|---|---|---|---|---|---|---|---|
| | | | | | | | PEV | d | PEV | d | Allele |
| **Yield related traits** | | | | | | | | | | | |
| **Grain yield** | | | | | | | | | | | |
| *QdfGy.huj.uh-2A* | 2A.1 | 3.8 | 38.3 | 32.7—42.6 | 9.9 | BobWhite_c10977_834 | 0.07 | -3.85±1.40 | 0.07 | -6.53±3.75 | L |
| *QGy.huj.uh-2B.1* | 2B.2 | 2.4 | 40.8 | 29.7—50.1 | 20.3 | GENE-1741_103 | 0.13 | 3.10±1.96 | 0.15 | 4.13±3.38 | G |
| *QdfGy.huj.uh-2B.1* | 2B.2 | 2.1 | 39.4 | 29.8—52.8 | 23.0 | Kukri_c62277_80 | 0.07 | 3.07±2.56 | 0.07 | 0.47±7.75 | G |
| *QGy.huj.uh-2B.2* | 2B.6 | 8.2 | 103.8 | 101.4—107.9 | 6.6 | BobWhite_c47357_535 | 0.13 | -4.12±1.56 | 0.15 | -10.98±2.16 | L |
| *QGy.huj.uh-3B* | 3B.2 | 3.1 | 67.9 | 64.5—86.0 | 21.5 | BobWhite_c17191_297 | 0.08 | -4.91±1.86 | 0.08 | -8.00±3.17 | L |
| *QdfGy.huj.uh-3B* | 3B.2 | 2.7 | 79.3 | 65.0—87.1 | 22.1 | wsnp_Ex_c12781_20280445 | 0.08 | -3.95±1.63 | 0.08 | -7.60±3.21 | L |
| *QGy.huj.uh-4A* | 4A.6 | 4.1 | 104.0 | 96.2—110.9 | 14.7 | wsnp_Ex_c3988_7221220 | 0.05 | -3.85±1.75 | 0.12 | -9.89±3.13 | L |
| *QdfGy.huj.uh-4A* | 4A.6 | 3.1 | 104.0 | 89.1—119.5 | 30.4 | wsnp_Ex_c3988_7221220 | 0.06 | -3.66±1.47 | 0.06 | -6.53±2.94 | L |
| *QGy.huj.uh-4B* | 4B.3 | 6.1 | 35.0 | 28.7—43.0 | 14.3 | RAC875_c1052_94 | 0.07 | -4.81±1.72 | 0.11 | -10.49±2.39 | L |
| *QdfGy.huj.uh-4B* | 4B.3 | 5.2 | 23.3 | 19.5—39.7 | 20.3 | BS00022431_51 | 0.07 | -3.98±1.66 | 0.14 | -10.59±2.35 | L |
| *QGy.huj.uh-5A* | 5A.3 | 11.2 | 39.1 | 36.3—41.6 | 5.3 | Ra_c69221_1167 | 0.03 | -2.98±1.17 | 0.09 | -9.01±1.22 | L |
| *QdfGy.huj.uh-5A* | 5A.3 | 3.4 | 39.1 | 33.1—52.8 | 19.7 | Ra_c69221_1167 | 0.05 | -2.38±2.52 | 0.08 | -7.39±3.96 | L |
| *QGy.huj.uh-7B* | 7B.1 | 11.1 | 14.9 | 13.0—16.5 | 3.4 | Tdurum_contig42423_1963 | 0.23 | 8.64±0.93 | -- | -- | G |
| *QdGy.huj.uh-7B* | 7B.1 | 6.3 | 14.9 | 12.8—18.9 | 6.2 | Tdurum_contig42423_1963 | 0.21 | 7.25±1.07 | -- | -- | G |
| **Thousand kernel weight** | | | | | | | | | | | |
| *QdfTkw.huj.uh-1A* | 1A.3 | 4.7 | 94.3 | 88.9—102.6 | 13.7 | Ra_c41164_730 | 0.10 | 3.19±0.42 | -- | -- | G |
| *QddfTkw.huj.uh-1A* | 1A.3 | 2.9 | 94.3 | 87.5—110.4 | 22.9 | Ra_c41164_730 | 0.10 | 3.08±0.49 | -- | -- | G |
| *QTkw.huj.uh-1B* | 1B.2 | 5.3 | 18.0 | 10.3—22.3 | 10.0 | CAP7_c3847_204 | 0.03 | -1.63±0.96 | 0.11 | -4.46±1.09 | L |
| *QdfTkw.huj.uh-1B* | 1B.2 | 3.1 | 18.0 | 0.1—34.12 | 34.0 | GENE-1756_115 | 0.05 | -1.84±1.18 | 0.12 | -4.53±1.58 | L |
| *QTkw.huj.uh-2B.1* | 2B.1 | 3.3 | 27.4 | 19.2—36.0 | 16.8 | IAAV3165 | 0.11 | 3.48±0.42 | -- | -- | G |
| *QdfTkw.huj.uh-2B.1* | 2B.1 | 5.9 | 25.2 | 23.4—29.3 | 5.9 | RAC875_c17720_436 | 0.10 | 3.12±0.48 | -- | -- | G |
| *QddfTkw.huj.uh-2B.2* | 2B.4 | 3.7 | 83.1 | 60.5—98.2 | 37.7 | IAAV5350 | 0.12 | 3.41±0.49 | -- | -- | G |
| *QTkw.huj.uh-2B.3* | 2B.7 | 2.3 | 117.7 | 110.4—127.5 | 17.1 | Excalibur_c18966_1008 | -- | -- | 0.09 | -3.61±1.87 | L |
| *QTkw.huj.uh-3A* | 3A.2 | 5.6 | 19.9 | 15.4—24.3 | 8.9 | Excalibur_c34649_556 | 0.10 | 3.29±0.69 | 0.04 | 2.64±1.13 | G |
| *QTkw.huj.uh-4A.1* | 4A.3 | 9.2 | 44.7 | 40.6—50.2 | 9.7 | wsnp_Ex_c24443_33688235 | 0.10 | -3.24±0.72 | 0.13 | -4.77±1.03 | L |
| *QdfTkw.huj.uh-4A.2* | 4A.4 | 6.6 | 56.3 | 35.9—64.0 | 28.1 | IAAV4351 | 0.11 | -3.30±0.83 | 0.08 | -3.86±1.05 | L |
| *QddfTkw.huj.uh-4A.2* | 4A.4 | 3.2 | 60.8 | 44.9—74.6 | 29.7 | RAC875_c48107_65 | 0.13 | -3.32±1.04 | | | L |
| *QTkw.huj.uh-4B* | 4B.3 | 2.8 | 36.4 | 32.4—43.1 | 10.7 | RAC875_c25710_297 | 0.10 | -3.00±1.24 | 0.06 | -2.82±1.75 | L |
| *QdfTkw.huj.uh-4B* | 4B.3 | 3.6 | 36.4 | 33.7—39.8 | 6.1 | RAC875_c25710_297 | 0.07 | -2.45±1.24 | 0.06 | -3.03±1.40 | L |
| *QdfTkw.huj.uh-5A* | 5A.1 | 4.7 | 0.0 | 0.0—7.9 | 7.8 | BobWhite_c7114_237 | 0.05 | 1.24±1.78 | 0.09 | 3.27±2.86 | G |
| *QTkw.huj.uh-5B* | 5B.1 | 5.2 | 43.6 | 37.8—46.5 | 8.7 | BS00023081_51 | 0.06 | 2.36±0.74 | 0.09 | 4.09±0.81 | G |
| *QdfTkw.huj.uh-5B* | 5B.1 | 2.0 | 43.9 | 31.1—67.1 | 36.0 | Excalibur_c8082_478 | 0.05 | 1.83±1.28 | 0.07 | 3.23±1.68 | G |
| *QTkw.huj.uh-6A* | 6A.3 | 3.5 | 55.3 | 46.5—63.3 | 16.8 | Kukri_c14877_303 | 0.09 | 3.16±0.49 | -- | -- | G |

| QTL effects | QTL | LOD | Position (cM) | Interval (cM) | Length (cM) | Nearest marker | WL | | WW | | ITV Allele |
|---|---|---|---|---|---|---|---|---|---|---|---|
| | | | | | | | PEV | d | PEV | d | |
| *QdfTkw.huj.uh-6A* | 6A.3 | 2.4 | 53.5 | 45.0—62.3 | 17.3 | Tdurum_contig76709_195 | 0.10 | 2.93±1.21 | 0.03 | 1.68±1.87 | G |
| *QTkw.huj.uh-6B* | 6B.3 | 2.9 | 41.5 | 36.7—49.8 | 13.0 | TA002500-1001 | -- | -- | 0.08 | -2.97±2.51 | L |
| *QdfTkw.huj.uh-6B* | 6B.3 | 2.3 | 41.5 | 28.2—48.9 | 20.7 | TA002500-1001 | 0.03 | -0.79±1.61 | 0.08 | -2.49±2.90 | L |
| *QdfTkw.huj.uh-7A.1* | 7A.2 | 4.7 | 19.6 | 14.6—26.5 | 11.9 | Excalibur_rep_c115261_135 | 0.03 | -0.70±1.55 | 0.12 | -4.11±2.62 | L |
| *QTkw.huj.uh-7A.2* | 7A.5 | 4.4 | 81.8 | 73.3—96.2 | 22.9 | BS00049871_51 | 0.06 | 2.47±0.98 | 0.06 | 2.68±1.95 | G |
| *QTkw.huj.uh-7B* | 7B.1 | 5.7 | 14.91 | 11.2—31.5 | 20.35 | Tdurum_contig42423_1963 | 0.15 | 4.18±0.52 | -- | -- | G |
| *QdTkw.huj.uh-7B* | 7B.1 | 3.9 | 14.9 | 10.8—24.1 | 13.3 | Tdurum_contig42423_1963 | 0.15 | 3.84±0.62 | -- | -- | G |
| **Kernel number per spike** | | | | | | | | | | | |
| *QdfKnsp.huj.uh-1A* | 1A.3 | 3.2 | 91.7 | 86.4—101.8 | 15.4 | Excalibur_c33470_175 | 0.05 | -2.84±2.33 | 0.05 | -3.38±2.31 | L |
| *QKnsp.huj.uh-2A* | 2A.1 | 3.3 | 36.4 | 32.5—38.9 | 6.3 | wsnp_Ex_c2033_3814035 | 0.04 | -2.67±2.09 | 0.07 | -4.22±2.20 | L |
| *QdfKnsp.huj.uh-2A* | 2A.1 | 3.7 | 38.3 | 35.0—40.1 | 5.1 | BobWhite_c10977_834 | 0.04 | -2.76±1.84 | 0.06 | -3.97±2.10 | L |
| *QKnsp.huj.uh-2B.1* | 2B.4 | 8.9 | 86.0 | 82.8—88.8 | 6.0 | wsnp_Ex_c55735_58127324 | 0.20 | -0.68±7.34 | 0.23 | -1.38±8.46 | L |
| *QdfKnsp.huj.uh-2B.1* | 2B.4 | 7.5 | 86.0 | 82.8—89.3 | 6.5 | wsnp_Ex_c55735_58127324 | 0.05 | -3.35±1.20 | 0.11 | -5.94±1.02 | L |
| *QKnsp.huj.uh-2B.2* | 2B.6 | 7.3 | 106.4 | 101.7—111.6 | 9.9 | Tdurum_contig12879_1200 | 0.20 | -2.49±7.19 | 0.23 | -3.72±8.20 | L |
| *QdfKnsp.huj.uh-3A* | 3A.6 | 4.2 | 130.7 | 127.2—135.4 | 8.2 | wsnp_Ex_rep_c66274_64426901 | 0.05 | -2.85±2.28 | 0.06 | -3.55±2.80 | L |
| *QKnsp.huj.uh-5A.1* | 5A.3 | 4.7 | 45.5 | 36.9—50.7 | 13.8 | wsnp_Ex_rep_c70343_69286072 | 0.05 | -3.36±1.45 | 0.12 | -6.07±1.10 | |
| *QdfKnsp.huj.uh-5A.1* | 5A.3 | 6.9 | 45.5 | 42.7—50.2 | 7.4 | wsnp_Ex_rep_c70343_69286072 | 0.06 | -3.71±1.11 | 0.11 | -5.93±1.11 | L |
| *QKnsp.huj.uh-5A.2* | 5A.7 | 3.6 | 136.6 | 130.1—144.0 | 13.9 | wsnp_Ex_c12684_20157261 | 0.04 | -2.83±1.67 | 0.12 | -5.95±1.32 | L |
| *QKnsp.huj.uh-6A* | 6A.2 | 2.3 | 51.4 | 44.1—57.0 | 12.9 | BS00063977_51 | 0.04 | -2.02±2.47 | 0.09 | -4.78±2.38 | L |
| *QdfKnsp.huj.uh-6A* | 6A.2 | 5.1 | 48.4 | 44.33—54.32 | 10.0 | wsnp_Ex_c19770_28768859 | 0.05 | -3.35±1.07 | 0.09 | -5.36±0.97 | L |
| *QKnsp.huj.uh-7A* | 7A.6 | 3.1 | 92.2 | 79.0—98.6 | 19.6 | BobWhite_rep_c49367_405 | 0.05 | -3.31±1.47 | 0.13 | -5.96±1.23 | L |
| *QKnsp.huj.uh-7B.1* | 7B.1 | 3.0 | 14.9 | 4.7—19.8 | 15.2 | Tdurum_contig42423_1963 | 0.10 | 5.21±1.16 | -- | -- | G |
| *QdKnsp.huj.uh-7B.1* | 7B.1 | 2.8 | 14.9 | 8.0—20.9 | 12.8 | Tdurum_contig42423_1963 | 0.13 | 4.18±0.76 | -- | -- | G |
| *QdfKnsp.huj.uh-7B.2* | 7B.4 | 7.0 | 123.4 | 120.4—128.2 | 7.8 | CAP7_c3950_160 | 0.13 | 5.70±0.96 | 0.04 | 3.27±1.15 | G |
| *QddfKnsp.huj.uh-7B.2* | 7B.4 | 2.8 | 123.4 | 113.2—128.4 | 15.2 | CAP7_c3950_160 | 0.13 | 3.98±0.86 | -- | -- | G |
| **Harvest index** | | | | | | | | | | | |
| *QddfHi.huj.uh-1A* | 1A.4 | 3.8 | 109.3 | 98.4—117.4 | 19.0 | CAP12_c8163_118 | 0.10 | 0.02±0.00 | -- | -- | G |
| *QHi.huj.uh-2B.1* | 2B.2 | 2.8 | 34.0 | 29.8—38.9 | 9.0 | RAC875_rep_c109207_706 | 0.08 | 0.01±0.01 | 0.17 | 0.03±0.01 | G |
| *QdfHi.huj.uh-2B.1* | 2B.2 | 2.9 | 39.4 | 30.1—50.0 | 19.9 | Kukri_c62277_80 | 0.07 | 0.01±0.01 | 0.11 | 0.02±0.01 | G |
| *QHi.huj.uh-2B.2* | 2B.6 | 9.3 | 98.2 | 90.9—103.9 | 13.0 | Excalibur_c64983_65 | 0.08 | -0.02±0.01 | 0.17 | -0.02±0.01 | L |
| *QdfHi.huj.uh-3B* | 3B.1 | 3.5 | 31.3 | 96.8—112.5 | 15.6 | BS00022242_51 | 0.10 | -0.02±0.00 | -- | -- | L |
| *QddfHi.huj.uh-3B* | 3B.1 | 3.9 | 31.3 | 13.7—39.3 | 25.6 | BS00022242_51 | 0.10 | -0.02±0.00 | -- | -- | L |
| *QHi.huj.uh-4A.1* | 4A.1 | 2.2 | 24.0 | 4.3—31.9 | 27.7 | Kukri_rep_c101034_516 | 0.13 | 0.02±0.03 | -- | -- | G |
| *QHi.huj.uh-4A.2* | 4A.6 | 2.4 | 95.1 | 84.2—118.6 | 34.4 | BS00072157_51 | -- | -- | 0.06 | -0.01±0.02 | L |
| *QdfHi.huj.uh-4A.2* | 4A.6 | 3.1 | 94.0 | 86.0—116.5 | 30.5 | RAC875_c17197_504 | 0.09 | -0.01±0.01 | 0.04 | -0.01±0.01 | L |
| *QddfHi.huj.uh-5A.1* | 5A.2 | 4.4 | 19.5 | 16.6—33.5 | 17.0 | BS00031073_51 | 0.12 | -0.02±0.01 | -- | -- | L |
| *QHi.huj.uh-5A.2* | 5A.3 | 7.8 | 41.9 | 38.3—44.0 | 5.8 | wsnp_Ra_c14112_22155312 | 0.08 | -0.02±0.01 | 0.07 | -0.02±0.01 | L |
| *QdfHi.huj.uh-5A.2* | 5A.3 | 3.2 | 39.1 | 16.7—45.4 | 28.7 | Ra_c69221_1167 | 0.09 | -0.02±0.01 | 0.07 | -0.01±0.02 | L |
| *QdHi.huj.uh-5A.3* | 5A.6 | 3.9 | 110.9 | 97.7—138.7 | 41.0 | Ku_c21235_676 | 0.15 | 0.02±0.02 | | | G |
| *QHi.huj.uh-5B* | 5B.4 | 5.7 | 98.4 | 86.5—105.0 | 18.6 | Tdurum_contig27797_654 | 0.03 | 0.00±0.01 | 0.09 | 0.03±0.00 | G |

| QTL effects | QTL | LOD | Position (cM) | Interval (cM) | Length (cM) | Nearest marker | WL | | WW | | ITV Allele |
|---|---|---|---|---|---|---|---|---|---|---|---|
| | | | | | | | PEV | d | PEV | d | |
| *QddfHi.huj.uh-6A* | 6A.1 | 4.5 | 7.4 | 2.1—15.0 | 12.9 | Excalibur_c20597_569 | 0.11 | -0.02±0.00 | | | L |
| *QHi.huj.uh-6B* | 6B.3 | 5.8 | 51.5 | 46.6—54.3 | 7.7 | BobWhite_c1059_1825 | 0.06 | 0.02±0.00 | 0.06 | 0.02±0.01 | G |
| *QdfHi.huj.uh-6B* | 6B.3 | 5.1 | 49.7 | 44.5—55.2 | 10.7 | IACX203 | 0.10 | 0.02±0.00 | 0.07 | 0.02±0.01 | G |
| *QHi.huj.uh-7B* | 7B.1 | 23.9 | 14.9 | 13.9—15.9 | 1.9 | Tdurum_contig42423_1963 | 0.34 | 0.05±0.00 | 0.15 | 0.03±0.01 | G |
| *QdHi.huj.uh-7B* | 7B.1 | 7.0 | 14.9 | 12.7—20.9 | 8.1 | Tdurum_contig42423_1963 | 0.19 | 0.03±0.00 | | | G |
| **Biomass related traits** | | | | | | | | | | | |
| **Spike dry matter** | | | | | | | | | | | |
| *QdfSpdm.huj.uh-2A* | 2A.1 | 3.1 | 38.3 | 34.6—44.9 | 10.3 | BobWhite_c10977_834 | 0.04 | -2.50±3.09 | 0.11 | 1-0.49±8.61 | L |
| *QSpdm.huj.uh-2B* | 2B.6 | 13.9 | 103.8 | 102.2—105.7 | 3.5 | BobWhite_c47357_535 | 0.14 | -8.78±1.60 | 0.17 | -18.64±2.87 | L |
| *QdSpdm.huj.uh-2B* | 2B.6 | 2.5 | 103.8 | 95.4—118.3 | 22.9 | BobWhite_c47357_535 | 0.09 | -5.86±3.34 | -- | -- | L |
| *QdSpdm.huj.uh-4B.1* | 4B.1 | 2.0 | 8.4 | 0.0—18.0 | 18.0 | RAC875_rep_c106667_302 | 0.09 | -5.16±4.26 | -- | -- | L |
| *QddfSpdm.huj.uh-4B.1* | 4B.1 | 2.5 | 8.4 | 0.0—16.0 | 15.9 | RAC875_rep_c106667_302 | 0.13 | -6.90±1.90 | -- | -- | L |
| *QSpdm.huj.uh-4B.2* | 4B.3 | 2.8 | 40.5 | 21.9—45.4 | 23.5 | RFL_Contig3563_1130 | 0.06 | -5.16±3.11 | 0.05 | -9.02±5.97 | L |
| *QSpdm.huj.uh-5A* | 5A.3 | 3.4 | 45.5 | 33.9—51.8 | 17.9 | wsnp_Ex_rep_c70343_69286072 | 0.03 | -0.86±2.44 | 0.10 | -14.32±3.14 | L |
| *QSpdm.huj.uh-5B* | 5B.3 | 3.2 | 75.9 | 71.8—96.0 | 24.2 | Ex_c24477_484 | 0.05 | 3.49±3.72 | 0.08 | 11.12±6.28 | G |
| *QSpdm.huj.uh-7A.1* | 7A.1 | 2.4 | 7.4 | 0.0—15.3 | 15.3 | RAC875_c63822_185 | 0.07 | -3.60±8.60 | 0.08 | -11.93±16.04 | L |
| *QSpdm.huj.uh-7A.2* | 7A.3 | 1.8 | 39.4 | 32—52 | 20 | IACX17522 | 0.07 | 3.85±8.77 | 0.08 | 6.50±18.27 | G |
| *QSpdm.huj.uh-7B.1* | 7B.1 | 10.6 | 14.9 | 5.0—19.1 | 14.1 | Tdurum_contig42423_1963 | 0.18 | 9.85±1.86 | 0.10 | 13.70±4.21 | G |
| *QdSpdm.huj.uh-7B.1* | 7B.1 | 6.5 | 14.9 | 13.2—20.7 | 7.5 | Tdurum_contig42423_1963 | 0.19 | 9.53±1.33 | | | G |
| *QddfSpdm.huj.uh-7B.2* | 7B.4 | 2.3 | 128.4 | 119.4—128.4 | 9.0 | RFL_Contig3607_648 | 0.10 | 5.48±3.04 | | | G |
| **Vegetative dry matter** | | | | | | | | | | | |
| *QVegdm.huj.uh-2A* | 2A.1 | 3.7 | 36.4 | 33.5—41.2 | 7.7 | wsnp_Ex_c2033_3814035 | 0.06 | -7.99±5.45 | 0.08 | -14.48±8.46 | L |
| *QdfVegdm.huj.uh-2A* | 2A.1 | 4.4 | 38.3 | 35.7—42.5 | 6.8 | BobWhite_c10977_834 | 0.06 | -8.12±3.81 | 0.07 | -13.24±7.53 | L |
| *QVegdm.huj.uh-4A* | 4A.1 | 4.1 | 17.1 | 7.2—19.6 | 12.4 | BS00065863_51 | 0.13 | -12.42±6.56 | 0.03 | -5.79±7.42 | L |
| *QdfVegdm.huj.uh-4A* | 4A.1 | 3.6 | 17.1 | 7.7—19.5 | 11.8 | BS00065863_51 | 0.11 | -10.99±5.77 | 0.03 | -6.70±6.86 | L |
| *QdVegdm.huj.uh-4A* | 4A.1 | 3.7 | 17.1 | 7.2—19.8 | 12.5 | BS00065863_51 | 0.12 | -12.23±4.98 | -- | -- | L |
| *QddfVegdm.huj.uh-4A* | 4A.1 | 2.4 | 17.1 | 4.0—33.7 | 29.7 | BS00065863_51 | 0.10 | -9.48±6.43 | -- | -- | L |
| *QVegdm.huj.uh-4B* | 4B.3 | 3.1 | 40.5 | 32.2—45.0 | 12.9 | RFL_Contig3563_1130 | 0.08 | -8.98±5.83 | 0.09 | -8.43±15.27 | L |
| *QdfVegdm.huj.uh-4B* | 4B.3 | 4.1 | 40.5 | 33.9—44.8 | 10.9 | RFL_Contig3563_1130 | 0.09 | -10.38±4.53 | 0.08 | -9.84±13.08 | L |
| *QddfVegdm.huj.uh-4B* | 4B.3 | 2.3 | 32.2 | 29.2—44.8 | 15.6 | TA002925-3757 | 0.11 | -11.71±2.67 | -- | -- | L |
| *QVegdm.huj.uh-5A* | 5A.2 | 2.6 | 29.1 | 11.9—46.0 | 34.1 | wsnp_CAP7_c2282_1107112 | 0.05 | 10.03±7.02 | 0.03 | 3.79±7.44 | G |
| *QdfVegdm.huj.uh-5A* | 5A.2 | 3.0 | 29.1 | 14.2—45.4 | 31.2 | wsnp_CAP7_c2282_1107112 | 0.11 | 11.48±4.02 | 0.02 | 2.14±6.97 | G |
| *QdVegdm.huj.uh-5A* | 5A.2 | 2.9 | 29.1 | 16.5—45.1 | 28.6 | wsnp_CAP7_c2282_1107112 | 0.13 | 12.68±5.13 | -- | -- | G |
| *QddfVegdm.huj.uh-5A* | 5A.2 | 2.5 | 29.1 | 11.6—45.6 | 33.9 | wsnp_CAP7_c2282_1107112 | 0.11 | 11.36±3.23 | -- | -- | G |
| *QVegdm.huj.uh-7A* | 7A.1 | 2.7 | 2.8 | 0.0—10.1 | 10.1 | IAAV6131 | 0.06 | -5.25±7.23 | 0.09 | -12.80±10.69 | L |
| *QdfVegdm.huj.uh-7A* | 7A.1 | 4.4 | 7.8 | 5.8—19.0 | 13.2 | Tdurum_contig13048_89 | 0.07 | -5.81±6.96 | 0.09 | -13.46±9.98 | L |
| *QdVegdm.huj.uh-7B* | 7B.3 | 2.2 | 43.0 | 23.9—57.7 | 33.8 | wsnp_Ex_c23755_32994701 | 0.09 | -9.43±5.71 | -- | -- | L |
| **Total dry matter** | | | | | | | | | | | |
| *QTotdm.huj.uh-2A* | 2A.1 | 3.6 | 36.4 | 32.1—40.1 | 8.1 | wsnp_Ex_c2033_3814035 | 0.05 | -8.68±7.83 | 0.09 | -23.60±12.50 | L |

| QTL effects | QTL | LOD | Position (cM) | Interval (cM) | Length (cM) | Nearest marker | WL | | WW | | ITV Allele |
|---|---|---|---|---|---|---|---|---|---|---|---|
| | | | | | | | PEV | d | PEV | d | |
| *QdfTotdm.huj.uh-2A* | 2A.1 | 3.8 | 38.3 | 35.3—44.5 | 9.2 | BobWhite_c10977_834 | 0.04 | -7.34±7.39 | 0.08 | -21.81±12.56 | L |
| *QTotdm.huj.uh-2B* | 2B.6 | 5.4 | 103.8 | 101.7—108.7 | 7.0 | BobWhite_c47357_535 | 0.06 | -11.78±5.00 | 0.13 | -32.15±5.96 | L |
| *QdfTotdm.huj.uh-2B* | 2B.6 | 3.1 | 106.4 | 101.0—112.5 | 11.5 | Tdurum_contig12879_1200 | 0.05 | -6.09±9.44 | 0.08 | -20.50±12.70 | L |
| *QTotdm.huj.uh-4A* | 4A.1 | 2.9 | 17.1 | 5.8—20.3 | 14.5 | BS00065863_51 | 0.11 | -14.49±9.80 | 0.02 | -3.56±10.41 | L |
| *QdfTotdm.huj.uh-4A* | 4A.1 | 3.5 | 17.1 | 5.9—20.3 | 14.3 | BS00065863_51 | 0.12 | -15.01±10.12 | 0.02 | -4.52±10.29 | L |
| *QdTotdm.huj.uh-4A* | 4A.1 | 2.5 | 10.2 | 3.5—33.6 | 30.1 | BS00043286_51 | 0.13 | -17.31±7.74 | -- | -- | L |
| *QddfTotdm.huj.uh-4A* | 4A.1 | 2.6 | 17.1 | 4.5—32.8 | 28.3 | BS00065863_51 | 0.14 | -17.42±6.84 | -- | -- | L |
| *QTotdm.huj.uh-4B* | 4B.3 | 4.6 | 41.5 | 36.7—45.3 | 8.7 | Kukri_c32064_629 | 0.08 | -13.14±7.33 | 0.11 | -24.94±16.72 | L |
| *QdfTotdm.huj.uh-4B* | 4B.3 | 3.3 | 40.5 | 32.0—52.6 | 20.5 | RFL_Contig3563_1130 | 0.07 | -10.06±9.23 | 0.08 | -13.30±21.27 | L |
| *QTotdm.huj.uh-5A* | 5A.2 | 2.0 | 29.1 | 17.0—59.7 | 42.7 | wsnp_CAP7_c2282_1107112 | 0.13 | 18.44±4.47 | -- | -- | G |
| *QTotdm.huj.uh-7A* | 7A.1 | 3.3 | 6.0 | 0.0—11.2 | 11.1 | RAC875_c18446_506 | 0.07 | -6.95±11.62 | 0.09 | -20.19±16.29 | L |
| *QdfTotdm.huj.uh-7A* | 7A.1 | 3.5 | 6.0 | 0.0—10.6 | 10.6 | RAC875_c18446_506 | 0.07 | -7.19±11.75 | 0.09 | -21.16±15.67 | L |
| **Morphological traits** | | | | | | | | | | | |
| **Culm length** | | | | | | | | | | | |
| *QCl.huj.uh-2B* | 2B.6 | 5.8 | 103.8 | 101.0—111.4 | 10.4 | BobWhite_c47357_535 | 0.10 | -8.06±1.59 | 0.05 | -5.17±2.92 | L |
| *QdCl.huj.uh-2B* | 2B.6 | 4.0 | 103.8 | 93.8—117.4 | 23.6 | BobWhite_c47357_535 | 0.12 | -7.47±1.69 | -- | -- | L |
| *QCl.huj.uh-3A* | 3A.4 | 2.5 | 58.4 | 46.0—69.5 | 23.5 | wsnp_Ex_c2331_4369782 | 0.07 | -2.68±6.01 | 0.08 | -4.58±6.02 | L |
| *QdfCl.huj.uh-3A* | 3A.4 | 4.4 | 58.0 | 52.6—71.2 | 18.5 | IAAV3900 | 0.08 | -5.46±1.51 | 0.07 | -5.95±3.07 | L |
| *QdCl.huj.uh-3B.1* | 3B.2 | 2.7 | 72.2 | 60.9—83.0 | 22.1 | BS00032926_51 | 0.13 | -7.70±1.10 | -- | -- | L |
| *QCl.huj.uh-3B* | 3B.3 | 5.1 | 97.2 | 94.9—104.5 | 9.7 | BS00063624_51 | 0.08 | -6.91±1.54 | 0.04 | -4.72±2.73 | L |
| *QCl.huj.uh-4A* | 4A.3 | 9.5 | 41.8 | 37.5—44.0 | 6.5 | TA004056-0809 | 0.10 | -7.68±1.69 | 0.13 | -9.41±1.93 | L |
| *QdfCl.huj.uh-4A* | 4A.3 | 7.8 | 41.8 | 39.2—44.9 | 5.7 | TA004056-0809 | 0.10 | -6.37±1.30 | 0.08 | -7.26±1.84 | L |
| *QCl.huj.uh-4B.1* | 4B.4 | 7.2 | 44.1 | 38.3—48.4 | 10.1 | Ex_c25467_851 | 0.09 | -7.54±1.79 | 0.07 | -6.89±1.95 | L |
| *QdfCl.huj.uh-4B.2* | 4B.6 | 4.2 | 84.6 | 77.9—84.6 | 6.7 | wsnp_Ex_c19844_28854305 | 0.03 | -2.83±1.87 | 0.11 | -8.33±2.35 | L |
| *QddfCl.huj.uh-5A.1* | 5A.3 | 3.3 | 41.9 | 38.5—43.9 | 5.4 | wsnp_Ra_c14112_22155312 | 0.11 | 5.83±2.16 | -- | -- | G |
| *QCl.huj.uh-5A.2* | 5A.6 | 2.5 | 110.9 | 95.6—129.9 | 34.4 | Ku_c21235_676 | -- | -- | 0.11 | 7.64±5.53 | G |
| *QdfCl.huj.uh-5A.3* | 5A.9 | 3.8 | 157.0 | 152.2—157.1 | 4.9 | IACX5850 | 0.05 | -2.47±4.05 | 0.05 | -4.62±3.78 | L |
| *QCl.huj.uh-5B* | 5B.8 | 4.4 | 133.5 | 129.1—137.7 | 8.6 | BS00050057_51 | 0.08 | 6.66±2.35 | 0.12 | 8.96±2.21 | G |
| *QdfCl.huj.uh-6B* | 6B.2 | 4.1 | 37.1 | 24.0—47.0 | 23.1 | BS00109708_51 | 0.10 | -6.39±1.41 | 0.03 | -3.54±2.74 | L |
| *QddfCl.huj.uh-6B* | 6B.2 | 2.5 | 30.1 | 22.1—49.9 | 27.8 | TA003005-0339 | 0.10 | -5.90±1.06 | -- | -- | L |
| *QCl.huj.uh-7A.1* | 7A.5 | 5.1 | 73.0 | 63.5—82.1 | 18.6 | Kukri_rep_c70389_57 | 0.07 | -6.11±2.31 | 0.04 | -4.38±2.80 | L |
| *QdfCl.huj.uh-7A.1* | 7A.5 | 5.4 | 78.5 | 71.4—84.2 | 12.8 | wsnp_Ex_c11636_18742884 | 0.10 | -6.08±2.28 | 0.05 | -5.29±2.62 | L |
| *QddfCl.huj.uh-7A.2* | 7A.7 | 2.9 | 104.5 | 94.7—111.7 | 17.0 | wsnp_Ex_c9428_15641639 | 0.11 | -5.84±1.61 | -- | -- | L |
| *QCl.huj.uh-7B* | 7B.1 | 7.5 | 11.5 | 5.8—18.1 | 12.3 | wsnp_CAP8_c334_304253 | 0.10 | 7.68±1.53 | 0.08 | 7.21±1.81 | G |
| **Spike length** | | | | | | | | | | | |
| *QSpl.huj.uh-1A* | 1A.2 | 3.9 | 53.0 | 50.1—56.0 | 6.0 | wsnp_Ex_c572_1138503 | 0.12 | 1.49±0.30 | 0.06 | 1.29±0.49 | G |
| *QdfSpl.huj.uh-1A* | 1A.2 | 2.3 | 53.0 | 48.8—59.7 | 10.9 | wsnp_Ex_c572_1138503 | 0.05 | 0.89±0.28 | 0.02 | 0.63±0.69 | G |
| *QSpl.huj.uh-2A* | 2A.5 | 4.6 | 130.8 | 126.3—133.0 | 6.8 | BS00065366_51 | 0.07 | -1.06±0.54 | 0.02 | -0.59±0.68 | L |
| *QSpl.huj.uh-4A.1* | 4A.3 | 2.6 | 41.8 | 22.9—51.1 | 28.2 | TA004056-0809 | 0.12 | -0.99±1.18 | -- | -- | L |
| *QddfSpl.huj.uh-4A.1* | 4A.3 | 2.6 | 41.8 | 33.0—51.1 | 18.1 | TA004056-0809 | 0.08 | -0.97±0.26 | -- | -- | L |

| QTL effects | QTL | LOD | Position (cM) | Interval (cM) | Length (cM) | Nearest marker | WL | | WW | | ITV Allele |
|---|---|---|---|---|---|---|---|---|---|---|---|
| | | | | | | | PEV | d | PEV | d | |
| *QSpl.huj.uh-4A.2* | 4A.6 | 3.4 | 101.4 | 90.8—108.9 | 18.1 | RAC875_c35979_263 | -- | -- | 0.10 | 1.82±0.62 | G |
| *QSpl.huj.uh-5A* | 5A.3 | 10.8 | 41.9 | 38.6—43.5 | 4.9 | wsnp_Ra_c14112_22155312 | 0.17 | 1.89±0.24 | 0.03 | 0.87±0.41 | G |
| *QdfSpl.huj.uh-5A* | 5A.3 | 10.1 | 41.9 | 39.0—43.7 | 4.7 | wsnp_Ra_c14112_22155312 | 0.18 | 1.86±0.28 | 0.04 | 1.13±0.44 | G |
| *QdSpl.huj.uh-5A* | 5A.3 | 5.5 | 39.8 | 36.1—45.2 | 9.1 | BS00077990_51 | 0.17 | 1.58±0.24 | -- | -- | G |
| *QddfSpl.huj.uh-5A* | 5A.3 | 4.5 | 39.1 | 32.8—56.0 | 23.1 | Ra_c69221_1167 | 0.16 | 1.42±0.21 | -- | -- | G |
| *QSpl.huj.uh-5B* | 5B.8 | 7.4 | 133.5 | 130.4—136.3 | 5.9 | BS00050057_51 | 0.06 | 1.13±0.25 | 0.08 | 1.63±0.37 | G |
| *QSpl.huj.uh-6A* | 6A.4 | 5.1 | 69.4 | 58.4—72.9 | 14.6 | BobWhite_c47143_592 | 0.09 | -1.34±0.22 | 0.02 | -0.72±0.47 | L |
| *QdfSpl.huj.uh-6A* | 6A.4 | 4.0 | 69.4 | 59.62—76.34 | 16.7 | BobWhite_c47143_592 | 0.08 | -1.15±0.46 | 0.02 | -0.56±0.59 | L |
| *QdSpl.huj.uh-6A* | 6A.4 | 2.3 | 74.1 | 54.7—98.4 | 43.7 | RFL_Contig5314_1147 | 0.09 | -1.11±0.23 | -- | -- | L |
| *QddfSpl.huj.uh-6A* | 6A.4 | 2.0 | 73.1 | 50.1—103.8 | 53.7 | BS00065152_51 | 0.07 | -0.91±0.28 | -- | -- | L |
| *QdfSpl.huj.uh-6B* | 6B.2 | 3.1 | 36.8 | 30.0—50.6 | 20.6 | IACX6023 | 0.05 | 0.96±0.33 | 0.04 | 1.09±0.45 | G |
| *QSpl.huj.uh-7A* | 7A.5 | 5.5 | 79.6 | 74.0—86.3 | 12.3 | Kukri_c64283_212 | 0.08 | 1.27±0.22 | 0.04 | 1.08±0.40 | G |
| *QdfSpl.huj.uh-7A* | 7A.5 | 3.8 | 80.7 | 72.8—91.9 | 19.1 | Kukri_c26517_691 | 0.09 | 1.25±0.30 | 0.03 | 0.93±0.51 | G |
| *QSpl.huj.uh-7B* | 7B.1 | 8.2 | 14.9 | 13.4—21.2 | 7.8 | Tdurum_contig42423_1963 | 0.10 | -1.45±0.26 | 0.06 | -1.45±0.39 | L |
| **Flag leaf length** | | | | | | | | | | | |
| *QFll.huj.uh-1B* | 1B.1 | 3.2 | 6.2 | 0.4—11.4 | 11.0 | TA012713-0667 | 0.06 | -1.31±0.65 | 0.06 | -1.52±0.61 | L |
| *QFll.huj.uh-2A* | 2A.4 | 2.1 | 82.5 | 70.3—94.6 | 24.3 | Excalibur_c34913_743 | 0.08 | 1.55±0.75 | 0.06 | 1.49±0.78 | G |
| *QFll.huj.uh-3A* | 3A.5 | 2.9 | 72.4 | 68.8—78.2 | 9.3 | IAAV8483 | 0.12 | -2.09±0.44 | 0.03 | -0.80±0.86 | L |
| *QFll.huj.uh-4B* | 4B.4 | 3.0 | 45.5 | 41.1—51.0 | 9.9 | RAC875_c18736_133 | 0.12 | 1.92±0.94 | 0.05 | 1.02±1.08 | G |
| *QFll.huj.uh-5A.1* | 5A.3 | 4.5 | 41.9 | 38.8—45.5 | 6.7 | wsnp_Ra_c14112_22155312 | 0.25 | 1.92±1.47 | 0.16 | 1.37±1.12 | G |
| *QdfFll.huj.uh-5A.1* | 5A.3 | 5.4 | 43.3 | 38.5—47.1 | 8.5 | BobWhite_rep_c49979_373 | 0.22 | 1.64±0.96 | 0.14 | 1.11±1.05 | G |
| *QddfFll.huj.uh-5A.1* | 5A.3 | 2.8 | 43.3 | 37.4—52.7 | 15.3 | BobWhite_rep_c49979_373 | 0.14 | 0.26±1.95 | -- | -- | G |
| *QFll.huj.uh-5A.2* | 5A.8 | 7.7 | 146.9 | 141.8—154.1 | 12.3 | wsnp_Ex_rep_c101323_86702546 | 0.25 | -2.05±1.19 | 0.16 | -2.03±0.95 | L |
| *QdfFll.huj.uh-5A.2* | 5A.8 | 6.4 | 153.5 | 144.4—148.4 | 4.0 | tplb0049a09_1302 | 0.22 | -1.62±0.95 | 0.14 | -2.00±0.84 | L |
| *QdFll.huj.uh-5A.2* | 5A.8 | 3.3 | 145.8 | 140.6—157.0 | 16.5 | wsnp_Ex_c3136_5798236 | 0.13 | -1.23±1.87 | -- | -- | L |
| *QFll.huj.uh-6A.1* | 6A.1 | 3.0 | 11.3 | 0.5—24.5 | 24.0 | Tdurum_contig97611_150 | 0.17 | -0.04±2.50 | 0.17 | -1.78±2.25 | L |
| *QdFll.huj.uh-6A.1* | 6A.1 | 2.0 | 12.3 | 0.0—29.3 | 29.3 | Tdurum_contig54957_624 | 0.08 | -1.71±0.38 | -- | -- | L |
| *QFll.huj.uh-6A.2* | 6A.4 | 2.6 | 69.4 | 63.4—86.0 | 22.7 | BobWhite_c47143_592 | 0.11 | -1.07±1.57 | 0.11 | -0.24±1.99 | L |
| *QdfFll.huj.uh-6A.2* | 6A.4 | 3.3 | 69.4 | 66.23—72.12 | 5.9 | BobWhite_c47143_592 | 0.05 | -1.20±0.48 | 0.08 | -1.79±0.74 | L |
| *QFll.huj.uh-7A* | 7A.3 | 2.3 | 36.8 | 26.2—40.5 | 14.3 | Tdurum_contig82438_73 | 0.08 | -1.37±1.09 | 0.07 | -1.34±1.12 | L |
| **Flag leaf width** | | | | | | | | | | | |
| *QFlw.huj.uh-1A* | 1A.1 | 4.1 | 27.6 | 8.1—32.0 | 23.9 | GENE-0002_856 | 0.03 | -0.07±0.02 | 0.05 | -0.11±0.02 | L |
| *QdfFlw.huj.uh-1A* | 1A.1 | 4.7 | 11.3 | 6.6—16.0 | 9.4 | Excalibur_c9676_163 | 0.04 | -0.07±0.02 | 0.03 | -0.09±0.02 | L |
| *QFlw.huj.uh-2A* | 2A.2 | 3.0 | 55.1 | 32.1—63.2 | 31.1 | Kukri_rep_c73477_888 | 0.10 | -0.12±0.03 | 0.05 | -0.09±0.05 | L |
| *QdfFlw.huj.uh-2A* | 2A.2 | 2.9 | 56.6 | 34.6—64.2 | 29.6 | Tdurum_contig5311_67 | 0.11 | -0.12±0.03 | 0.05 | -0.09±0.05 | L |
| *QFlw.huj.uh-2B.1* | 2B.6 | 5.5 | 103.8 | 98.6—110.8 | 12.2 | BobWhite_c47357_535 | 0.02 | -0.04±0.03 | 0.08 | -0.15±0.02 | L |
| *QdfFlw.huj.uh-2B.1* | 2B.6 | 6.7 | 106.4 | 98.1—111.5 | 13.4 | Tdurum_contig12879_1200 | 0.02 | -0.05±0.03 | 0.07 | -0.13±0.03 | L |
| *QdFlw.huj.uh-2B.2* | 2B.7 | 2.1 | 118.4 | 110.5—132.5 | 22.1 | BS00068181_51 | 0.08 | 0.08±0.01 | -- | -- | G |
| *QFlw.huj.uh-3A* | 3A.1 | 4.1 | 8.3 | 0.9—17.0 | 16.1 | Ra_c6118_350 | 0.07 | 0.09±0.05 | 0.10 | 0.14±0.06 | G |
| *QdfFlw.huj.uh-3A* | 3A.1 | 5.2 | 11.9 | 6.9—18.3 | 11.4 | wsnp_JD_c2722_3653988 | 0.03 | 0.06±0.03 | 0.04 | 0.10±0.03 | G |
| *QFlw.huj.uh-3B* | 3B.4 | 6.7 | 118.2 | 115.9—122.4 | 6.5 | BobWhite_c15763_205 | 0.05 | 0.09±0.02 | 0.05 | 0.11±0.03 | G |

| QTL effects | QTL | LOD | Position (cM) | Interval (cM) | Length (cM) | Nearest marker | WL | | WW | | ITV Allele |
|---|---|---|---|---|---|---|---|---|---|---|---|
| | | | | | | | PEV | d | PEV | d | |
| *QdfFlw.huj.uh-3B* | 3B.4 | 5.0 | 129.9 | 124.9—133.6 | 8.6 | Tdurum_contig100787_79 | 0.05 | 0.09±0.02 | 0.03 | 0.08±0.03 | G |
| *QFlw.huj.uh-4A.1* | 4A.5 | 2.8 | 74.4 | 62.7—78.9 | 16.2 | Kukri_c19216_123 | 0.03 | 0.02±0.07 | 0.10 | 0.12±0.10 | G |
| *QFlw.huj.uh-4A.2* | 4A.7 | 6.2 | 120.9 | 118.1—123.5 | 5.4 | RAC875_rep_c69632_65 | 0.13 | 0.01±0.08 | 0.09 | 0.07±0.12 | G |
| *QdfFlw.huj.uh-4A.2* | 4A.7 | 4.3 | 120.9 | 113.1—123.9 | 10.8 | RAC875_rep_c69632_65 | 0.00 | 0.00±0.03 | 0.06 | 0.12±0.03 | G |
| *QdFlw.huj.uh-4A.2* | 4A.7 | 3.7 | 120.9 | 110.2—125.0 | 14.8 | RAC875_rep_c69632_65 | 0.13 | 0.10±0.02 | -- | -- | G |
| *QddfFlw.huj.uh-4A.2* | 4A.7 | 3.0 | 120.9 | 108.7—125.6 | 16.9 | RAC875_rep_c69632_65 | 0.11 | 0.08±0.05 | -- | -- | G |
| *QFlw.huj.uh-4B* | 4B.5 | 26.6 | 61.0 | 60.3—63.0 | 2.7 | RAC875_c24515_602 | 0.25 | -0.20±0.03 | 0.25 | -0.26±0.03 | L |
| *QdfFlw.huj.uh-4B* | 4B.5 | 32.4 | 61.0 | 60.4—62.2 | 1.8 | RAC875_c24515_602 | 0.28 | -0.21±0.02 | 0.27 | -0.26±0.03 | L |
| *QFlw.huj.uh-5A* | 5A.5 | 18.8 | 100.3 | 98.0—102.9 | 4.9 | Tdurum_contig50175_875 | 0.20 | -0.18±0.02 | 0.12 | -0.18±0.03 | L |
| *QdfFlw.huj.uh-5A* | 5A.5 | 16.1 | 99.5 | 97.8—102 | 4.2 | RAC875_rep_c109969_119 | 0.14 | -0.15±0.02 | 0.08 | -0.14±0.03 | L |
| *QdFlw.huj.uh-5A* | 5A.5 | 2.9 | 97.0 | 91.8—120.8 | 29.0 | Excalibur_c23354_306 | 0.12 | -0.10±0.02 | -- | -- | L |
| *QddfFlw.huj.uh-5A* | 5A.5 | 2.3 | 97.0 | 91.0—126.7 | 35.7 | Excalibur_c23354_306 | 0.12 | -0.10±0.02 | -- | -- | L |
| *QdfFlw.huj.uh-5B.1* | 5B.3 | 4.8 | 56.1 | 49.0—67.5 | 18.5 | Ra_c35412_1057 | 0.11 | -0.01±0.13 | 0.14 | -0.07±0.10 | L |
| *QdfFlw.huj.uh-5B.2* | 5B.7 | 3.0 | 117.4 | 107.0—135.0 | 28.0 | BS00098292_51 | 0.11 | 0.04±0.11 | 0.14 | 0.12±0.07 | G |
| *QdfFlw.huj.uh-5B.3* | 5B.8 | 6.1 | 132.8 | 126.5—137.6 | 11.1 | BS00084106_51 | -- | -- | 0.08 | 0.14±0.03 | G |
| *QFlw.huj.uh-6A* | 6A.2 | 8.8 | 49.1 | 45.6—52.5 | 6.9 | IACX14305 | 0.06 | -0.10±0.02 | 0.06 | -0.13±0.03 | L |
| *QdfFlw.huj.uh-6A* | 6A.2 | 8.3 | 49.1 | 45.03—53.47 | 8.4 | IACX14305 | 0.05 | -0.09±0.02 | 0.05 | -0.12±0.02 | L |
| *QFlw.huj.uh-6B* | 6B.4 | 3.2 | 51.5 | 47.4—58.4 | 11.0 | BobWhite_c1059_1825 | -- | -- | 0.05 | -0.10±0.04 | L |
| *QdfFlw.huj.uh-7A* | 7A.6 | 4.8 | 87.3 | 81.8—101.2 | 19.4 | BS00109319_51 | 0.03 | 0.06±0.02 | 0.06 | 0.12±0.02 | G |
| *QFlw.huj.uh-7B* | 7B.4 | 5.7 | 128.4 | 125.4—128.4 | 3.0 | RFL_Contig3607_648 | 0.11 | -0.13±0.03 | 0.10 | -0.14±0.04 | L |
| **Drought adaptive physiological traits** | | | | | | | | | | | |
| **Carbon isotope ratio** | | | | | | | | | | | |
| *Qδ13c.huj.uh-1A* | 1A.2 | 4.4 | 59.8 | 48.7—66.9 | 18.2 | IAAV5535 | -- | -- | 0.13 | 0.36±0.07 | L |
| *Qdfδ13c.huj.uh-1A* | 1A.2 | 4.5 | 59.8 | 48.4—67.3 | 18.9 | IAAV5535 | 0.02 | 0.03±0.08 | 0.12 | 0.34±0.08 | L |
| *Qδ13c.huj.uh-3A.1* | 3A.1 | 2.2 | 8.3 | 0.0—18.1 | 18.0 | Ra_c6118_350 | 0.07 | -0.07±0.19 | 0.11 | -0.03±0.16 | G |
| *Qddfδ13c.huj.uh-3A.1* | 3A.1 | 2.5 | 11.9 | 3.4—18.0 | 14.6 | wsnp_JD_c2722_3653988 | 0.10 | -0.22±0.09 | -- | -- | G |
| *Qδ13c.huj.uh-3A.2* | 3A.5 | 6.6 | 68.1 | 64.9—73.8 | 8.9 | wsnp_Ex_c18223_27035083 | 0.07 | -0.06±0.13 | 0.11 | -0.25±0.18 | G |
| *Qdfδ13c.huj.uh-3A.2* | 3A.5 | 5.7 | 68.1 | 65.0—74.9 | 9.9 | wsnp_Ex_c18223_27035083 | 0.02 | -0.05±0.08 | 0.13 | -0.36±0.10 | G |
| *Qδ13c.huj.uh-4A* | 4A.3 | 8.0 | 41.8 | 31.4—45.8 | 14.4 | TA004056-0809 | 0.14 | -0.30±0.06 | 0.06 | -0.23±0.08 | G |
| *Qdfδ13c.huj.uh-4A* | 4A.3 | 9.3 | 36.4 | 31.7—45.6 | 13.8 | RAC875_c31051_490 | 0.19 | -0.35±0.06 | 0.06 | -0.25±0.08 | G |
| *Qdδ13c.huj.uh-4A* | 4A.3 | 3.1 | 24.0 | 22.6—47.6 | 25.0 | Kukri_rep_c101034_516 | 0.16 | -0.30±0.06 | -- | -- | G |
| *Qddfδ13c.huj.uh-4A* | 4A.3 | 3.1 | 36.4 | 23.2—48.3 | 25.1 | RAC875_c31051_490 | 0.11 | -0.25±0.04 | -- | -- | G |
| *Qδ13c.huj.uh-5A.1* | 5A.4 | 5.6 | 69.2 | 63.3—73.9 | 10.6 | Kukri_c63163_141 | 0.10 | -0.25±0.06 | 0.07 | -0.26±0.09 | G |
| *Qdfδ13c.huj.uh-5A.1* | 5A.4 | 7.0 | 68.1 | 63.1—72.4 | 9.4 | Ex_c104539_35 | 0.11 | -0.26±0.06 | 0.08 | -0.28±0.07 | G |
| *Qdδ13c.huj.uh-5A.2* | 5A.7 | 2.3 | 133.7 | 129.1—140.5 | 11.4 | wsnp_Ex_c54211_57168122 | 0.10 | -0.26±0.04 | -- | -- | G |
| *Qδ13c.huj.uh-6B* | 6B.4 | 5.1 | 55.7 | 42.6—60.5 | 18.0 | Tdurum_contig93283_513 | 0.08 | -0.22±0.05 | 0.07 | -0.27±0.06 | G |
| *Qdfδ13c.huj.uh-6B* | 6B.4 | 4.3 | 55.7 | 43.1—61.5 | 18.5 | Tdurum_contig93283_513 | 0.07 | -0.21±0.06 | 0.06 | -0.24±0.09 | G |
| *Qδ13c.huj.uh-7B* | 7B.2 | 5.6 | 23.3 | 14.6—28.0 | 13.4 | Tdurum_contig99020_139 | 0.14 | -0.31±0.04 | 0.01 | -0.03±0.08 | G |
| *Qdδ13c.huj.uh-7B* | 7B.2 | 6.0 | 23.7 | 21.0—27.5 | 6.5 | BobWhite_c44404_312 | 0.17 | -0.33±0.05 | -- | -- | G |
| *Qddfδ13c.huj.uh-7B* | 7B.2 | 3.9 | 23.7 | 21.5—31.7 | 10.2 | BobWhite_c44404_312 | 0.13 | -0.27±0.07 | -- | -- | G |

| QTL effects | QTL | LOD | Position (cM) | Interval (cM) | Length (cM) | Nearest marker | WL | | WW | | ITV Allele |
|---|---|---|---|---|---|---|---|---|---|---|---|
| | | | | | | | PEV | d | PEV | d | |
| **Osmotic potential at heading** | | | | | | | | | | | |
| *QdfOp.huj.uh-1A* | 1A.3 | 2.3 | 94.3 | 81.3—110.6 | 29.3 | Ra_c41164_730 | -- | -- | 0.11 | 0.74±0.13 | L |
| *QOp.huj.uh-2A* | 2A.3 | 3.9 | 64.3 | 52.8—70.8 | 18.0 | Tdurum_contig27887_55 | -- | -- | 0.15 | 0.96±0.12 | L |
| *QdfOp.huj.uh-2A* | 2A.3 | 4.0 | 66.2 | 51.4—71.6 | 20.2 | wsnp_Ex_c22862_32074455 | 0.02 | 0.00±0.39 | 0.13 | 0.80±0.28 | L |
| *QOp.huj.uh-2B* | 2B.3 | 3.2 | 75.6 | 66.5—87.9 | 21.4 | RFL_Contig4718_1269 | 0.04 | 0.36±0.22 | 0.14 | 0.62±0.19 | L |
| *QdfOp.huj.uh-2B* | 2B.3 | 4.0 | 66.2 | 51.4—71.6 | 20.2 | wsnp_Ex_c22862_32074455 | 0.02 | 0.00±0.39 | 0.13 | 0.80±0.28 | L |
| *QOp.huj.uh-3B* | 3B.2 | 4.5 | 64.7 | 61.8—72.0 | 10.2 | BS00032926_51 | 0.10 | 0.88±0.19 | 0.08 | 0.65±0.22 | L |
| *QdfOp.huj.uh-3B* | 3B.2 | 3.8 | 72.6 | 65.5—75.7 | 10.2 | RAC875_c16281_195 | 0.10 | 0.81±0.28 | 0.06 | 0.52±0.25 | L |
| *QOp.huj.uh-4B* | 4B.5 | 3.7 | 60.7 | 50.3—70.8 | 20.5 | Tdurum_contig59914_323 | 0.16 | 1.12±0.24 | 0.04 | 0.16±0.44 | L |
| *QdfOp.huj.uh-4B* | 4B.5 | 3.3 | 65.2 | 50.6—59.6 | 9.0 | IAAV8848 | 0.12 | 0.87±0.35 | 0.05 | 0.25±0.47 | L |
| *QdOp.huj.uh-4B* | 4B.5 | 3.4 | 65.2 | 55.3—75.2 | 19.9 | IAAV8848 | 0.19 | 1.11±0.24 | -- | -- | L |
| *QddfOp.huj.uh-4B* | 4B.5 | 3.3 | 65.2 | 55.0—75.7 | 20.8 | IAAV8848 | 0.18 | 1.07±0.30 | -- | -- | L |
| *QdOp.huj.uh-5A* | 5A.6 | 2.1 | 128.0 | 99.8—145.0 | 45.1 | wsnp_Ex_c54655_57455110 | 0.11 | 0.62±0.17 | -- | -- | L |
| *QOp.huj.uh-5B* | 5B.5 | 2.9 | 106.9 | 103.6—109.5 | 5.9 | BobWhite_c28333_454 | 0.07 | -0.50±0.20 | 0.08 | -0.46±0.19 | G |
| *QdfOp.huj.uh-5B* | 5B.5 | 2.9 | 106.9 | 103.3—109.8 | 6.5 | BobWhite_c28333_454 | 0.07 | -0.49±0.23 | 0.06 | -0.37±0.21 | G |
| *QOp.huj.uh-6A* | 6A.3 | 2.8 | 53.5 | 46.7—62.8 | 16.1 | Tdurum_contig76709_195 | -- | -- | 0.10 | 0.58±0.12 | L |
| *QdfOp.huj.uh-6A* | 6A.3 | 3.2 | 53.5 | 44.7—59.5 | 14.8 | Tdurum_contig76709_195 | 0.02 | 0.11±0.27 | 0.11 | 0.59±0.12 | L |
| *QOp.huj.uh-6B* | 6B.2 | 4.5 | 36.4 | 32.0—39.2 | 7.2 | RAC875_c13920_747 | 0.05 | 0.40±0.18 | 0.10 | 0.58±0.13 | L |
| *QdfOp.huj.uh-6B* | 6B.2 | 4.9 | 36.4 | 32.1—39.1 | 7.0 | RAC875_c13920_747 | 0.05 | 0.45±0.17 | 0.12 | 0.61±0.12 | L |
| **Chlorophyll content at heading** | | | | | | | | | | | |
| *QChl.huj.uh-2A* | 2A.2 | 3.0 | 43.6 | 38.1—57.6 | 19.6 | Excalibur_rep_c105284_110 | -- | -- | 0.11 | -2.47±0.39 | L |
| *QdfChl.huj.uh-2A* | 2A.2 | 3.8 | 43.6 | 36.9—51.7 | 14.9 | Excalibur_rep_c105284_110 | 0.08 | -1.85±0.60 | 0.09 | -2.12±0.69 | L |
| *QdfChl.huj.uh-2B.1* | 2B.3 | 3.3 | 78.2 | 69.0—88.0 | 19.0 | Kukri_c4097_898 | 0.06 | -0.10±1.81 | 0.07 | -1.27±0.98 | L |
| *QChl.huj.uh-2B.2* | 2B.5 | 4.4 | 98.5 | 77.7—111.0 | 33.2 | Excalibur_rep_c86807_132 | 0.05 | -1.15±1.23 | 0.11 | -1.95±0.78 | L |
| *QChl.huj.uh-3A* | 3A.4 | 2.4 | 60.1 | 49.6—69.1 | 19.5 | BS00011612_51 | -- | -- | 0.06 | 0.99±1.08 | G |
| *QdChl.huj.uh-4A* | 4A.2 | 2.1 | 28.4 | 22.1—34.5 | 12.5 | Ku_c3891_395 | 0.09 | 1.41±1.38 | -- | -- | G |
| *QddfChl.huj.uh-4A* | 4A.2 | 2.0 | 28.4 | 22.1—34.7 | 12.5 | Ku_c3891_395 | 0.09 | 1.45±1.36 | -- | -- | G |
| *QChl.huj.uh-4B.1* | 4B.1 | 2.9 | 4.1 | 0.0—21.4 | 21.4 | IAAV7323 | 0.09 | 1.99±0.64 | 0.08 | 1.81±0.80 | G |
| *QdfChl.huj.uh-4B.1* | 4B.2 | 7.0 | 16.0 | 12.9—20.8 | 7.9 | Tdurum_contig76213_1156 | 0.09 | 2.19±0.65 | 0.11 | 2.04±0.52 | G |
| *QChl.huj.uh-5A* | 5A.3 | 4.6 | 45.5 | 33.2—53.4 | 20.2 | wsnp_Ex_rep_c70343_69286072 | 0.06 | -1.60±0.85 | 0.10 | -1.98±0.43 | L |
| *QdfChl.huj.uh-5A* | 5A.3 | 3.8 | 35.4 | 32.8—41 | 8.2 | BS00021873_51 | 0.05 | -1.34±0.85 | 0.13 | -2.04±0.51 | L |
| *QChl.huj.uh-5B* | 5B.6 | 4.7 | 112.2 | 103.8—119.6 | 15.9 | Tdurum_contig70554_1004 | 0.09 | 1.97±0.52 | 0.12 | 2.49±0.51 | G |
| *QdfChl.huj.uh-5B* | 5B.6 | 2.9 | 111.8 | 96.3—122.4 | 26.1 | Tdurum_contig47833_484 | 0.06 | 0.71±1.62 | 0.10 | 1.41±1.28 | G |
| *QChl.huj.uh-6A.1* | 6A.2 | 2.8 | 48.4 | 38.3—54.3 | 16.0 | wsnp_Ex_c19770_28768859 | 0.09 | -1.93±0.73 | 0.03 | -0.96±0.93 | L |
| *QdfChl.huj.uh-6A.1* | 6A.2 | 2.8 | 47.7 | 37.3—53.9 | 16.6 | BS00012297_51 | 0.08 | -1.84±0.76 | 0.05 | -1.30±0.92 | L |
| *QdChl.huj.uh-6A.1* | 6A.2 | 2.5 | 47.7 | 40.2—52.8 | 12.6 | BS00012297_51 | 0.10 | -1.85±0.78 | -- | -- | L |
| *QddfChl.huj.uh-6A.1* | 6A.2 | 2.5 | 47.7 | 40.0—52.8 | 12.8 | BS00012297_51 | 0.09 | -1.74±0.94 | -- | -- | L |
| *QdfChl.huj.uh-6A.2* | 6A.4 | 6.4 | 65.8 | 63.6—68.3 | 4.7 | wsnp_Ra_c12086_19452422 | 0.04 | 1.44±0.63 | 0.12 | 2.13±0.41 | G |
| *QdfChl.huj.uh-6B* | 6B.5 | 4.7 | 99.8 | 93.0—105.0 | 12.1 | Excalibur_c4261_342 | 0.08 | 2.07±0.51 | 0.07 | 1.69±0.37 | G |
| *QChl.huj.uh-7A* | 7A.6 | 4.8 | 86.6 | 78.9—91.8 | 12.9 | Kukri_rep_c105999_572 | 0.07 | 1.78±0.80 | 0.09 | 1.86±0.41 | G |

| QTL effects | QTL | LOD | Position (cM) | Interval (cM) | Length (cM) | Nearest marker | WL | | WW | | ITV Allele |
|---|---|---|---|---|---|---|---|---|---|---|---|
| | | | | | | | PEV | d | PEV | d | |
| *QdfChl.huj.uh-7A* | 7A.6 | 2.8 | 86.6 | 77.5—92.2 | 14.7 | Kukri_rep_c105999_572 | 0.06 | 1.17±1.36 | 0.08 | 1.48±0.73 | G |
| *QdfChl.huj.uh-7B* | 7B.4 | 5.2 | 128.4 | 121.8—128.4 | 6.6 | RFL_Contig3607_648 | 0.04 | -1.47±0.63 | 0.10 | -2.00±0.40 | L |
| **Flag leaf rolling index** | | | | | | | | | | | |
| *QdLr.huj.uh-1B* | 1B.3 | 2.1 | 40.1 | 28.6—52.0 | 23.4 | IAAV1158 | 0.10 | -0.31±0.16 | -- | -- | L |
| *QLr.huj.uh-2A* | 2A.5 | 3.3 | 133.7 | 126.9—140.0 | 13.1 | Tdurum_contig93508_295 | 0.06 | 0.27±0.10 | 0.09 | 0.33±0.10 | G |
| *QdfLr.huj.uh-2A* | 2A.5 | 3.2 | 134.4 | 129.9—140.7 | 10.8 | BS00022813_51 | 0.06 | 0.26±0.10 | 0.08 | 0.28±0.10 | G |
| *QLr.huj.uh-5A* | 5A.3 | 5.4 | 47.7 | 43.9—56.1 | 12.3 | Excalibur_c53930_53 | 0.07 | 0.28±0.11 | 0.13 | 0.39±0.08 | G |
| *QdfLr.huj.uh-5A* | 5A.3 | 5.5 | 47.7 | 43.5—54.7 | 11.2 | Excalibur_c53930_53 | 0.06 | 0.27±0.11 | 0.14 | 0.40±0.08 | G |
| *QLr.huj.uh-5B* | 5B.3 | 3.4 | 83.3 | 60.5—86.4 | 25.9 | RAC875_c32611_347 | 0.08 | 0.29±0.12 | 0.10 | 0.33±0.09 | G |
| *QLr.huj.uh-7A* | 7A.4 | 5.8 | 61.3 | 56.1—64.3 | 8.1 | RAC875_c67063_703 | 0.14 | -0.43±0.09 | 0.05 | -0.23±0.10 | L |
| *QdfLr.huj.uh-7A* | 7A.4 | 5.9 | 61.3 | 56.1—64.4 | 8.3 | RAC875_c67063_703 | 0.14 | -0.43±0.08 | 0.05 | -0.23±0.09 | L |
| *QdLr.huj.uh-7A* | 7A.4 | 3.0 | 61.3 | 55.3—65.5 | 10.2 | RAC875_c67063_703 | 0.12 | -0.37±0.08 | -- | -- | L |
| *QddfLr.huj.uh-7A* | 7A.4 | 3.1 | 61.3 | 53.2—65.6 | 12.4 | RAC875_c67063_703 | 0.14 | -0.38±0.08 | -- | -- | L |
| **Phenology related traits** | | | | | | | | | | | |
| **Days from planting to heading** | | | | | | | | | | | |
| *QDph.huj.uh-1B* | 1B.1 | 3.3 | 6.2 | 0.0—8.7 | 8.7 | TA012713-0667 | 0.08 | -2.39±2.66 | 0.08 | -1.46±2.16 | L |
| *QDph.huj.uh-2B* | 2B.6 | 21.7 | 106.4 | 103.0—107.9 | 5.0 | Tdurum_contig12879_1200 | 0.17 | 5.03±0.58 | 0.14 | 3.51±0.48 | G |
| *QDph.huj.uh-3A* | 3A.3 | 7.5 | 44.6 | 38.7—49.0 | 10.3 | wsnp_Ex_rep_c101942_87217430 | 0.03 | -2.01±0.60 | 0.07 | -2.48±0.45 | L |
| *QDph.huj.uh-4B* | 4B.4 | 2.4 | 45.5 | 37.1—50.5 | 13.4 | RAC875_c18736_133 | 0.08 | 3.35±0.96 | 0.07 | 2.44±0.91 | G |
| *QDph.huj.uh-5A.1* | 5A.3 | 2.8 | 41.9 | 36.9—49.9 | 12.9 | wsnp_Ra_c14112_22155312 | 0.11 | 1.71±2.45 | 0.07 | 0.60±1.90 | G |
| *QDph.huj.uh-5A.2* | 5A.5 | 4.9 | 100.3 | 88—106.5 | 18.5 | Tdurum_contig50175_875 | 0.11 | -3.82±1.28 | 0.11 | -2.98±0.91 | L |
| *QDph.huj.uh-5A.3* | 5A.8 | 7.1 | 146.9 | 140.3—149.7 | 9.3 | wsnp_Ex_rep_c101323_86702546 | 0.11 | -2.38±2.02 | 0.07 | -1.28±1.62 | L |
| *QDph.huj.uh-5B* | 5B.7 | 4.6 | 120.3 | 112.7—123.7 | 11.0 | wsnp_RFL_Contig1548_762547 | 0.03 | -2.20±0.56 | 0.04 | -1.81±0.43 | L |
| *QDph.huj.uh-6B* | 6B.1 | 2.4 | 3.9 | 0.0—17.0 | 16.9 | CAP11_c1087_327 | 0.08 | -3.19±1.24 | 0.07 | -2.21±1.13 | L |
| *QDph.huj.uh-7B* | 7B.1 | 40.8 | 14.9 | 14.1—15.7 | 1.6 | Tdurum_contig42423_1963 | 0.37 | -7.45±0.62 | 0.37 | -5.75±0.50 | L |
| **Days from heading to maturity** | | | | | | | | | | | |
| *QDhm.huj.uh-2B* | 2B.6 | 10.4 | 106.4 | 102.6—108.4 | 5.8 | Tdurum_contig12879_1200 | 0.15 | -3.47±0.46 | 0.08 | -1.38±0.33 | L |
| *QdDhm.huj.uh-2B* | 2B.6 | 4.5 | 110.7 | 101.4—115.7 | 14.3 | Ra_c13298_783 | 0.12 | -2.50±1.46 | -- | -- | L |
| *QDhm.huj.uh-4A* | 4A.3 | 4.2 | 44.3 | 36.7—51.9 | 15.2 | wsnp_Ex_c10886_17694220 | 0.02 | -0.77±0.67 | 0.10 | -1.52±0.26 | L |
| *QdfDhm.huj.uh-4A* | 4A.3 | 4.1 | 45.0 | 36.2—54.7 | 18.5 | BS00022174_51 | 0.04 | -0.63±0.35 | 0.09 | -1.29±0.32 | L |
| *QDhm.huj.uh-5A.1* | 5A.3 | 4.5 | 41.9 | 37.4—45.7 | 8.4 | wsnp_Ra_c14112_22155312 | 0.16 | -1.21±3.64 | 0.11 | -0.40±1.34 | L |
| *QDhm.huj.uh-5A.2* | 5A.8 | 5.9 | 146.2 | 141.4—151.3 | 9.9 | Ku_c19516_384 | 0.16 | 1.06±3.11 | 0.11 | 0.54±1.30 | G |
| *QDhm.huj.uh-5B* | 5B.7 | 3.8 | 120.3 | 112.2—128.3 | 16.1 | wsnp_RFL_Contig1548_762547 | 0.04 | 1.64±0.57 | 0.05 | 1.04±0.38 | G |
| *QDhm.huj.uh-6A* | 6A.1 | 3.0 | 14.1 | 7.0—27.9 | 20.9 | IAAV2577 | 0.06 | 1.84±1.13 | 0.11 | 1.50±0.38 | G |
| *QDhm.huj.uh-6B* | 6B.3 | 3.9 | 38.2 | 31.4—46.7 | 15.3 | CAP7_c6528_99 | 0.02 | -0.28±0.65 | 0.13 | -1.73±0.44 | L |
| *QDhm.huj.uh-7B* | 7B.1 | 13.6 | 14.9 | 12.9—16.4 | 3.5 | Tdurum_contig42423_1963 | 0.28 | 4.70±0.50 | -- | -- | G |
| *QdDhm.huj.uh-7B* | 7B.1 | 13.1 | 14.9 | 11.6—16.4 | 4.9 | Tdurum_contig42423_1963 | 0.32* | -4.63±0.62* | | | G |

**Table S14** Summary of QTLs. Summary of QTLs detected for 17 initial traits, 16 traits adjusted to heading and 33 drought plasticity traits in G×L RIL population under two water regimes.

| QTL | Interval, cM | LOD | # of trait effects | ITV alleles | Traits | Phenology | Drought | Drought strategy | Candidate genes |
|---|---|---|---|---|---|---|---|---|---|
| 1A.1 | 6.6—32.3 | 4.1—4.7 | single | -- | FLW | plastic | non-plastic | -- | -- |
| 1A.2 | 48.4—67.3 | 2.3—4.5 | multi | opposite | SpL, δ13C | non-plastic | non-plastic | avoidance | *TaGID1-A1, TaGA20ox-A4, GASA family* |
| 1A.3 | 81.3—110.6 | 2.3—4.7 | multi | opposite | TKW, KNSP, OP | plastic | plastic | tolerance | *GASA family, TaGA2ox-A8* |
| 1A.4 | 98.4—117.4 | 3.8 | single | -- | HI | plastic | plastic | -- | -- |
| 1B.1 | 0.0—11.4 | 3.2—3.3 | multi | uniform | DPH, FLL | linked | non-plastic | -- | -- |
| 1B.2 | 10.3—22.3 | 3.1—5.3 | single | -- | TKW | non-plastic | non-plastic | -- | -- |
| 1B.3 | 28.6—52.0 | 2.1 | single | -- | LR | non-plastic | plastic | avoidance* | *Glu-B3, TaGA2ox-B1, TaGA2ox-B10, TaGID1-B1, TaGA20ox-B4, GASA family* |
| 2A.1 | 32.7—44.9 | 3.1—4.4 | multi | uniform | GY, KNSP, SpDM, VegDM, TotDM | plastic | non-plastic | -- | *Ppd-A1* |
| 2A.2 | 34.6—64.2 | 2.9—3.8 | multi | uniform | Chl, FLW | plastic | non-plastic | stay green | *TaGA13ox-A2* |
| 2A.3 | 51.4—70.9 | 3.9—4.0 | single | -- | OP | non-plastic | non-plastic | tolerance | *FPF1-like, TaFT4-A1* |
| 2A.4 | 70.3—94.6 | 2.1 | single | -- | FLL | non-plastic | non-plastic | -- | *TaFT9-A1, TaGA2ox-A6* |
| 2A.5 | 126.3—140.7 | 3.3—4.6 | multi | opposite | SpL, LR | non-plastic | non-plastic | avoidance | *TaGly1/Zeo* |
| 2B.1 | 18.6—36.0 | 3.3—5.9 | single | -- | TKW | plastic | non-plastic | -- | -- |
| 2B.2 | 27.9—52.8 | 2.1—5.8 | multi | uniform | HI, GY | non-plastic | non-plastic | -- | *TaGA13ox-B2, FPF1-like, TaFT4-B1* |
| 2B.3 | 65.9—88.0 | 3.2—4.0 | multi | uniform | OP, Chl | plastic | non-plastic | stay green, tolerance | *TaFT9-B1, TaGA2ox-B6* |
| 2B.4 | 82.8—89.3 | 3.7—8.9 | multi | opposite | KNSP, TKW | non-plastic | plastic | -- | -- |
| 2B.5 | 77.7—111.0 | 4.4—9.3 | multi | uniform | HI, Chl | non-plastic | non-plastic | -- | -- |
| 2B.6 | 95.4—118.3 | 2.5—21.7 | multi | opposite | DPH, DHM, GY, KNSP, SpDM, CL, FLW, TotDM | linked | plastic | escape | *TaFT8-B1* |

| QTL | Interval, cM | LOD | # of trait effects | ITV alleles | Traits | Phenology | Drought | Drought strategy | Candidate genes |
|---|---|---|---|---|---|---|---|---|---|
| 2B.7 | 110.4—132.5 | 2.1—2.3 | multi | uniform | TKW, FLW | non-plastic | plastic | -- | *TaFT8-B2, TaFT8-B3, TaGA1ox-B1, TaGA3ox-B3* |
| 3A.1 | 0.0—18.3 | 2.2—5.2 | multi | opposite | FLW, δ13C | non-plastic | plastic | avoidance* | -- |
| 3A.2 | 15.4—23.4 | 5.6 | single | -- | TKW | linked | non-plastic | -- | *TaGID2-A1* |
| 3A.3 | 38.7—49.0 | 7.5 | single | -- | DPH | linked | non-plastic | -- | *TaGI-A, TaGA3ox, TaGA2ox, TaFT2-A1, FPF1-like, TaGA2ox-A3* |
| 3A.4 | 46.0—71.2 | 2.4—4.4 | multi | opposite | Chl, Cl | plastic | non-plastic | stay green | -- |
| 3A.5 | 64.9—78.0 | 2.9—6.6 | multi | uniform | FLL, δ13C | non-plastic | non-plastic | avoidance | -- |
| 3A.6 | 127.2—135.4 | 4.2 | single | -- | KNSP | plastic | non-plastic | -- | -- |
| 3B.1 | 13.7—40.1 | 3.5—3.9 | single | -- | HI | plastic | plastic | -- | -- |
| 3B.2 | 60.9—87.5 | 2.7—4.5 | multi | uniform | GY, CL, OP | non-plastic | plastic | tolerance* | -- |
| 3B.3 | 94.9—104.5 | 5.1 | single | -- | CL | linked | non-plastic | -- | -- |
| 3B.4 | 115.9—133.6 | 5.0—6.7 | single | -- | FLW | non-plastic | non-plastic | -- | -- |
| 4A.1 | 3.5—33.6 | 2.2—4.1 | multi | opposite | HI, TotDM, VegDM | plastic | plastic | -- | -- |
| 4A.2 | 22.1—74.6 | 2.0—2.1 | single | -- | Chl | plastic | plastic | stay green* | *TaGA2ox-A11* |
| 4A.3 | 23.2—54.7 | 2.6—9.5 | multi | uniform | CL, DHM, TKW, SpL, δ13C | plastic | plastic | avoidance* | *GID1, TaPRR73-A, TaGA13ox-A1* |
| 4A.4 | 35.9—74.6 | 3.2—6.6 | single | -- | TKW | plastic | plastic | -- | *GASA_family, TaFT11-A1* |
| 4A.5 | 62.7—78.9 | 2.8 | single | -- | FLW | plastic | non-plastic | -- | *TaGA20ox-A1* |
| 4A.6 | 79.1—119.5 | 2.4—4.1 | multi | opposite | GY, HI, SpL | plastic | non-plastic | -- | *Wx-B1* |
| 4A.7 | 108.7—125.6 | 3.0—6.2 | single | -- | FLW | non-plastic | plastic | -- | -- |
| 4B.1 | 0.0—21.4 | 2.0—2.9 | multi | opposite | SpDM, Chl | plastic | plastic | stay green | -- |
| 4B.2 | 8.2—23.3 | 7.0 | single | -- | Chl | plastic | non-plastic | stay green | -- |
| 4B.3 | 21.9—52.6 | 2.3—6.1 | multi | uniform | GY, TKW, SpDM, VegDM, TotDM | non-plastic | plastic | -- | *TaGA13ox-B1, TaGA2ox-B11, GID1* |
| 4B.4 | 37.1—51.0 | 2.4—7.2 | multi | opposite | DPH, FLL, CL | linked | non-plastic | -- | -- |
| 4B.5 | 50.3—75.7 | 3.3—32.4 | multi | uniform | FLW, OP | non-plastic | plastic | tolerance | -- |
| 4B.6 | 77.9—84.6 | 4.2 | single | -- | CL | plastic | non-plastic | -- | *TaGA2ox* |
| 5A.1 | 0.0—7.9 | 4.7 | single | -- | TKW | plastic | non-plastic | -- | -- |
| 5A.2 | 11.9—59.7 | 2.0—4.4 | multi | opposite | HI, VegDM, TotDM | non-plastic | plastic | -- | -- |

| QTL | Interval, cM | LOD | # of trait effects | ITV alleles | Traits | Phenology | Drought | Drought strategy | Candidate genes |
|---|---|---|---|---|---|---|---|---|---|
| 5A.3 | 32.3—56.1 | 2.8—11.2 | multi | opposite | DPH, DHM, GY, KNSP, HI, FLL, SpL, CL, Chl, SpDM, LR | plastic | plastic | avoidance | *TaFT7-A1, FPA* |
| 5A.4 | 63.1—73.9 | 5.6—7.0 | single | -- | δ13C | plastic | non-plastic | avoidance | *TaFT10-A, FPA* |
| 5A.5 | 88.0—120.8 | 2.3—18.8 | multi | uniform | FLW, DPH | linked | plastic | -- | *VRN-A1, GASA_family* |
| 5A.6 | 95.6—145.0 | 2.1—3.9 | multi | opposite | HI, CL, OP | plastic | plastic | tolerance | -- |
| 5A.7 | 129.1—144.0 | 2.3—3.6 | multi | opposite | KNSP, δ13C | linked | plastic | avoidance | *TaGA2ox-A8, FPF1* |
| 5A.8 | 140.3—157.0 | 3.3—7.7 | multi | opposite | DHP, DHM, FLL | linked | plastic | escape | *FPF1* |
| 5A.9 | 152.2—157.1 | 3.8 | single | -- | CL | plastic | non-plastic | -- | *TaFT5-A1* |
| 5B.1 | 37.8—46.5 | 2.0—5.2 | single | -- | TKW | non-plastic | non-plastic | -- | *GASA_family* |
| 5B.2 | 49.0—67.5 | 4.8 | single | -- | FLW | plastic | non-plastic | -- | -- |
| 5B.3 | 60.5—96.0 | 3.2—3.4 | multi | uniform | LR, SpDM | linked | non-plastic | -- | *TaFT10-B1, FPA,* |
| 5B.4 | 86.5—105.0 | 5.7 | single | -- | HI | linked | non-plastic | -- | *VRN-B1, GASA_family* |
| 5B.5 | 103.3—109.8 | 2.9 | single | -- | OP | non-plastic | non-plastic | tolerance | -- |
| 5B.6 | 96.3—135.0 | 2.9—4.7 | single | -- | Chl | non-plastic | non-plastic | stay green | -- |
| 5B.7 | 112.2—128.3 | 3.0—4.6 | multi | opposite | DPH, DHM, FLW | linked | non-plastic | -- | -- |
| 5B.8 | 126.5—137.6 | 4.4—7.4 | multi | uniform | CL, SpL, FLW | linked | non-plastic | -- | *WAP2-B* |
| 6A.1 | 0.0—29.3 | 2.0—4.5 | multi | opposite | HI, FLL, DHM | linked | plastic | escape | -- |
| 6A.2 | 37.3—57.0 | 2.3—8.8 | multi | uniform | FLW, KNSP, Chl | plastic | plastic | stay green | *TaFT6-A1* |
| 6A.3 | 44.7—63.3 | 2.4—3.5 | multi | uniform | TKW, OP | non-plastic | non-plastic | tolerance | *TaFT12-A1, TaGA2ox-A9* |
| 6A.4 | 58.4—90.2 | 2.0—6.4 | multi | opposite | FLL, SpL, Chl | plastic | plastic | stay green | -- |
| 6B.1 | 0.0—17.0 | 2.4 | single | -- | DPH | linked | non-plastic | -- | -- |
| 6B.2 | 22.1—49.9 | 2.5—4.9 | multi | opposite | CL, SpL, OP | plastic | plastic | tolerance | -- |
| 6B.3 | 31.4—49.8 | 2.3—3.9 | multi | uniform | TKW, DHM | non-plastic | non-plastic | -- | *Gpc-B1* |
| 6B.4 | 42.6—61.5 | 3.2—5.8 | multi | opposite | HI, FLW, δ13C | non-plastic | non-plastic | avoidance | *TaFT6-B1, TaFT12-B, TaGA2ox-B9* |
| 6B.5 | 93.0—105.0 | 4.7 | single | -- | Chl | plastic | non-plastic | stay green | -- |
| 7A.1 | 0.0—19.0 | 2.4—4.4 | multi | uniform | SpDM, VegDM, TotDM | plastic | non-plastic | -- | *KAO1* |
| 7A.2 | 14.6—26.5 | 4.7 | single | -- | TKW | plastic | non-plastic | -- | *Wx-A1* |
| 7A.3 | 26.2—40.5 | 2.3 | single | -- | FLL, SpDM | non-plastic | non-plastic | -- | -- |
| 7A.4 | 53.2—65.6 | 3.0—5.9 | single | -- | LR | non-plastic | plastic | avoidance | *TaSS-A1* |
| 7A.5 | 63.5—91.9 | 3.8—5.5 | multi | opposite | CL, SpL, TKW | plastic | non-plastic | -- | *GID1, TaFT4-A2, KO-A* |
| 7A.6 | 73.3—102.7 | 2.8—4.8 | multi | opposite | FLW, Chl, KNSP | plastic | non-plastic | -- | *TaGA2ox-A2* |

| QTL | Interval, cM | LOD | # of trait effects | ITV alleles | Traits | Phenology | Drought | Drought strategy | Candidate genes |
|---|---|---|---|---|---|---|---|---|---|
| 7A.7 | 94.7—111.7 | 2.9 | single | -- | CL | plastic | plastic | -- | -- |
| 7B.1 | 4.7—29.5 | 2.8—40.8 | multi | opposite | GY, HI, KNSP, TKW, DPH, DHM, SpDM, SpL, CL | linked | plastic | escape | *VRN-B3* |
| 7B.2 | 14.6—31.7 | 3.9—6.0 | single | -- | δ13C | non-plastic | plastic | avoidance | -- |
| 7B.3 | 23.9—57.7 | 2.2 | single | -- | VegDM | non-plastic | plastic | -- | *GASA_family* |
| 7B.4 | 113.2—128.4 | — | multi | opposite | KNSP, SpDM, Chl, FLW | plastic | plastic | stay green | *CPS-B* |

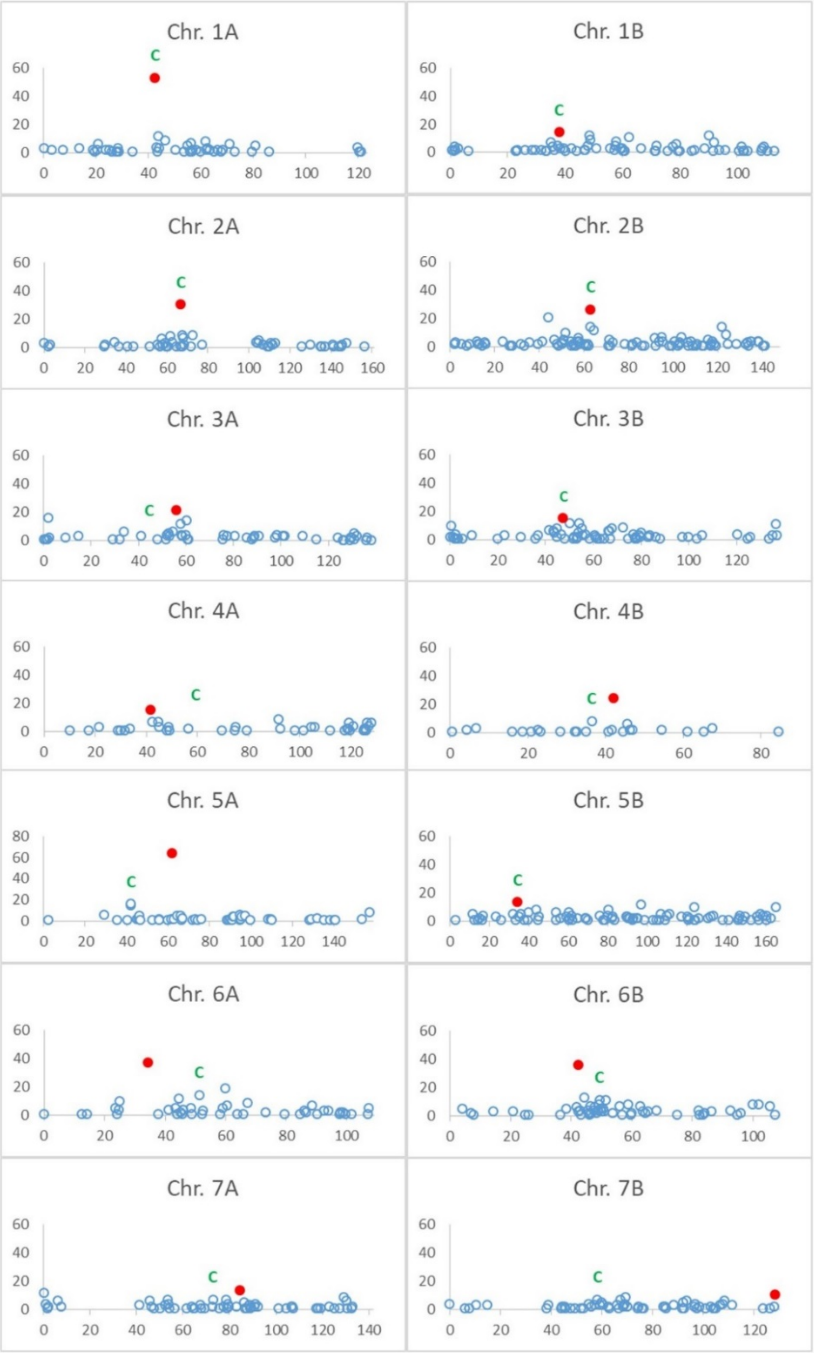


**Fig. S1** Distribution of attached markers along 14 wheat chromosomes.
Unique genetic positions on the genetic map (Skeletal markers) include attached markers that are completely linked. The distributions of attached markers are shown for each chromosome in a separated plot. The position with the highest number of attached markers in each chromosome is marked by red circle; the putative position of the centromere is marked with green “C”.

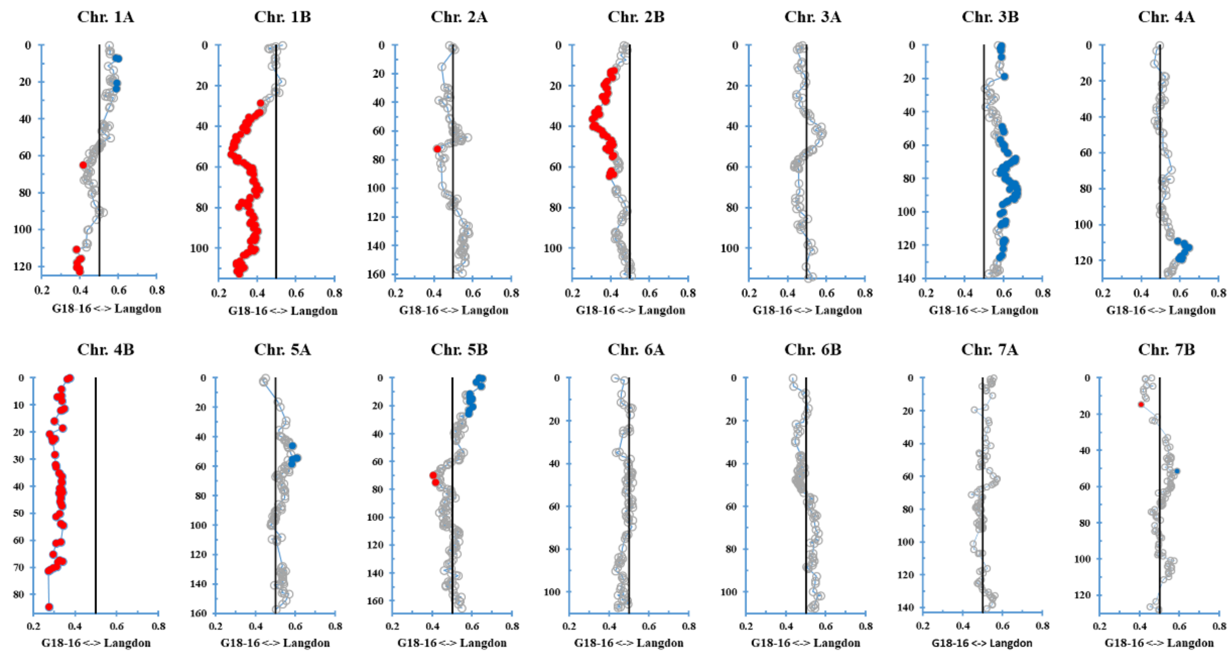


**Fig. S2** Segregation distortion for each chromosome in the G×L RIL population. Markers with significant (P<0.05) distortion are shown in color, dark blue for LDN and red for G18-16 alleles excess.

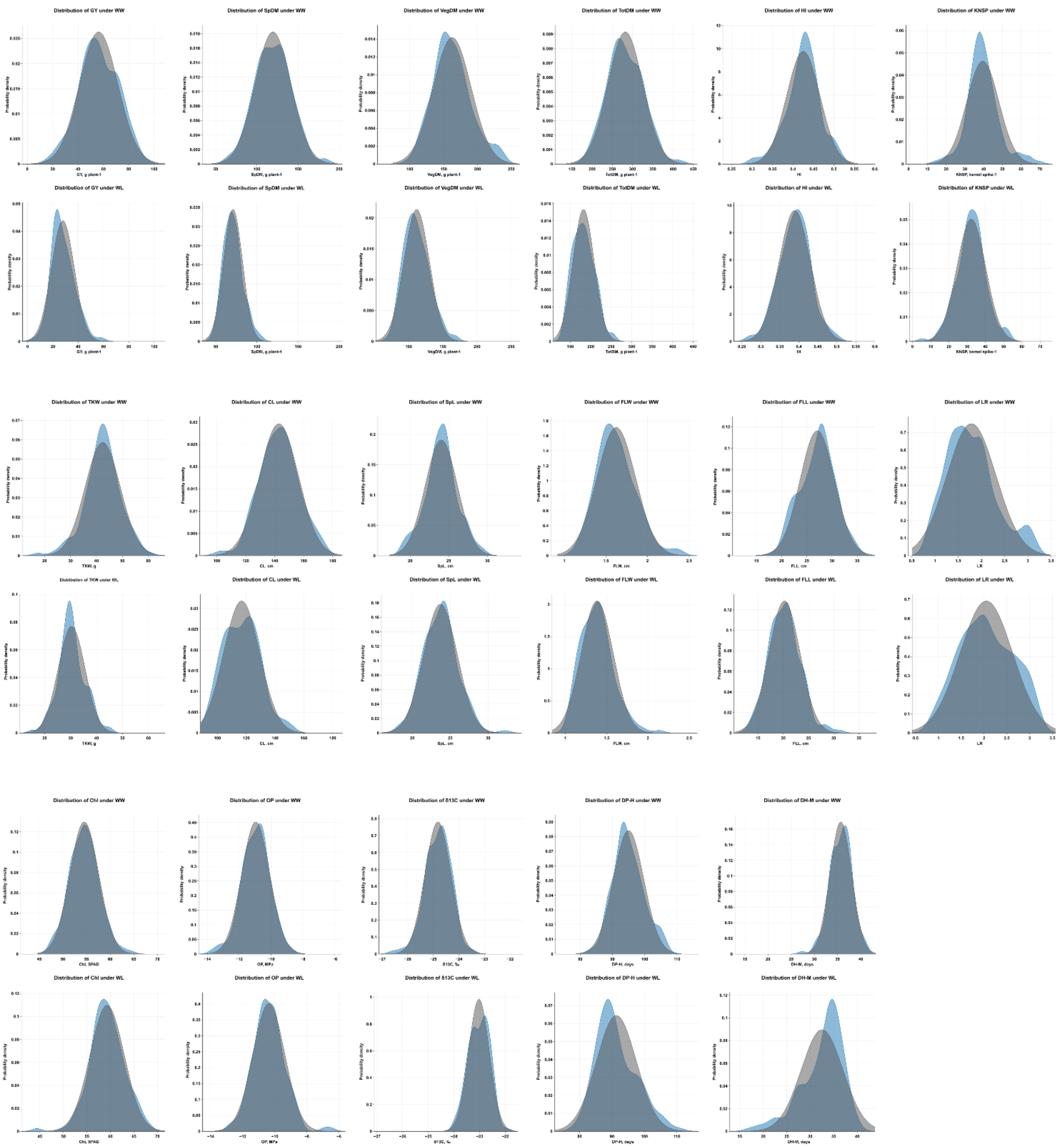


**Fig. S3** Frequency distribution of the 150 F6 RILs for 17 initial phenotypic traits. Phenotypic distribution of 17 initial traits demonstrated by frequency of the trait among 150 $F_6$ RILs. (a) Grain yield (GY), (b) thousand kernel weight (TKW), (c) kernel number per spike (KNSP), (d) harvest index (HI), (e) spike dry matter (SpDM), (f) vegetative dry matter (VegDM), (g) total dry matter (TotDM), (h) culm length (CL), (i) spike length (SpL), (j) flag leave length (FLL), (k) flag leave width (FLW), (l) carbon isotope ratio (δ13C), (m) osmotic potential (OP), (n) chlorophyll content (Chl), (o) leave rolling index (LR), (p) days from planting to heading (DP-H), (q) days from heading to maturity (DH-M) under WL and WW irrigation regimes. The expected normal distribution for each trait is presented in gray, while the observed distribution is presented in blue.

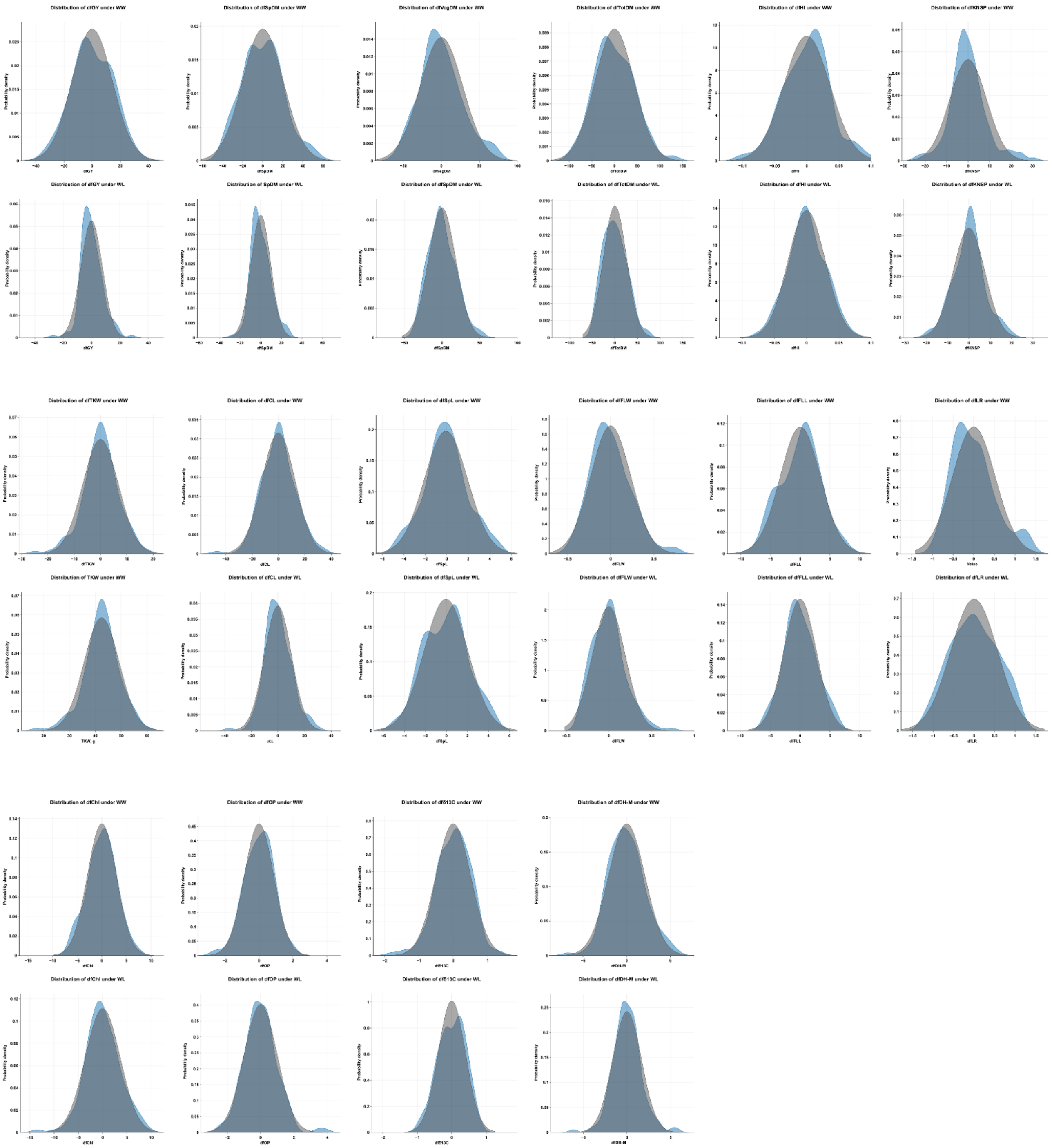

**Fig. S4** Frequency distribution of the 150 F6 RILs for 16 adjusted for heading date traits. Phenotypic distribution of 16 adjusted traits for heading demonstrated by frequency of the trait among 150 F6 RILs. (a) Grain yield (GY), (b) thousand kernel weight (TKW), (c) kernel number per spike (KNSP), (d) harvest index (HI), (e) spike dry matter (SpDM), (f) vegetative dry matter (VegDM), (g) total dry matter (TotDM), (h) culm length (CL), (i) spike length (SpL), (j) flag leave length (FLL), (k) flag leave width (FLW), (l) carbon isotope ratio (δ13C), (m) osmotic potential (OP), (n) chlorophyll content (Chl), (o) leave rolling index (LR), (p) days from planting to heading (DP-H), (q) days from heading to maturity (DH-M) under WL and WW irrigation regimes. The expected normal distribution for each trait is presented in gray, while the observed distribution is presented in blue.

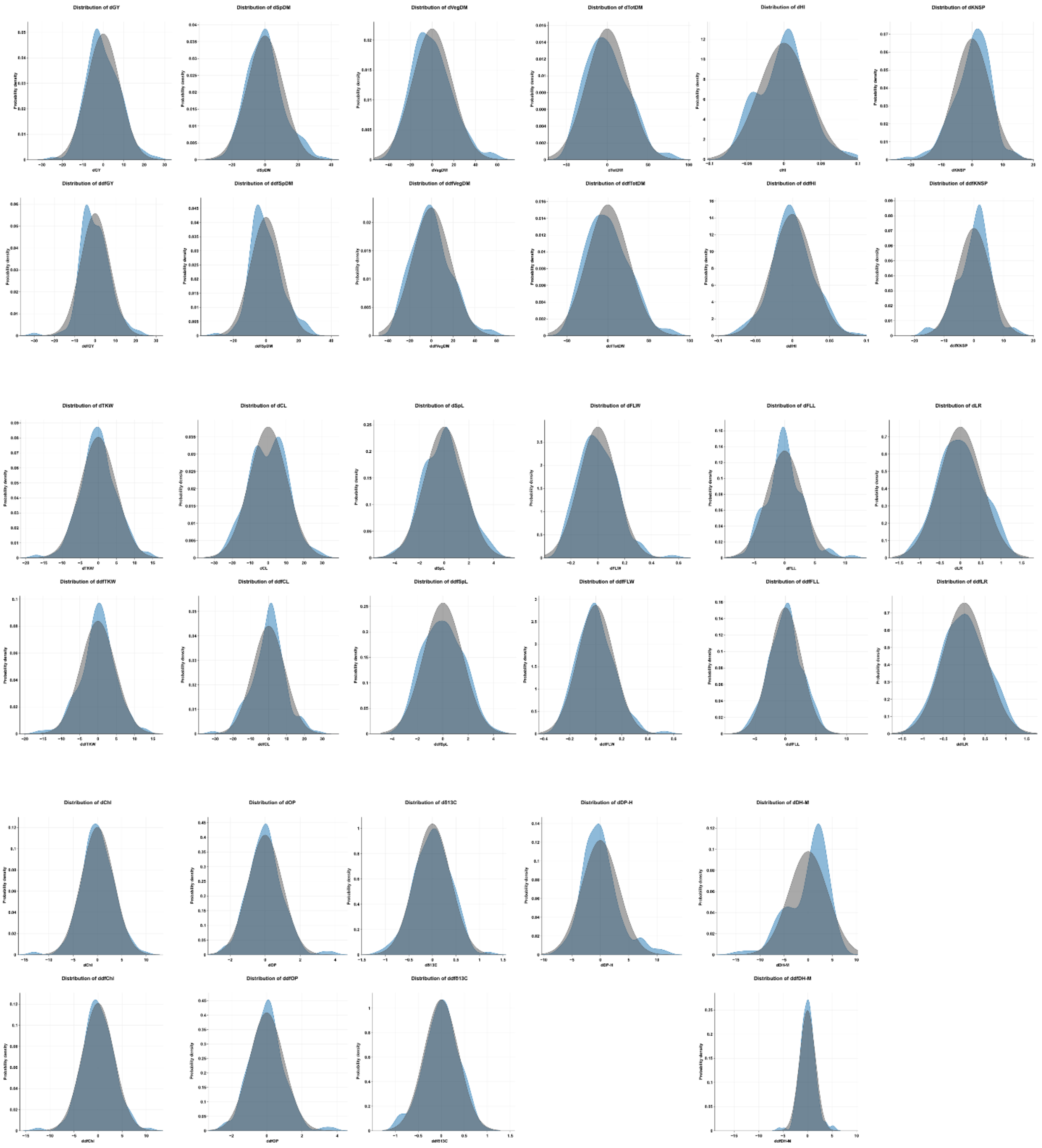


**Fig. S5** Frequency distribution of the 150 F6 RILs for 33 drought plasticity traits. Phenotypic distribution of 33 drought plasticity traits demonstrated by frequency of the trait among 150 $F_6$ RILs. (a) Grain yield (GY), (b) thousand kernel weight (TKW), (c) kernel number per spike (KNSP), (d) harvest index (HI), (e) spike dry matter (SpDM), (f) vegetative dry matter (VegDM), (g) total dry matter (TotDM), (h) culm length (CL), (i) spike length (SpL), (j) flag leave length (FLL), (k) flag leave width (FLW), (l) carbon isotope ratio (δ13C), (m) osmotic potential (OP), (n) chlorophyll content (Chl), (o) leave rolling index (LR), (p) days from planting to heading (DP-H), (q) days from heading to maturity (DH-M) under WL and WW irrigation regimes. The expected normal distribution for each trait is presented in gray, while the observed distribution is presented in blue.

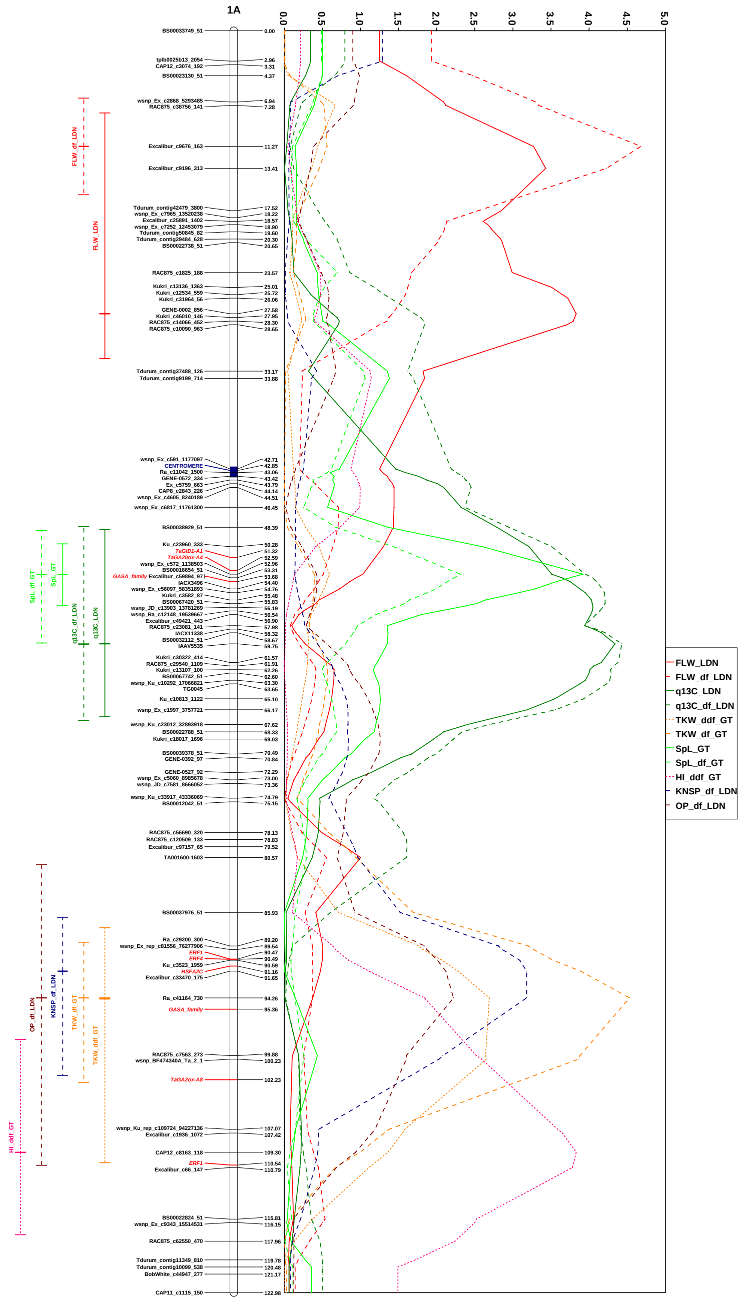


**Fig. S6** Genetic map, putative genetic positions of target genes, likelihood plots and 1.5 LOD support intervals of QTL effects for chromosome 1A.

Genetic map based on skeletal markers, putative genetic positions of target genes were marked in red. Likelihood plots and 1.5 LOD support intervals were named according to corresponding traits, where first part is name of trait, where name of trait without suffix is name of initial trait (solid line), suffix “df” is marked traits accounting to heading (dash line), suffix “d” is marked plasticity traits trait without accounting to heading (dash line with dots) and suffix “ddf” is marked plasticity traits trait with accounting to heading (dotted line). Last part of name includes association with increase trait value allele: G18-16 allele is named with suffix “GT” and Langdon allele is named with suffix “LDN”.

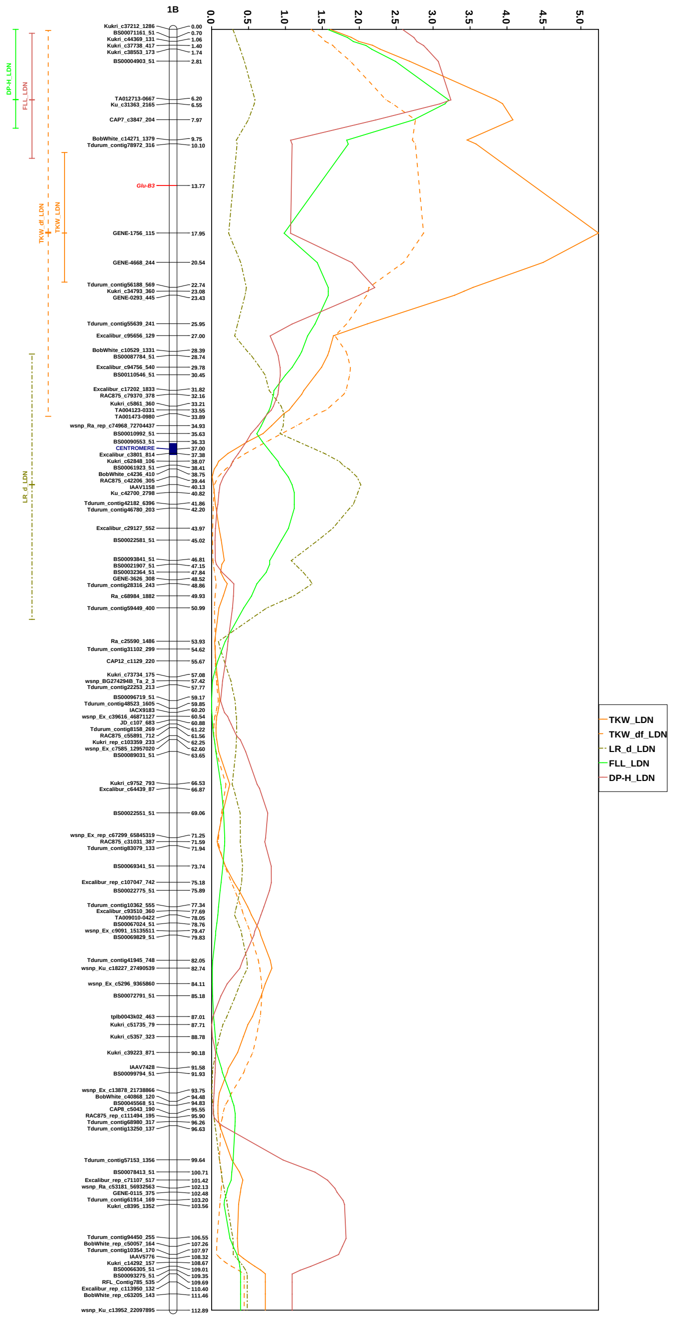


**Fig. S7** Genetic map, putative genetic positions of target genes, likelihood plots and 1.5 LOD support intervals of QTL effects for chromosome 1B.

Genetic map based on skeletal markers, putative genetic positions of target genes were marked in red. Likelihood plots and 1.5 LOD support intervals were named according to corresponding traits, where first part is name of trait, where name of trait without suffix is name of initial trait (solid line), suffix “df” is marked traits accounting to heading (dash line), suffix “d” is marked plasticity traits trait without accounting to heading (dash line with dots) and suffix “ddf” is marked plasticity traits trait with accounting to heading (dotted line). Last part of name includes association with increase trait value allele: G18-16 allele is named with suffix “GT” and Langdon allele is named with suffix “LDN”.

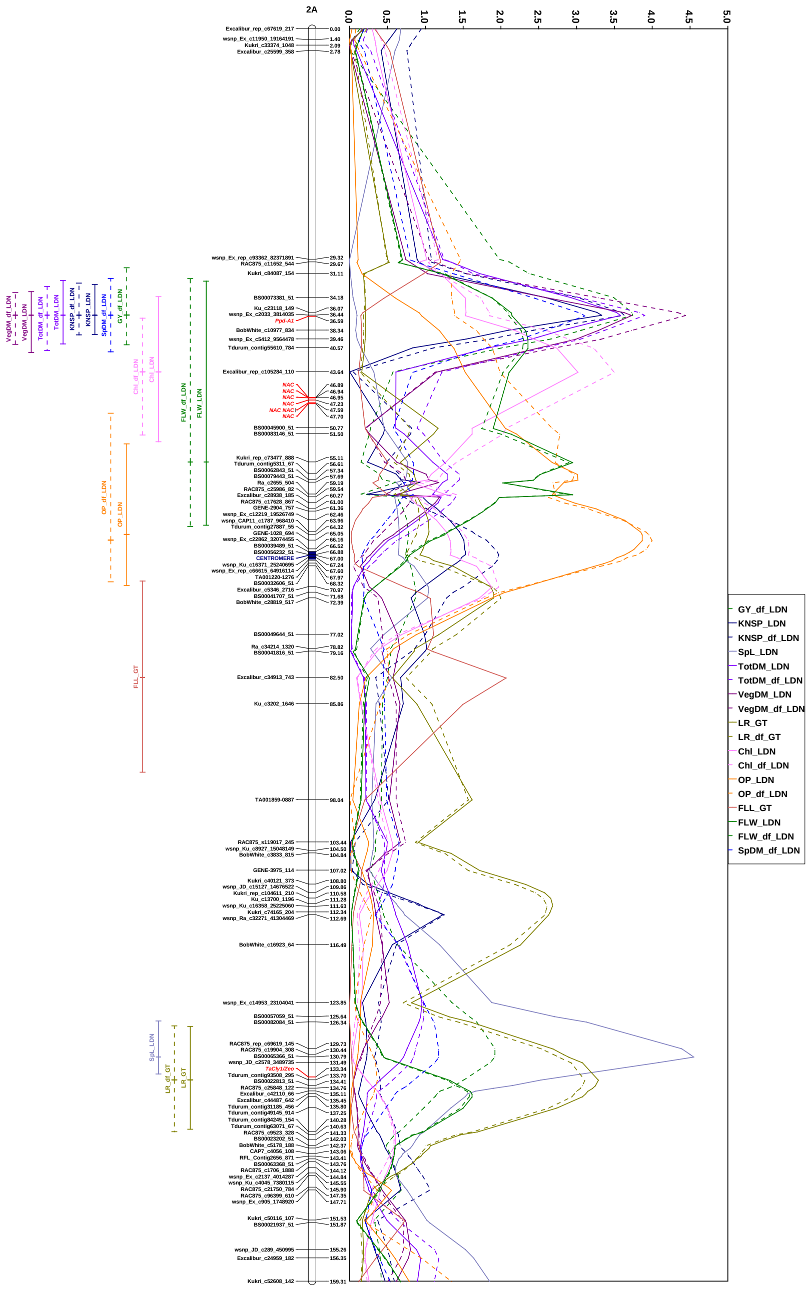


**Fig. S8** Genetic map, putative genetic positions of target genes, likelihood plots and 1.5 LOD support intervals of QTL effects for chromosome 2A.

Genetic map based on skeletal markers, putative genetic positions of target genes were marked in red. Likelihood plots and 1.5 LOD support intervals were named according to corresponding traits, where first part is name of trait, where name of trait without suffix is name of initial trait (solid line), suffix "df" is marked traits accounting to heading (dash line), suffix "d" is marked plasticity traits trait without accounting to heading (dash line with dots) and suffix "ddf" is marked plasticity traits trait with accounting to heading (dotted line). Last part of name includes association with increase trait value allele: G18-16 allele is named with suffix "GT" and Langdon allele is named with suffix "LDN".

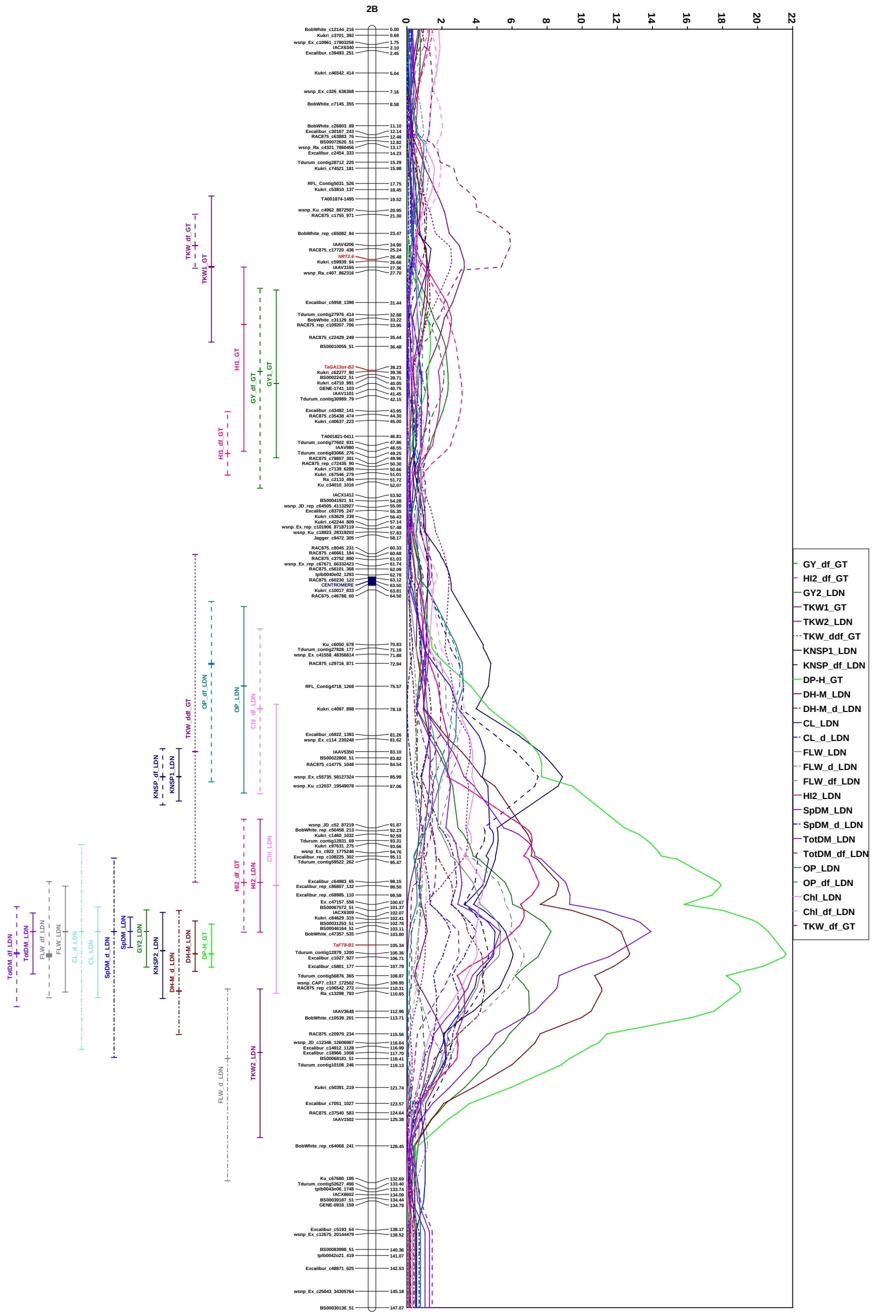

**Fig. S9** Genetic map, putative genetic positions of target genes, likelihood plots and 1.5 LOD support intervals of QTL effects for chromosome 2B.

Genetic map based on skeletal markers, putative genetic positions of target genes were marked in red. Likelihood plots and 1.5 LOD support intervals were named according to corresponding traits, where first part is name of trait, where name of trait without suffix is name of initial trait (solid line), suffix "df" is marked traits accounting to heading (dash line), suffix "d" is marked plasticity traits trait without accounting to heading (dash line with dots) and suffix "ddf" is marked plasticity traits trait with accounting to heading (dotted line). Last part of name includes association with increase trait value allele: G18-16 allele is named with suffix "GT" and Langdon allele is named with suffix "LDN".

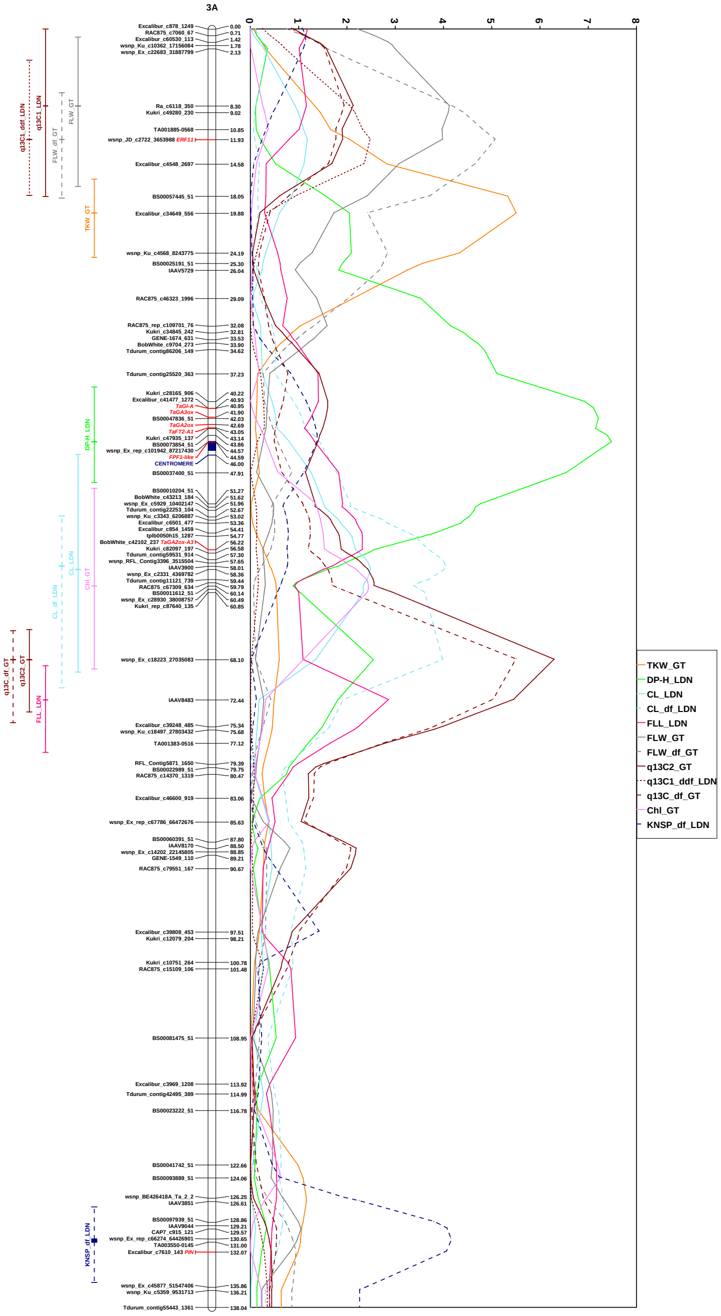


**Fig. S10** Genetic map, putative genetic positions of target genes, likelihood plots and 1.5 LOD support intervals of QTL effects for chromosome 3A.

Genetic map based on skeletal markers, putative genetic positions of target genes were marked in red. Likelihood plots and 1.5 LOD support intervals were named according to corresponding traits, where first part is name of trait, where name of trait without suffix is name of initial trait (solid line), suffix "df" is marked traits accounting to heading (dash line), suffix "d" is marked plasticity traits trait without accounting to heading (dash line with dots) and suffix "ddf" is marked plasticity traits trait with accounting to heading (dotted line). Last part of name includes association with increase trait value allele: G18-16 allele is named with suffix "GT" and Langdon allele is named with suffix "LDN".

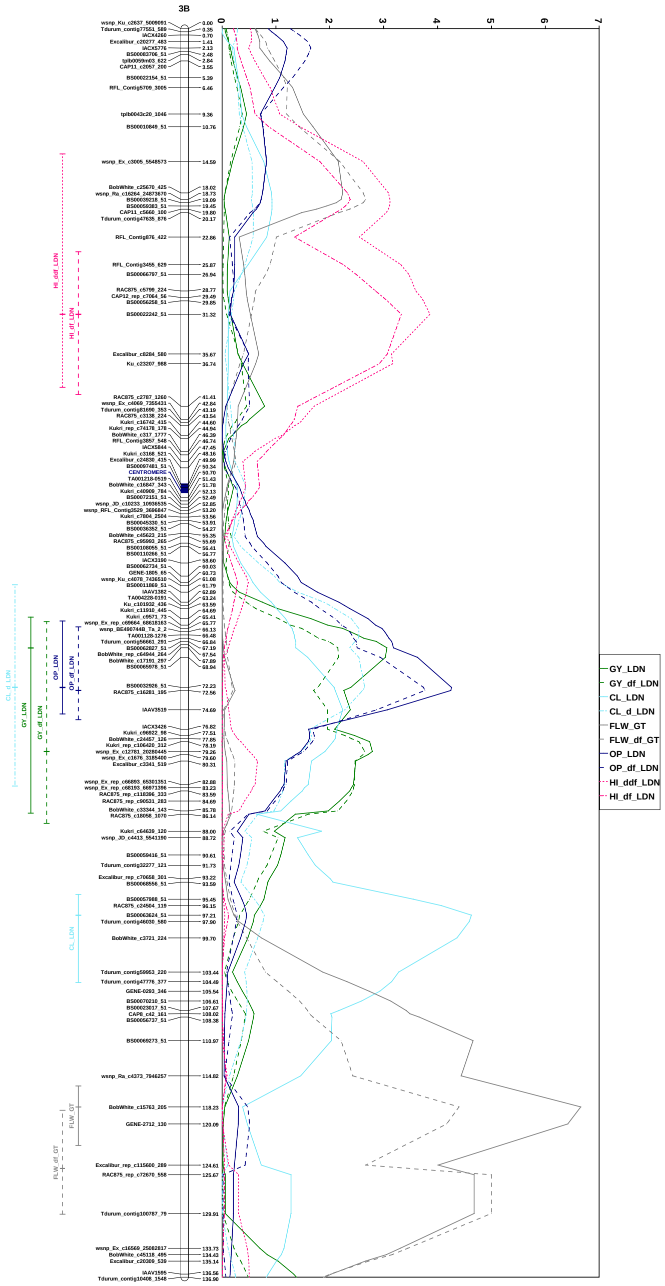


**Fig. S11** Genetic map, putative genetic positions of target genes, likelihood plots and 1.5 LOD support intervals of QTL effects for chromosome 3B.

Genetic map based on skeletal markers, putative genetic positions of target genes were marked in red. Likelihood plots and 1.5 LOD support intervals were named according to corresponding traits, where first part is name of trait, where name of trait without suffix is name of initial trait (solid line), suffix "df" is marked traits accounting to heading (dash line), suffix "d" is marked plasticity traits trait without accounting to heading (dash line with dots) and suffix "ddf" is marked plasticity traits trait with accounting to heading (dotted line). Last part of name includes association with increase trait value allele: G18-16 allele is named with suffix "GT" and Langdon allele is named with suffix "LDN".

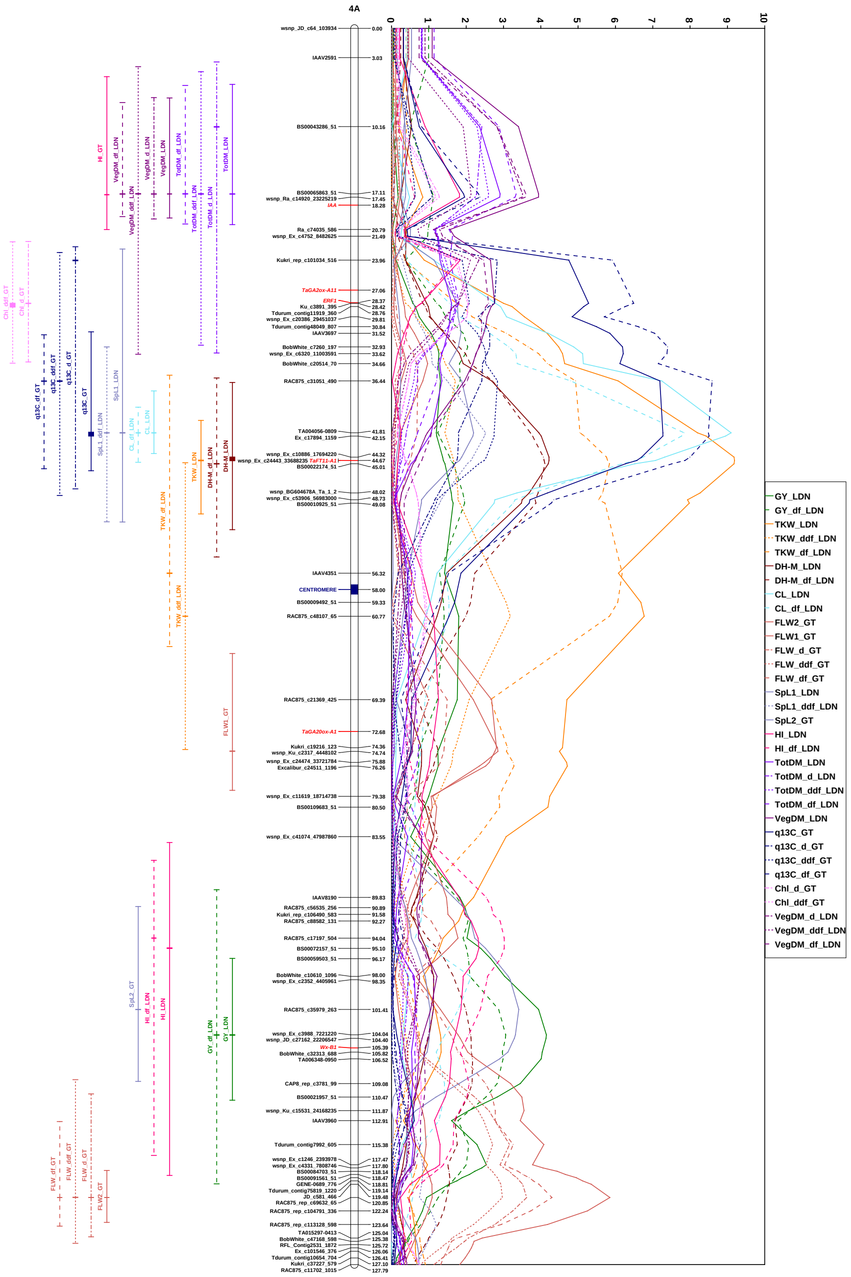


**Fig. S12** Genetic map, putative genetic positions of target genes, likelihood plots and 1.5 LOD support intervals of QTL effects for chromosome 4A.

Genetic map based on skeletal markers, putative genetic positions of target genes were marked in red. Likelihood plots and 1.5 LOD support intervals were named according to corresponding traits, where first part is name of trait, where name of trait without suffix is name of initial trait (solid line), suffix "df" is marked traits accounting to heading (dash line), suffix "d" is marked plasticity traits trait without accounting to heading (dash line with dots) and suffix "ddf" is marked plasticity traits trait with accounting to heading (dotted line). Last part of name includes association with increase trait value allele: G18-16 allele is named with suffix "GT" and Langdon allele is named with suffix "LDN".

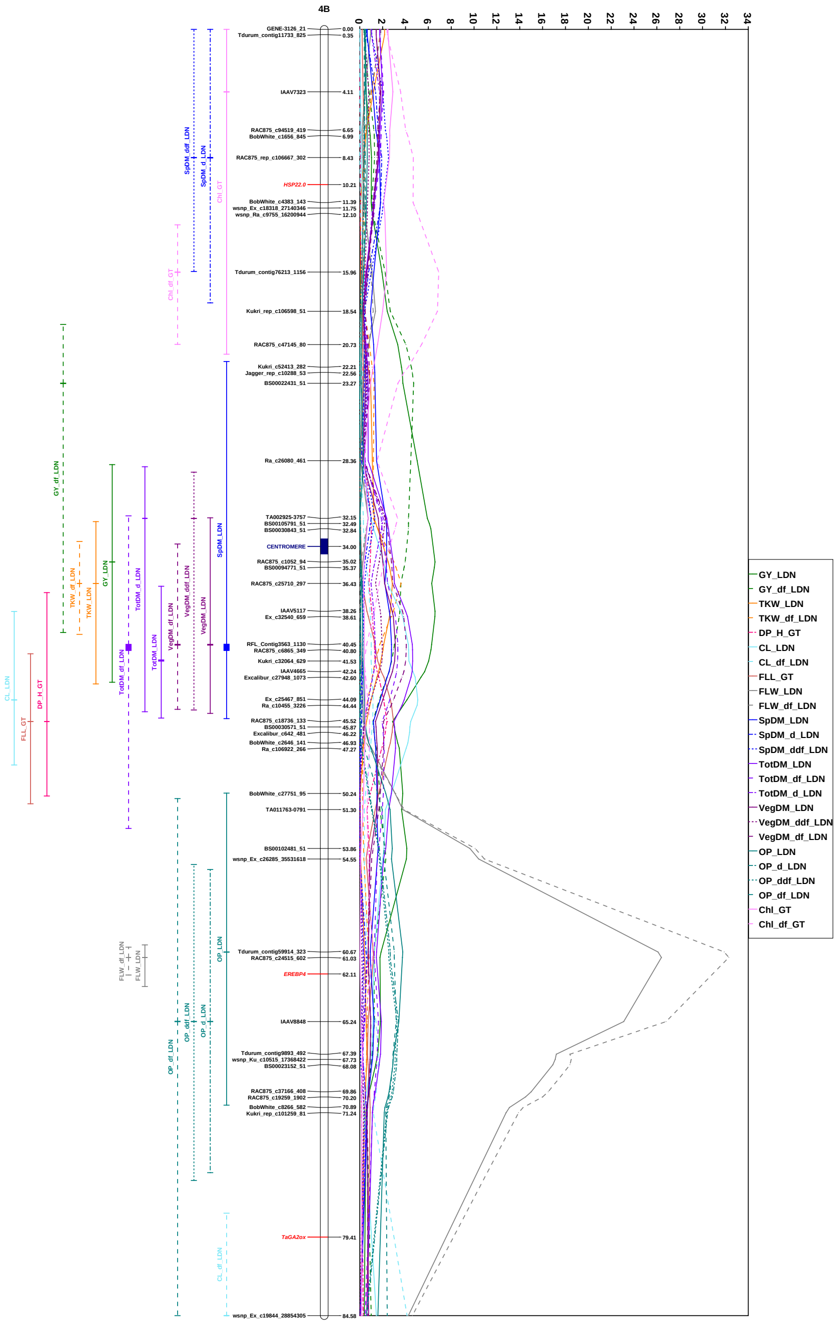


**Fig. S13** Genetic map, putative genetic positions of target genes, likelihood plots and 1.5 LOD support intervals of QTL effects for chromosome 4B.

Genetic map based on skeletal markers, putative genetic positions of target genes were marked in red. Likelihood plots and 1.5 LOD support intervals were named according to corresponding traits, where first part is name of trait, where name of trait without suffix is name of initial trait (solid line), suffix "df" is marked traits accounting to heading (dash line), suffix "d" is marked plasticity traits trait without accounting to heading (dash line with dots) and suffix "ddf" is marked plasticity traits trait with accounting to heading (dotted line). Last part of name includes association with increase trait value allele: G18-16 allele is named with suffix "GT" and Langdon allele is named with suffix "LDN".

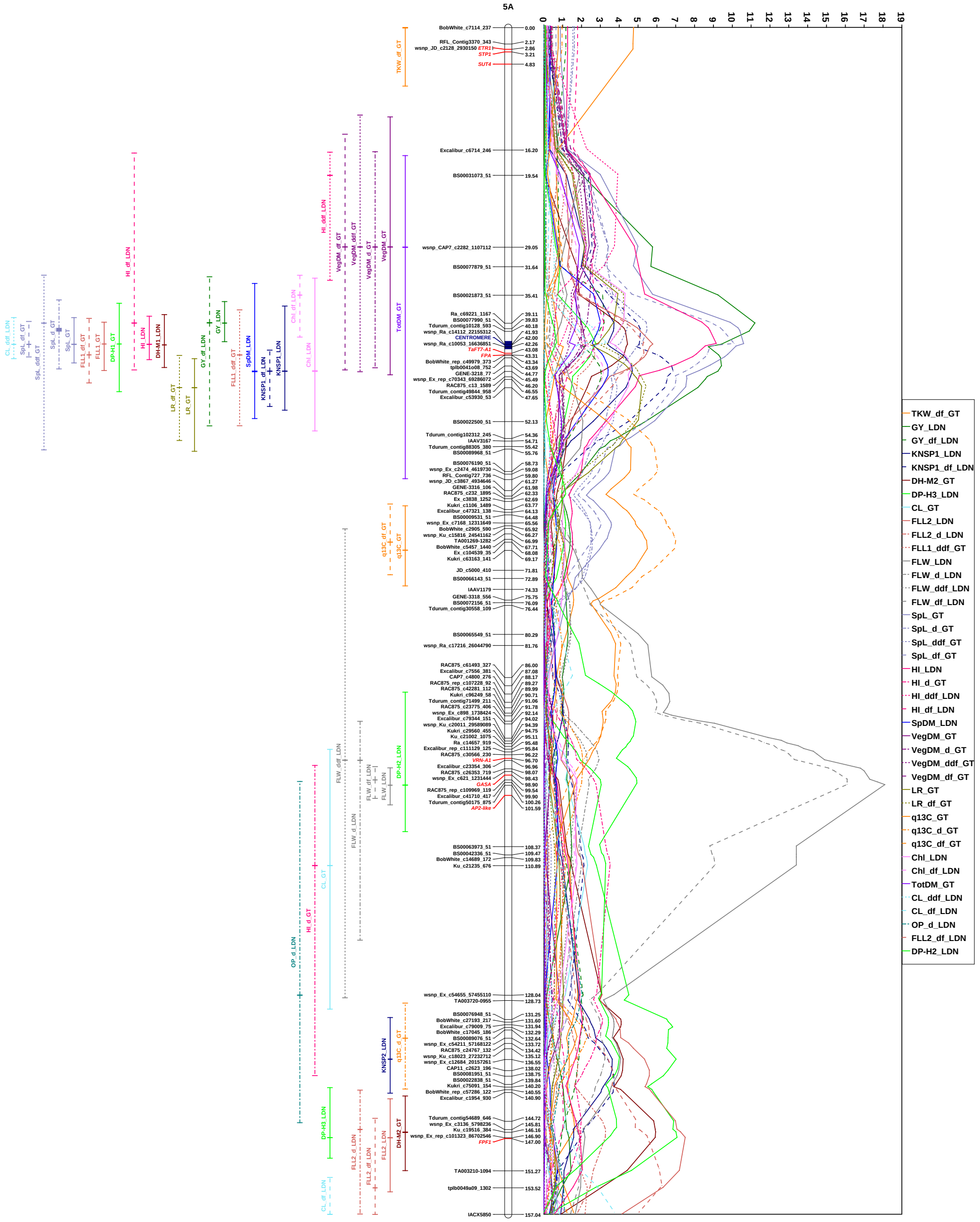


**Fig. S14** Genetic map, putative genetic positions of target genes, likelihood plots and 1.5 LOD support intervals of QTL effects for chromosome 5A.

Genetic map based on skeletal markers, putative genetic positions of target genes were marked in red. Likelihood plots and 1.5 LOD support intervals were named according to corresponding traits, where first part is name of trait, where name of trait without suffix is name of initial trait (solid line), suffix “df” is marked traits accounting to heading (dash line), suffix “d” is marked plasticity traits trait without accounting to heading (dash line with dots) and suffix “ddf” is marked plasticity traits trait with accounting to heading (dotted line). Last part of name includes association with increase trait value allele: G18-16 allele is named with suffix “GT” and Langdon allele is named with suffix “LDN”.

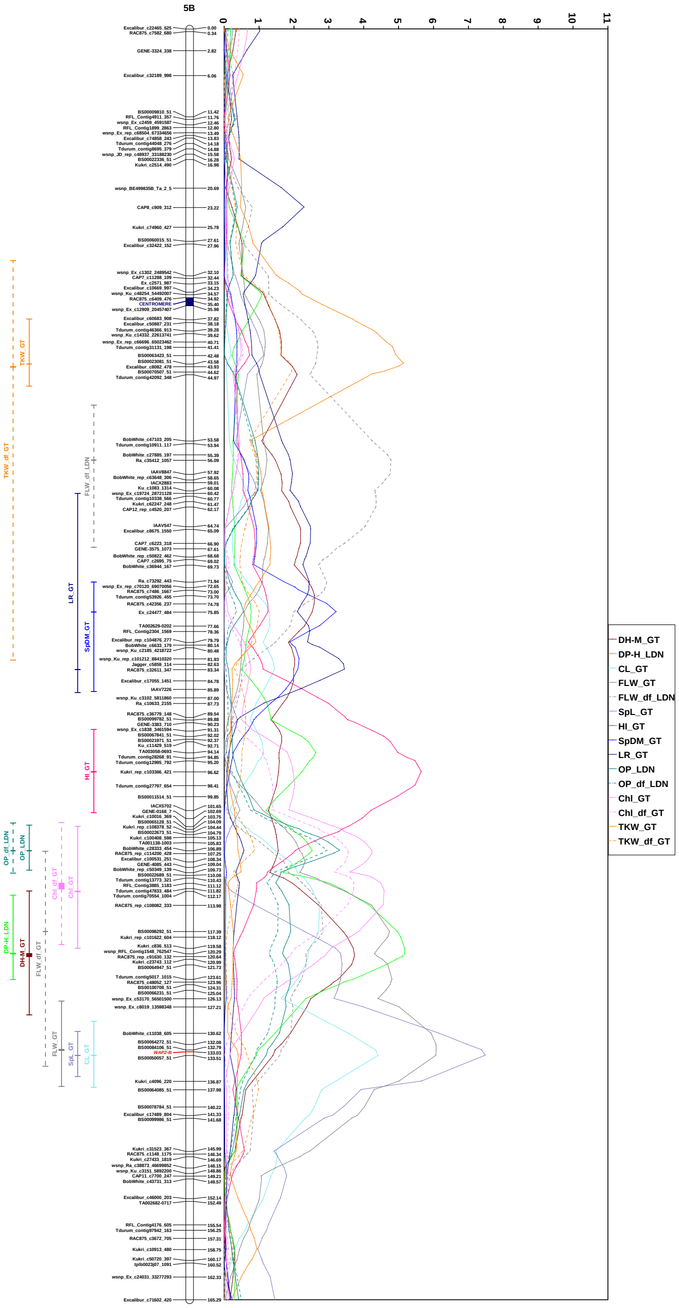


**Fig. S15** Genetic map, putative genetic positions of target genes, likelihood plots and 1.5 LOD support intervals of QTL effects for chromosome 5B.

Genetic map based on skeletal markers, putative genetic positions of target genes were marked in red. Likelihood plots and 1.5 LOD support intervals were named according to corresponding traits, where first part is name of trait, where name of trait without suffix is name of initial trait (solid line), suffix “df” is marked traits accounting to heading (dash line), suffix “d” is marked plasticity traits trait without accounting to heading (dash line with dots) and suffix “ddf” is marked plasticity traits trait with accounting to heading (dotted line). Last part of name includes association with increase trait value allele: G18-16 allele is named with suffix “GT” and Langdon allele is named with suffix “LDN”.

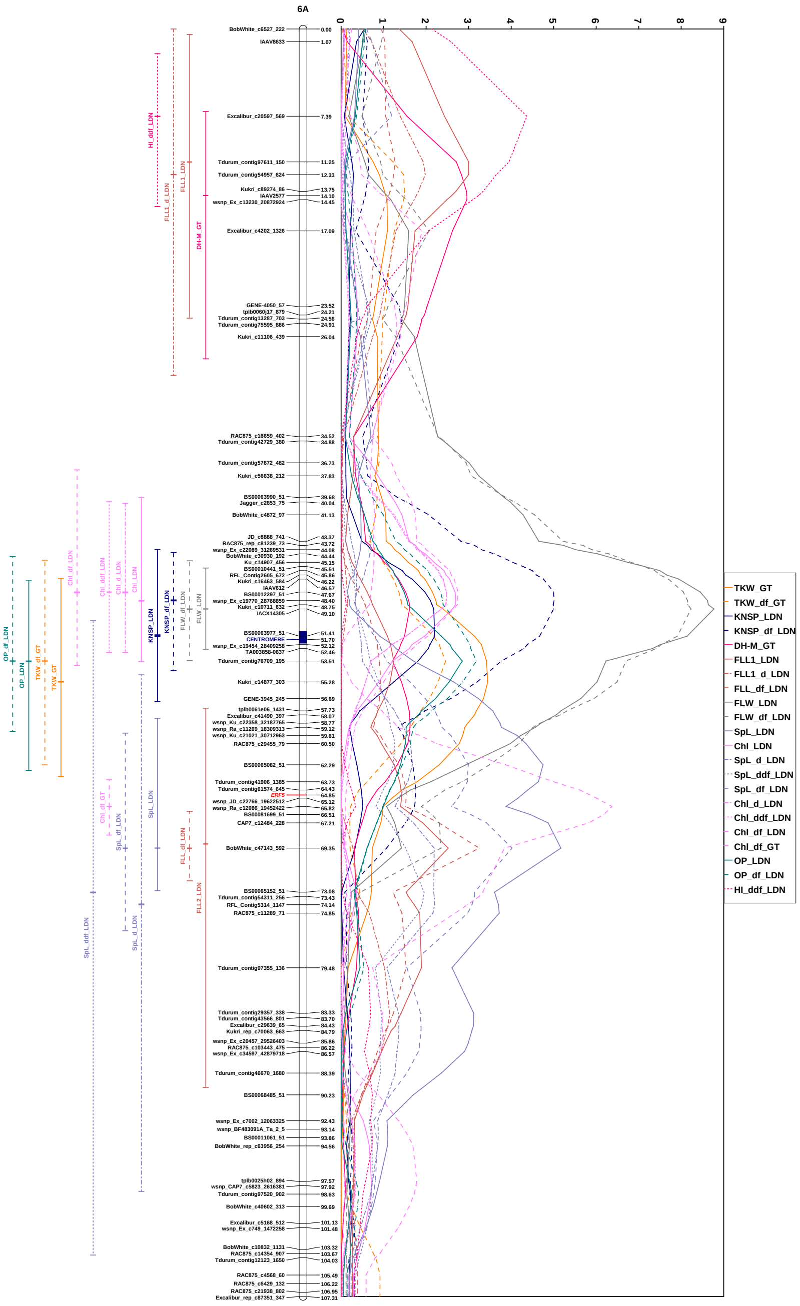


**Fig. S16** Genetic map, putative genetic positions of target genes, likelihood plots and 1.5 LOD support intervals of QTL effects for chromosome 6A.

Genetic map based on skeletal markers, putative genetic positions of target genes were marked in red. Likelihood plots and 1.5 LOD support intervals were named according to corresponding traits, where first part is name of trait, where name of trait without suffix is name of initial trait (solid line), suffix "df" is marked traits accounting to heading (dash line), suffix "d" is marked plasticity traits trait without accounting to heading (dash line with dots) and suffix "ddf" is marked plasticity traits trait with accounting to heading (dotted line). Last part of name includes association with increase trait value allele: G18-16 allele is named with suffix "GT" and Langdon allele is named with suffix "LDN".

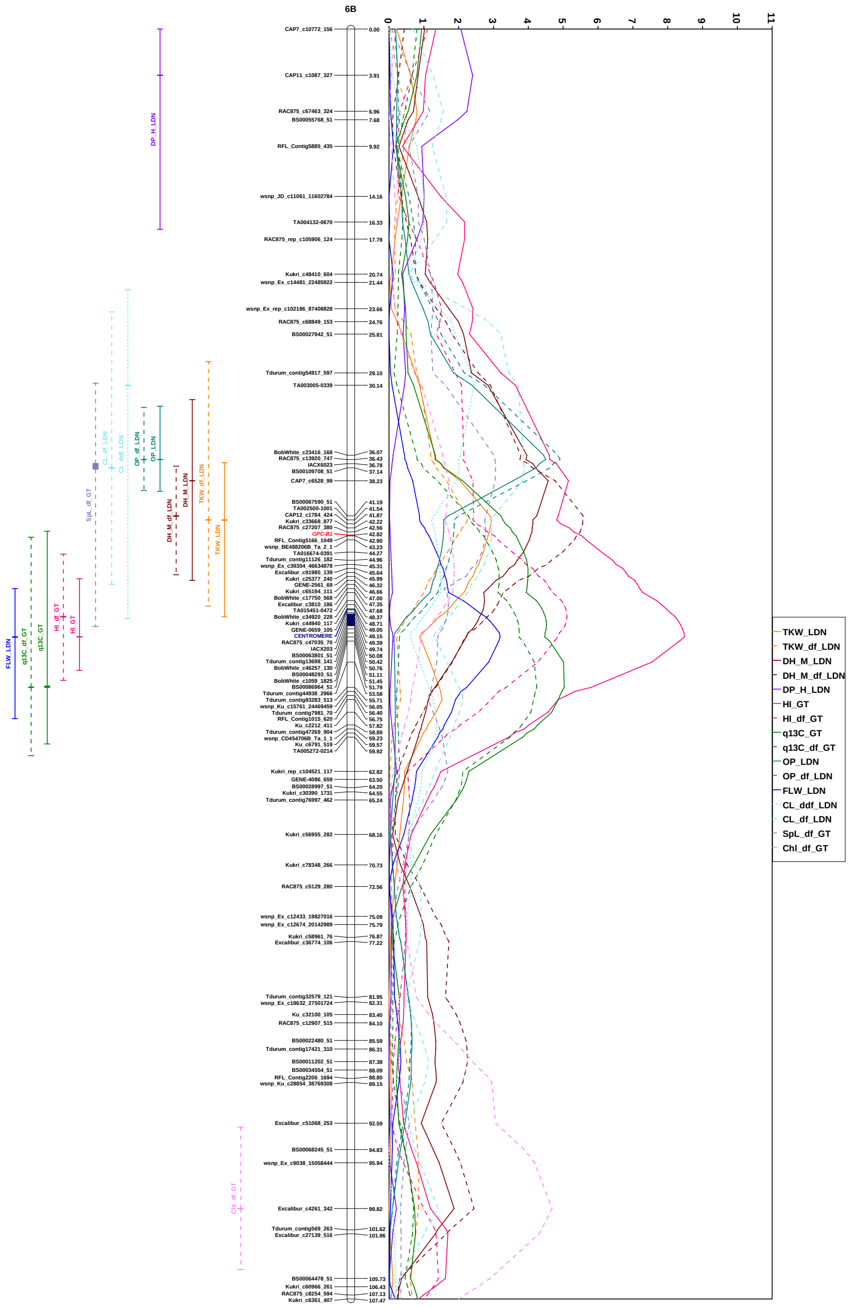


**Fig. S17** Genetic map, putative genetic positions of target genes, likelihood plots and 1.5 LOD support intervals of QTL effects for chromosome 6B.

Genetic map based on skeletal markers, putative genetic positions of target genes were marked in red. Likelihood plots and 1.5 LOD support intervals were named according to corresponding traits, where first part is name of trait, where name of trait without suffix is name of initial trait (solid line), suffix “df” is marked traits accounting to heading (dash line), suffix “d” is marked plasticity traits trait without accounting to heading (dash line with dots) and suffix “ddf” is marked plasticity traits trait with accounting to heading (dotted line). Last part of name includes association with increase trait value allele: G18-16 allele is named with suffix “GT” and Langdon allele is named with suffix “LDN”.

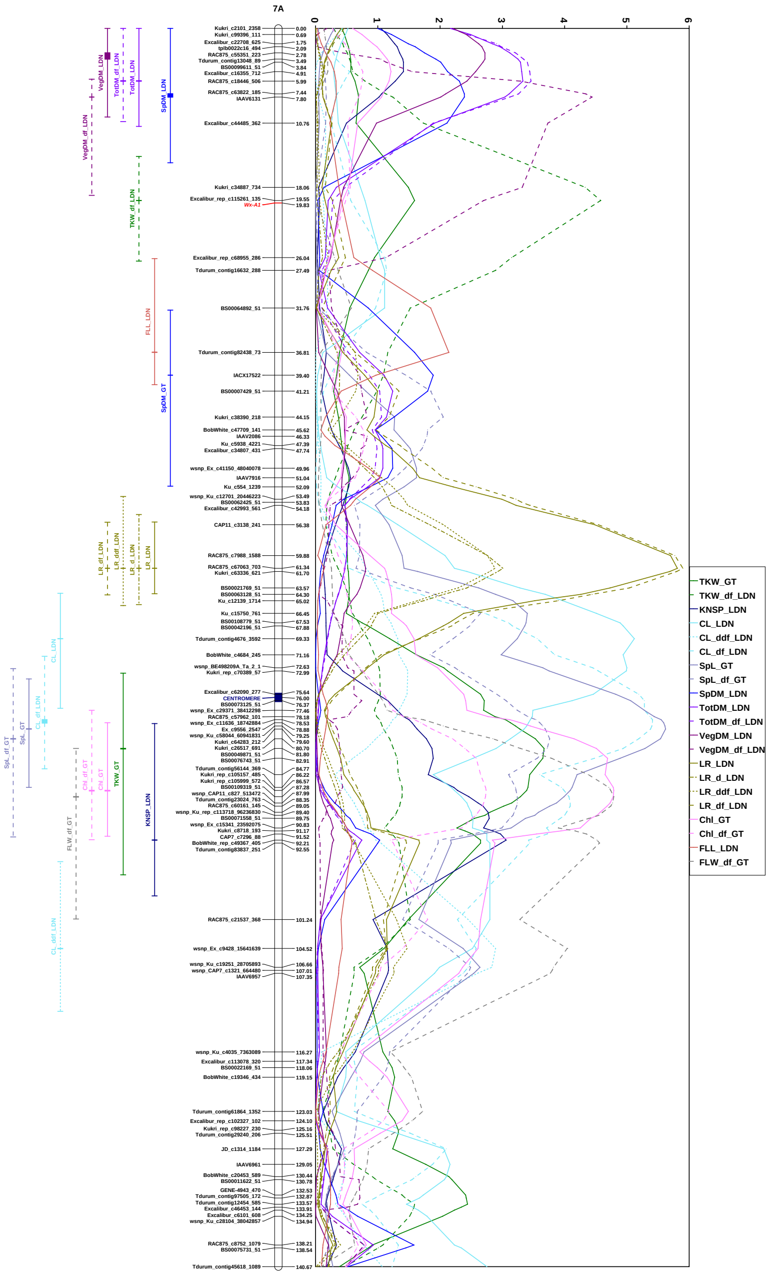


**Fig. S18** Genetic map, putative genetic positions of target genes, likelihood plots and 1.5 LOD support intervals of QTL effects for chromosome 7A.

Genetic map based on skeletal markers, putative genetic positions of target genes were marked in red. Likelihood plots and 1.5 LOD support intervals were named according to corresponding traits, where first part is name of trait, where name of trait without suffix is name of initial trait (solid line), suffix "df" is marked traits accounting to heading (dash line), suffix "d" is marked plasticity traits trait without accounting to heading (dash line with dots) and suffix "ddf" is marked plasticity traits trait with accounting to heading (dotted line). Last part of name includes association with increase trait value allele: G18-16 allele is named with suffix "GT" and Langdon allele is named with suffix "LDN".

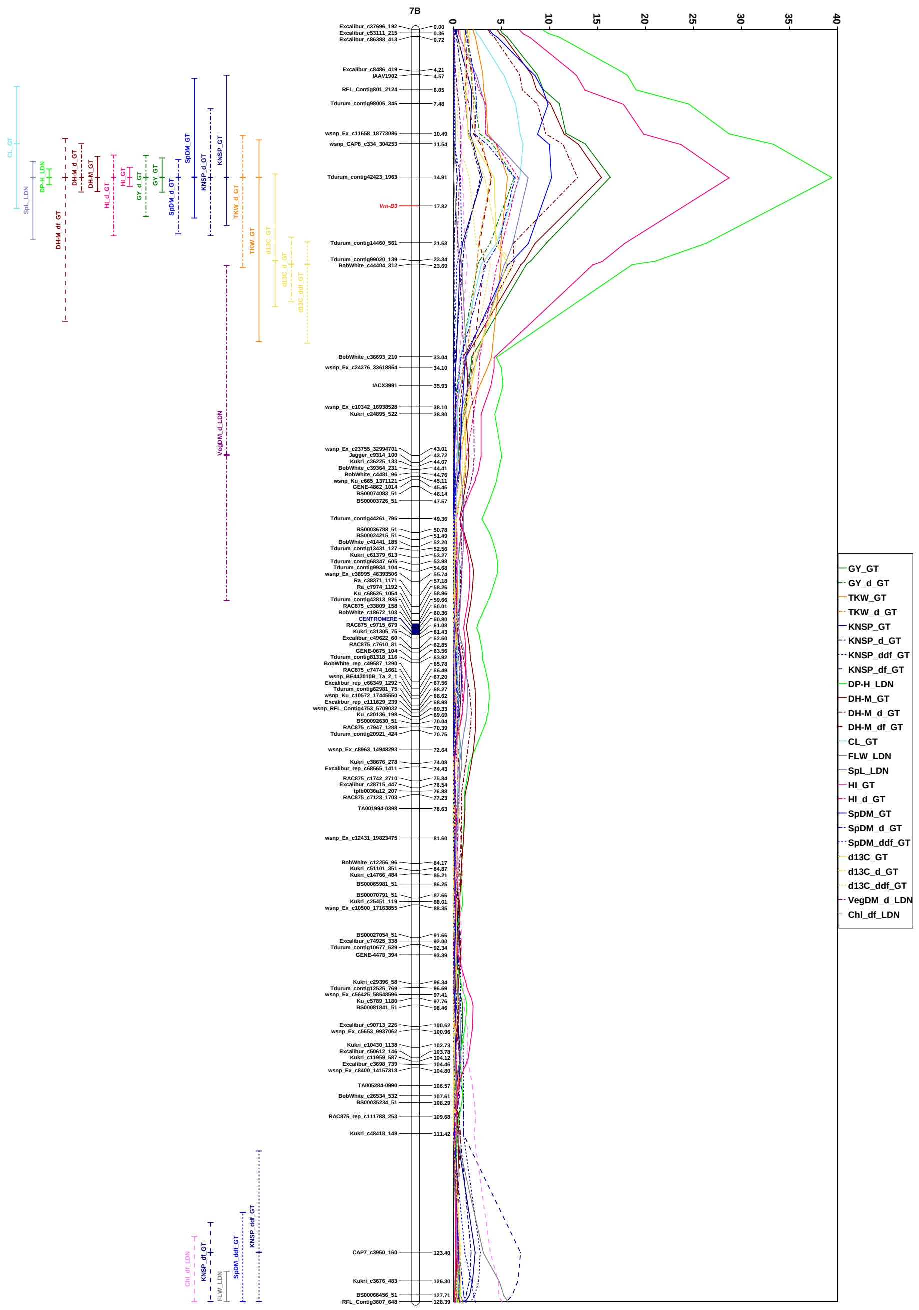


**Fig. S19** Genetic map, putative genetic positions of target genes, likelihood plots and 1.5 LOD support intervals of QTL effects for chromosome 7B.

Genetic map based on skeletal markers, putative genetic positions of target genes were marked in red. Likelihood plots and 1.5 LOD support intervals were named according to corresponding traits, where first part is name of trait, where name of trait without suffix is name of initial trait (solid line), suffix “df” is marked traits accounting to heading (dash line), suffix “d” is marked plasticity traits trait without accounting to heading (dash line with dots) and suffix “ddf” is marked plasticity traits trait with accounting to heading (dotted line). Last part of name includes association with increase trait value allele: G18-16 allele is named with suffix “GT” and Langdon allele is named with suffix “LDN”.
